# Regioselective Rapid Ene-type Reaction (RRER) Enables Bioconjugation of Histone Serotonylation

**DOI:** 10.1101/2025.04.22.650073

**Authors:** Jinghua Wu, Huapeng Li, Adam R. Lovato, Andrew Symasek, Qingfei Zheng

## Abstract

Triazolinedione (TAD) derivatives have been commonly utilized as protection and labeling reagents for indole and phenol moieties via a reversible ene-type reaction. Previous studies showed that the TAD probes could selectively modify tyrosine and tryptophan side-chains within proteins and peptides under distinct pH conditions. Here, we report a pH-controlled regioselective rapid ene-type reaction (RRER) between TAD and 5-hydroxyindole, where the modification occurs on the C4 position rather than the C3 of inactivated indole rings. Employing this unique reaction, we have performed the selective bioconjugation of serotonylation occurring on the fifth amino acid residue, glutamine, of histone H3 (H3Q5), which does not contain any tryptophan in its protein sequence. Finally, RRER was applied to determine the H3Q5 serotonylation levels in cultured cells and tissue samples, which served as a newly developed powerful tool for in vitro and in vivo histone monoaminylation analysis. Overall, our findings in this research expanded the chemical biology toolbox for investigating histone monoaminylation and facilitated the understandings of TAD-mediated ene-type reactions.

## Introduction

The triazolinedione (TAD) reagents were originally developed and applied to protect the indole moieties in the total synthesis of natural products *via* a reversible ene-type reaction^1,2^, where the diazo structure of TAD attacks the indole C2 atom and thus protects its 2,3-π bond (Figure 1). Due to its rapidity, selectivity, and biocompatibility, this ene-type reaction has been used to label tyrosine in a selective manner, where TAD is conjugated to the ortho carbon (C3 or C5) of tyrosine phenolic hydroxyl group (Figure 1).^3,4,5^ Intriguingly, the bioconjugation between TAD and tryptophan was not observed under neutral or slightly alkaline pH.^3,4^ However, recent studies have shown that tryptophan is the preferred substrate over tyrosine under acidic pH (Figure 1).^6^ These results indicate that this ene-type reaction is pH-sensitive and its chemoselectivity is tunable. However, the reactivity of TADs against substituted indoles remains elusive due to the low abundance of substituted indole moieties in naturally occurring biomacromolecules.^7^

**Figure 1.**
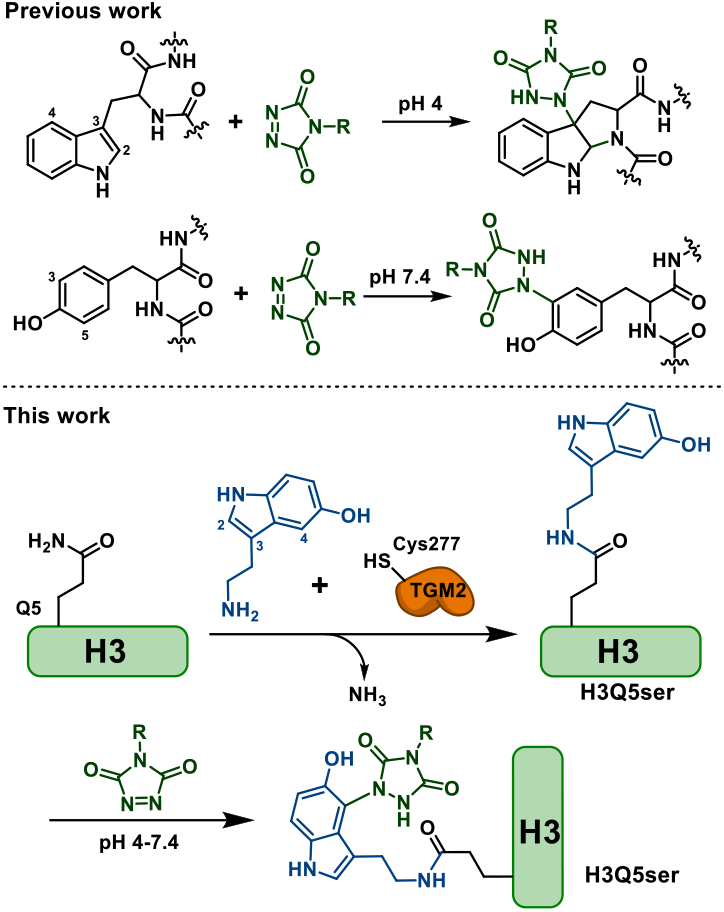
pH-Controlled chemo-and regio-selectivity of triazolinedione (TAD)-mediated ene-type reactions on tryptophan, tyrosine, and serotonin residues.

Recently, a novel histone post-translational modification (PTM) has been discovered, where the fifth amino acid residue, glutamine, of histone H3 (H3Q5) is modified by serotonin *via* an isopeptide bond.^8,9^ This enzymatic transamination reaction is catalyzed by transglutaminase 2 (TGM2) in a reversible manner (Figure 1).^10^ The H3Q5 serotonylation (H3Q5ser) has been shown to play a role in gene transcription and disease states including major depressive disorder^11^ and cancer.^12-14^ However, due to the lack of molecular tools, the efficient analysis of histone serotonylation *in vivo* and *in situ* remains a challenge.^9,15,16^ Notably, 5-hydroxyindole of H3Q5ser contains the only indole moiety as tryptophan is not present in histone proteins. Thus, we sought to utilize the TAD probe to label H3Q5ser in a selective fashion (Figure 1), however, the reactivity of TADs on 5-hydroxyindole-containing peptides/proteins have not been systematically studied.

## Results and discussion

To understand the TAD activities on 5-hydroxyindole-containing peptides, we first synthesized two tripeptides, AcNH-AWA-CONH_2_ and AcNH-AQserA-CONH_2_ (Figure 2), for the model reactions. The structures of these tripeptides were confirmed by using ^1^H and ^13^C nuclear magnetic resonance (NMR) and liquid chromatography-mass spectrometry (LC-MS). An alkynyl TAD (AlkTAD) probe was synthesized to react with the peptide substrates (Figure 2A). The structure of AlkTAD was characterized by using ^1^H and ^13^C NMR (*Supporting Information*). When AWA was incubated with AlkTAD under pH 4, as expected, AlkTAD selectively modified the C3 carbon of tryptophan indole and prompted the formation of a covalent bond between the C2 and α-amino nitrogen of tryptophan (Figure 2B). The AlkTAD-peptide adduct (**1**) structure was confirmed using ^1^H/^13^C NMR, heteronuclear multiple bond correlation (HMBC), and heteronuclear single quantum coherence (HSQC) (Figure 2B and *Supporting Information*). Notably, due to the dearomatization, **1** lost the characteristic ultraviolet (UV) absorption (λ = 280 nm) of indole (Figure 2C).^17^ Thereafter, AQserA was also incubated with AlkTAD and a new product was observed to form. Unexpectedly, distinct from **1**, the AlkTAD-peptide adduct (**2**) retained UV absorption at 280 nm, suggesting that **2** preserved the complete indole structure and did not undergo dearomatization during the addition reaction. Furthermore, 1D and 2D NMR showed that in this reaction TAD attacked the C4 carbon of 5-hydroxyindole and did not protect its 2,3-π bond (Figures 2B and *Supporting Information*). The pH gradient experiments indicated that the reactivity of 5-hydroxyindole-containing substrate could be significantly increased under higher pH values (Figures S4). These findings suggested that the substituents on indole influenced the regioselectivity of the ene-type reaction between TAD and indole moieties. Moreover, the obvious color change (from pink-red to colorless) of this ene-type reaction is a sensitive indicator for the accomplishment of bioconjugation and consumption of the probe (Figure 2D).

**Figure 2.**
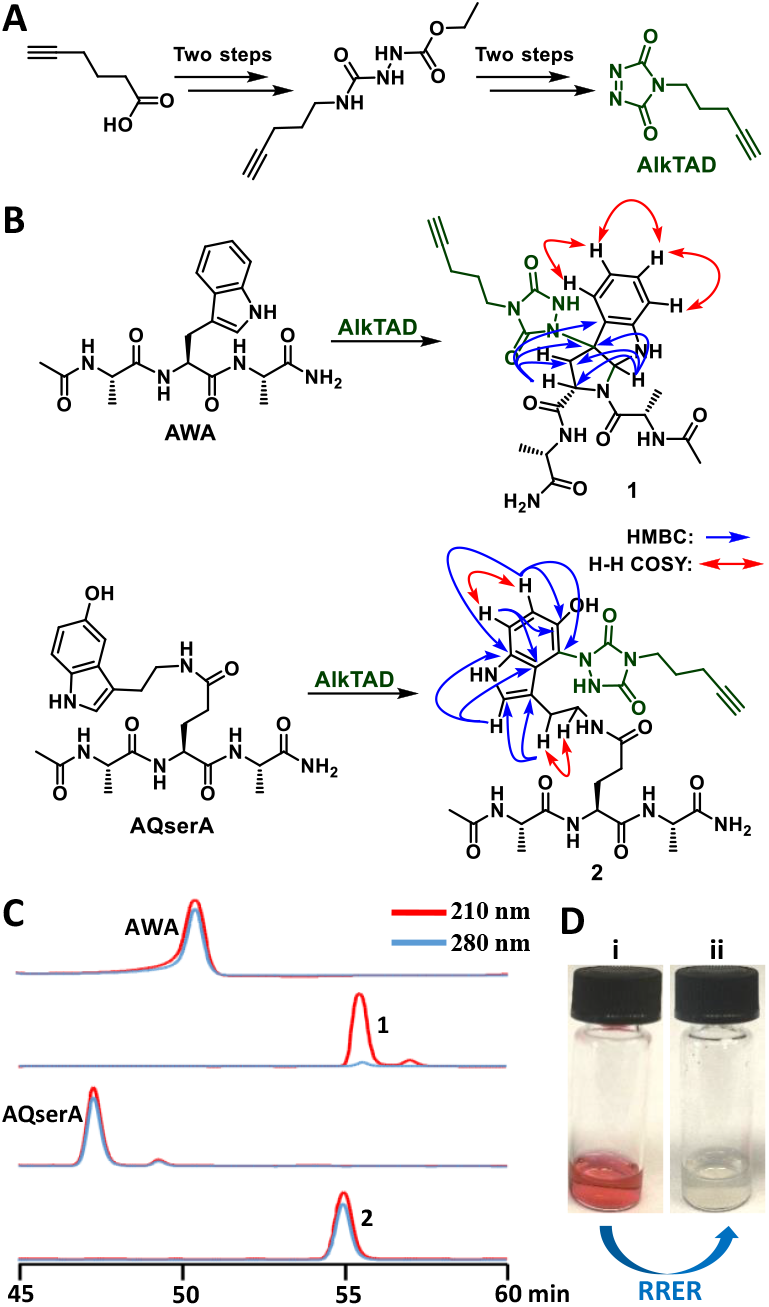
The regio-selectivity of AlkTAD-mediated RRER on indole and 5-hydroxyindole. (A) The 4-step synthesis of AlkTAD. (B) The structure validation of RRER products, where the peptides, AWA and AQserA, were used as model substrates. (C) HPLC and UV absorption wavelength (λ = 210 and 280 nm) analyses of the RRER substrates and corresponding products (**1** and **2**). (D) The obvious color change of RRER in 20 seconds: i. before the reaction; ii. after the reaction.

After discovering the addition between TAD and substituted indole is a pH-controlled regioselective rapid ene-type reaction (RRER), we tested the chemoselectivity and regioselectivity of the AlkTAD probe using diverse peptide substrates (**P1**-**P13**). These peptides shared a common backbone sequence with the N-terminal H3 (1-12 aa), however, they possessed different substitutions on their fifth amino acid residues which could react with AlkTAD *via* RRER (Figure 3A). The LC-MS analyses showed that the peptide substrates containing indole or phenol groups could form the adducts with the AlkTAD probe, while the other major nucleophiles in histones (such as imidazole of histidine and guanidine of arginine) did not react with AlkTAD (Figures 3B and S4). As the products of **P1**-**P13** contain distinct aromatic ring (or non-aromatic) structures and exhibited different UV-absorption features, the relative conversion rate of each reaction was determined by using the consumption ratio of the corresponding substrates (Figures 3B, 3C, and S4). Notably, these modified H3 peptide substrates exhibited different reactivity to AlkTAD under distinct pH values (Figure 3B). After the optimization of reaction conditions, we found that the serotonin residue-containing peptide, H3Q5ser (**P5**), is the most reactive substrate in comparison with other naturally occurring structures under acidic (pH 4) and low-temperature conditions (Figures 3C and S4). Similar to H3Q5W (**P2**),^6^ the reactivity of H3Q5dop (**P8**) was unexpectedly not sensitive to low pH. To avoid the possible cross-labelling by TAD, the dopamine residue on H3Q5dop was oxidized to a dopamine quinone (DAQ) structure (H3Q5daq, **P13**) by Fe (III), which is also a naturally occurring PTM (Figure 3A).^18^ Notably, **P13** could not react with TAD (Figures 3B, 3C, and S4), due to its electron-deficient structure, which is a bad nucleophile. Moreover, the substrates with unnatural substituents (such as fluorine and methoxy groups) further provided insights to the structure-reactivity-relationship of RRER, that is, the electron-rich substrates (*i*.*e*., good nucleophiles) are more reactive to TAD reagents. Additionally, based on the UV absorption results, the activated indole rings were easier to react with AlkTAD and did not undergo dearomatization (Figure S4). Overall, the *in vitro* reactions using modified peptides as substrates have shown that AlkTAD is a chemo-and regio-selective probe that can label the 5-hydroxy indole moieties within proteins in a rapid and specific manner. Importantly, in the presence of serotonylated peptides and under the low pH, other competitive substrates (*e*.*g*., tyrosine and dopaminylated/tyraminylated glutamine residues) cannot react with the TAD probe as priority.

**Figure 3.**
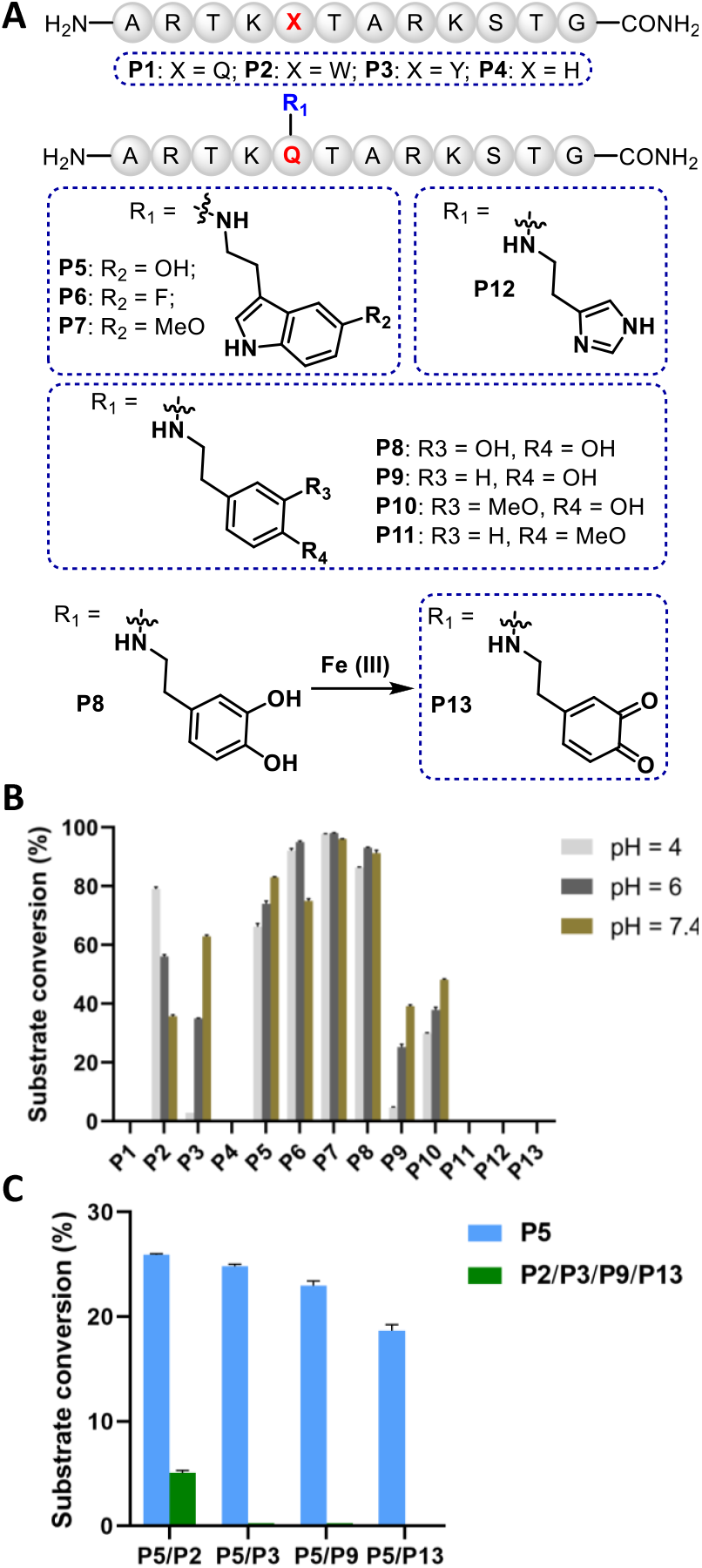
The optimization and application of RRER for selective bioconjugation of serotonylated histone H3 peptides. (A) The structures of H3 N-terminal peptide substrates. (B) The reactivity of AlkTAD to the peptides, **P1**-**P13**. The relative conversion rate was calculated based on the LC-MS analysis. (C) The competition reactions between reactive peptide substrates and AlkTAD, showing the serotonylated peptide (**P5**) is the most reactive in RRER. The error bars represent the standard deviation from three different experiments.

Given the reactivity of TAD probe on peptide substrates, we then performed the *in vitro* bioconjugation on nucleosome core particles (NCPs) that contained H3Q5ser. The serotonylated NCPs were prepared *via* the incubation with recombinant TGM2. The catalytically inactive mutant, TGM2-C277A, and non-reactive-site-containing substrate, NCP-H3Q5E, were used for the negative control experiments. The AlkTAD-modified proteins were conjugated with Cyanine5 (Cy5) azide through copper-catalysed azide-alkyne cycloaddition (CuAAC) and then visualized by using sodium dodecyl sulfate-polyacrylamide gel electrophoresis (SDS-PAGE) and in-gel fluorescence imaging.^9,10,13,15^ The in-gel imaging and immunoblotting analyses showed that H3Q5-serotonylated NCPs could be rapidly modified by AlkTAD *via* RRER in a selective fashion, while the unmodified wild-type (WT) H3Q5 NCPs or H3E5 mutation were not able to react with the probe (Figure 4A). This further confirms that AlkTAD can be utilized as a selective probe to label TGM2-catalyzed histone serotonylation. Notably, the core histones within recombinant NCPs do not possess any tryptophan residue or indole moiety. Even though the wild-type NCPs are rich in tyrosine, only the enzymatically H3Q5-serotonylated NCPs can be labeled by the AlkTAD probe *via* RRER under the optimized reaction conditions.

**Figure 4.**
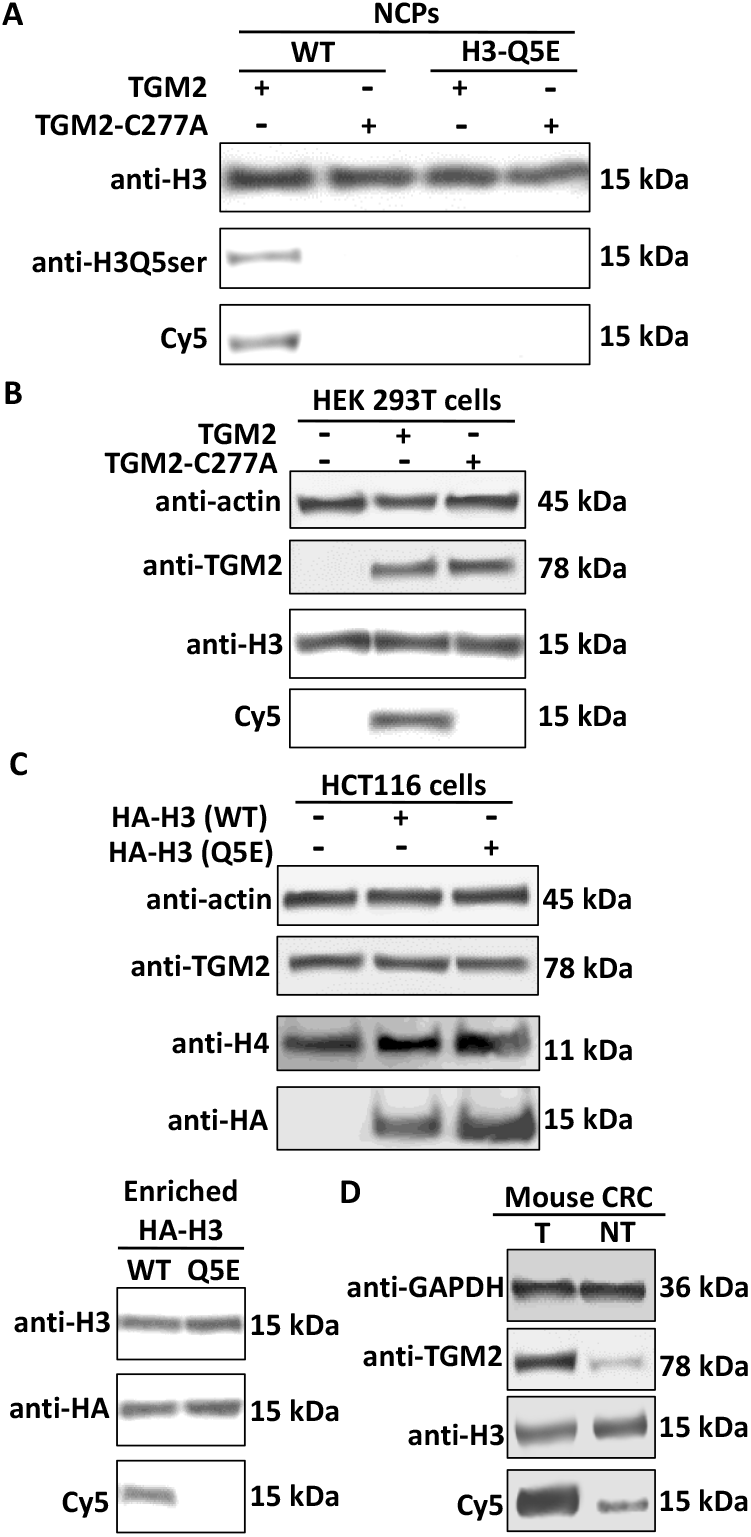
The in-gel imaging and immunoblotting analyses of AlkTAD-labeled histone serotonylation *via* RRER. (A) *In vitro* labeling of histone serotonylation, where NCPs were used as substrates and the site-specific antibody anti-H3Q5ser was employed as a positive control. (B) RRER-mediated labeling of serotonylated histones extracted from HEK 293T cells, which do not naturally express TGM2. (C) RRER-mediated labeling of HA-tagged serotonylated histones (WT H3 and H3-Q5E) from HCT116 cells, which naturally express TGM2. Histone H4 was used as a loading control. (D) RRER-mediated labeling of serotonylated histones extracted from tumor (T) and non-tumor (NT) samples of mouse CRC.

To analyze the histone serotonylation levels in cultured cell and tissue samples using the AlkTAD probe and RRER, we extracted the histone fractions from these samples and performed bioorthogonal labeling and in-gel imaging to illustrate H3Q5 serotonylation. First, wild-type TGM2 and its mutant TGM2-C277A were overexpressed in HEK 293T cells, a naturally occurring negative control cell line for monoaminylation studies, due to its *tgm2* gene silencing.^9,10,13^ After the serotonin treatment, the cells were harvested and their histone fractions were purified using acid extraction.^12^ The extracted histones were treated with K_3_[Fe(CN)_6_] and then incubated with AlkTAD under pH 4 on ice, coupled with Cy5 azide *via* CuAAC, and then imaged after SDS-PAGE. It showed that WT-TGM2-catalyzed cellular H3Q5ser could be specifically illustrated by AlkTAD through RRER (Figure 4B). In the previous studies, we have found that H3Q5 serotonylation is highly enriched in colorectal cancer (CRC).^13^ Here, we overexpressed HA-tagged WT H3 as well as H3-Q5E (*i*.*e*., a negative control that cannot be serotonylated by TGM2) in HCT116 cells and treated the transfected cells with serotonin. Thereafter, the HA-tagged H3 was purified by using anti-HA magnetic beads and the corresponding H3Q5ser levels were analyzed by using the AlkTAD probe. The in-gel imaging results indicated that histone serotonylation occurred on H3Q5 and accumulated in colon cancer cells (Figure 4C). Finally, to analyze the histone serotonylation levels *in vivo*, the CRC and non-tumor (NT) tissues were collected from C57BL/6J Apc^Min/+^ mice and their histones were extracted *via* acid extraction.^10,13,19^ The AlkTAD-based chemical biology imaging further confirmed that histone H3 serotonylation was highly enriched in CRC tissues, due to the much higher expression level of TGM2 compared with NT tissues (Figure 4D).^13,14^

## Conclusions

The TAD-mediated reversible ene-type reactions have been well established in organic chemistry to protect the 2,3-π bond of indole and wildly applied in bioconjugations to specifically label tyrosine residues within proteins.^1-5^ Recent studies showed that the chemoselectivity of the TAD-mediated ene-type reaction was tunable under different pH values.^6^ Specifically, tryptophan and tyrosine are the preferred substrates of TAD under low and high pH, respectively. However, the reaction characteristics of TAD to substituted indole, especially the activated indole rings, remains poorly understood due to their low abundance in natural biomacromolecules. In our previous work, we have uncovered the detailed enzymatic mechanism TGM2-catalyzed protein monoaminylation, which can introduce 5-hydroxy indole onto glutamine residues through the transamination with serotonin.^10,13^ In this study, we for the first time, found that TAD could rapidly and selectively conjugate the C4 of 5-hydroxy indole in serotonylated peptides/proteins, without the protection of indole 2,3-π bond. This unique chemoselective reaction was thus named as RRER and utilized by us to study a newly identified epigenetic mark on histone, *i*.*e*., H3Q5ser.

Recently, we have developed a series of chemical biology tools to study histone monoaminylation.^13,18,20^ Utilizing these powerful tools, we reported the accumulation of H3Q5 monoaminylation in different types of cancer cells as well as its regulatory function in controlling the cellular chromatin structure and gene transcription.^13^ These findings provided a novel link between the monoamine metabolism and epigenetic regulations. Briefly, serotonin is biosynthesized from the indispensable amino acid, tryptophan, by the decarboxylase and oxidase of the gut microbiome and host cells.^21,22^ The discovery of histone serotonylation offered a new hub connecting the gut microbiome, diet, signal transduction, and disease, especially cancer. However, due to the lack of efficient analytical tools, visualization and detection of histone serotonylation levels within cell and tissue samples remains a challenge. For example, the site-specific antibody for H3Q5ser could not be employed for the analysis of cultured cells or tissues.^9,10,13^ In comparison with other reported bioconjugation approaches for aromatic ring-containing amino acid residues in proteins,^23-34^ RRER is not only easy to operate, without the usage of harsh conditions (*e*.*g*., transition metal, free radicals, or electrochemistry), but also has a wider application range for selective labeling of diverse residues, which can be precisely tuned by pH values. The unique reaction properties of RRER and successful applications of the AlkTAD probe in this study not only facilitated our understandings of the selectivity of TAD-mediated ene-type reactions,^1-6^ but also further expanded the chemical biology toolbox for investigating histone monoaminylation and chemical epigenetics,^35,36^ paving the way to future discoveries of novel drugs targeting serotonylation-representative indole-containing proteome in cancer cells.^16,20^

## Data availability

The data supporting this article have been included as part of the ESI† available at [https://doi.org/XXXXXXXXXXXX].

## Acknowledgements

This research work was financially supported by the NIH (R35 GM150676) and startup funds from Purdue University for Q.Z.

---

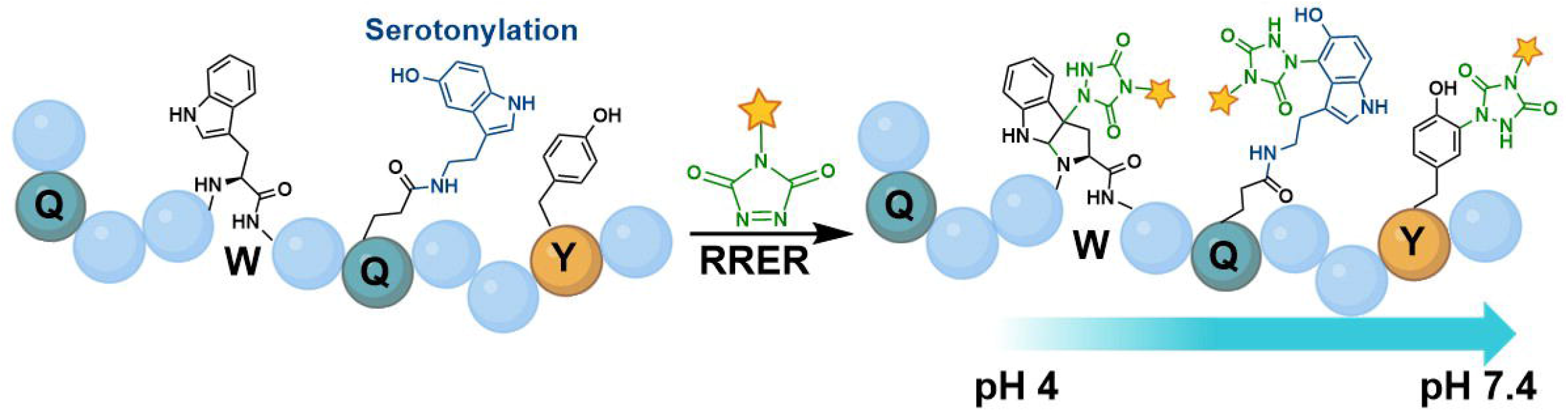

## Notes and references

1. P. S. Baran, C. A. Guerrero, E. J. Corey, Short, enantioselective total synthesis of okaramine N, J. Am. Chem. Soc. 2003, 125, 5628–5629.

2. P. S. Baran, C. A. Guerrero, E. J. Corey, The first method for protection-deprotection of the indole 2,3-π bond, Org. Lett. 2003, 5, 1999–2001.

3. H. Ban, J. Gavrilyuk, C. F. Barbas, Tyrosine bioconjugation through aqueous ene-type reactions, J. Am. Chem. Soc. 2010, 132, 1523–1525.

4. Q.-Y. Hu, M. Allan, R. Adamo, D. Quinn, H. Zhai, G. Wu, K. Clark, J. Zhou, S. Ortiz, B. Wang, E. Danieli, S. Crotti, M. Tontini, G. Brogioni, F. Berti, Synthesis of a well-defined glycoconjugate vaccine by a tyrosine-selective conjugation strategy, Chem. Sci., 2013, 4, 3827–3832.

5. D. Alvarez-Dorta, C. Thobie-Gautier, M. Croyal, M. Bouzelha, M. Mevel D., Deniaud, M. Boujtita, S. G. Gouin, Electrochemically promoted tyrosine-click-chemistry for protein labelling. J. Am. Chem. Soc. 2018, 140, 17120–17126.

6. K. W. Decoene, K. Unal, A. Staes, O. Zwaenepoel, J. Gettemans, K. Gevaert, J. M. Winne and A. Madder, Triazolinedione protein modification: From an overlooked off-target effect to a tryptophan-based bioconjugation strategy. Chem. Sci., 2022, 13, 5390–5397.

7. K. K. Kaushik, N. Kaushik, P. Attri, N. Kumar, C. H. Kim, A. K. Verma and E. H. Choi, Biomedical importance of indoles. Molecules 2013, 18, 6620–6662.

8. A. Al-Kachak, I. Maze, Post-translational modifications ofhistone proteins by monoamine neurotransmitters. Curr. Opin. Chem. Biol. 2023, 74, 102302.

9. L. A. Farrelly, R. E. Thompson, S. Zhao, A. E. Lepack, Y. Lyu, N. V. Bhanu, B. Zhang, Y. E. Loh, A. Ramakrishnan, K. C. Vadodaria, K. J. Heard, G. Erikson, T. Nakadai, R. M. Bastle, B. J. Lukasak, H. Zebroski, N. Alenina, M. Bader, O. Berton, R. G. Roeder, H. Molina, F. H. Gage, L. Shen, B. A. Garcia, H. Li, T. W. Muir and I. Maze, Histone serotonylation is a permissive modification thatenhances TFIID binding to H3K4me3. Nature 2019, 567, 535–539.

10. Q. Zheng, B. H. Weekley, D. A. Vinson, S. Zhao, R. M. Bastle, R. E. Thompson, S. Stransky, A. Ramakrishnan, A. M. Cunningham, S. Dutta, J. C. Chan, G. Di Salvo, M. Chen, N. Zhang, J. Wu, S. L. Fulton, L. Kong, H. Wang, B. Zhang, L. Vostal, A. Upad, L. Dierdorff, L. Shen, H. Molina, S. Sidoli, T. W. Muir, H. Li, Y. David and I. Maze, Bidirectional, histone monoaminylation dynamics regulate neural rhythmicity. Nature 2025, 637, 974–982.

11. A. Al-Kachak, G. Di Salvo, S. L. Fulton, J. C. Chan, L. A. Farrelly, A. E. Lepack, R. M. Bastle, L. Kong, F. Cathomas, E. L. Newman, C. Menard, A. Ramakrishnan, P. Safovich, Y. Lyu, H. E. Covington, L. Shen, K. Gleason, C. A. Tamminga, S. J. Russo and I. Maze, Histone serotonylation in dorsal raphe nucleus contributes to stress- and antidepressant-mediated gene expression and behavior. Nat. Commun. 2024, 15, 5042.

12. H.-C. Chen, P. He, M. McDonald, M. R. Williamson, S. Varadharajan, B. Lozzi, J. Woo, D.-J. Choi, D. Sardar, E. Huang-Hobbs, H. Sun, S. M. Ippagunta, A. Jain, G. Rao, T. E. Merchant, D. W. Ellison, J. L. Noebels, K. C. Bertrand, S. C. Mack and B. Deneen, Histone serotonylation regulates ependymoma tumorigenesis. Nature 2024, 632, 903–910.

13. N. Zhang, J. Wu, F. Hossain, H. Peng, H. Li, C. Gibson, M. Chen, H. Zhang, S. Gao, X. Zheng, Y. Wang, J. J. Wang, I. Maze and Q. Zheng, Bioorthogonal labeling and enrichment ofhistone monoaminylation reveal its accumulation and regulatoryfunction in cancer cell chromatin. J. Am. Chem. Soc. 2024, 146, 16714–16720.

14. Li, H.; Wu, J.; Zhang, N.; Zheng, Q. Transglutaminase 2-mediated histone monoaminylation and its role in cancer. Biosci. Rep. 2024, 44, BSR20240493.

15. J. C.-Y. Lin, C.-C. Chou, S. Gao, S.-C. Wu, K.-H. Khoo and C.-H. Lin, An in vivo tagging method reveals that Ras undergoes sustained activation upon transglutaminase-mediated protein serotonylation. ChemBioChem 2013, 14, 813–817.

16. N. Zhang, J. Wu, Q. Zheng, Chemical proteomics approaches for protein post-translational modification studies. Biochim. Biophys. Acta Proteins Proteom. 2024, 1872, 141017.

17. S. P. Roche, J-J. Y. Tendoung, B. Tréguier, Advances in dearomatization strategies of indoles. Tetrahedron, 2015, 71, 3549–3591.

18. N. Zhang, S. Gao, H. Peng, J. Wu, H. Li, C. Gibson, S. Wu, J. Zhu, Q. Zheng, Chemical proteomic profiling of protein dopaminylation in colorectal cancer cells. J. Proteome Res. 2024, 23, 2651–2660.

19. D. Shechter, H. L. Dormann, C. D. Allis and S. B. Hake, Extraction, purification and analysis of histones. Nat. Protoc. 2007, 2, 1445– 145.

20. N. Zhang, J. Wu, S. Gao, H. Peng, H. Li, C. Gibson, S. Wu, J. Zhu, Q. Zheng, pH-Controlled chemoselective rapid azo-coupling reaction (CRACR) enables global profiling of serotonylation proteome in cancer cells. J. Proteome Res. 2024, 23, 4457–4466.

21. Y. Hou, J. Li, S. Ying, Tryptophan metabolism and gut microbiota: A novel regulatory axis integrating the microbiome, immunity, and cancer. Metabolites 2023, 13, 1166.

22. C. Li, Y. Liang, Y. Qiao, Messengers from the gut: Gut microbiota-derived metabolites on host regulation. Front. Microbiol. 2022, 13, 863407.

23. P. S. Addy, S. B. Erickson, J. S. Italia, A. Chatterjee, d A Chemoselective rapid azo-coupling reaction (CRACR) for unclickable bioconjugation. J. Am. Chem. Soc. 2017, 139, 11670– 11673.

24. F. Feng, Y. Gao, Q. Zhao, T. Luo, Q. Yang, N. Zhao, Y. Xiao, Y. Han, J. Pan, S. Feng, L. Zhang, M. Wu, Single-electron transfer between sulfonium and tryptophan enables site-selective photo crosslinking of methyllysine reader proteins. Nat. Chem. 2024. 16, 1267–1277.

25. J. M. Antos, J. M. McFarland, A. T. Iavarone, M. B. Francis, Chemoselective tryptophan labeling with rhodium carbenoids at mild pH. J. Am. Chem. Soc. 2009, 131, 6301–6308.

26. Y. Seki, T. Ishiyama, D. Sasaki, J. Abe, Y. Sohma, K. Oisaki, M. Kanai, Transition metal-free tryptophan-selective bioconjugation of proteins. J. Am. Chem. Soc. 2016, 138, 10798–10801.

27. S. J. Tower, W. J. Hetcher, T. E. Myers, N. J. Kuehl and M. T. Taylor, Selective modification of tryptophan residues in peptides and proteins using a biomimetic electron transfer process. J. Am. Chem. Soc. 2020, 142, 9112–9118.

28. Y. Yu, L. K. Zhang, A. V. Buevich, G. Li, H. Tang, P. Vachal, S. L. Colletti and Z. C. Shi, Chemoselective peptide modification via photocatalytic tryptophan β-position conjugation. J. Am. Chem. Soc. 2018, 140, 6797–6800.

29. L. E. Guzmán, A. N. Wijetunge, B. F. Riske, B. B. Massani, M. A. Riehle and J. C. Jewett, Chemical probes to interrogate the extreme environment of mosquito larval guts. J. Am. Chem. Soc. 2024, 146, 8480–8485.

30. D. Alvarez Dorta, D. Deniaud, M. Mével and S. G. Gouin, Tyrosine conjugation methods for protein labelling. Chemistry 2020, 26, 14257–14269.

31. X. Xie, P. J. Moon, S. W. M. Crossley, A. J. Bischoff, D. He, G. Li, N. Dao, A. Gonzalez-Valero, A. G. Reeves, J. M. McKenna, S. K. Elledge, J. A. Wells, F. D. Toste and C. J. Chang, Oxidative cyclization reagents reveal tryptophan cation-π interactions. Nature 2024, 627, 680–687.

32. N. L. Kjaersgaard, T. B. Nielsen and K. V. Gothelf, Chemical conjugation to less targeted proteinogenic amino acids. ChemBioChem 2022, 23, e202200245.

33. C. Wan, R. Sun, W. Xia, H. Jiang, B. X. Chen, P. C. Kuo, W. R. Zhang, G. Yang, D. Li, C. W. Chiang and Y. Weng, Electrochemical bioconjugation of tryptophan residues: A strategy for peptide modification. Org. Lett. 2024, 5, 5447–5452.

34. M. S. Kang, T. W. S. Kong, J. Y. X. Khoo and T. P. Loh, Recent developments in chemical conjugation strategies targeting native amino acids in proteins and their applications in antibody-drug conjugates. Chem. Sci. 2021, 12, 13613–13647.

35. J. N. Beyer, N. R. Raniszewski and G. M. Burslem, Advances and opportunities in epigenetic chemical biology. ChemBioChem 2021, 22, 17–42.

36. Y. David, T. W. Muir, Emerging chemistry strategies for engineering native chromatin. J. Am. Chem. Soc. 2017, 139, 9090–9096.

