## Supplemental Information for "Regioselective Rapid Ene-type Reaction (RRER) Enables Bioconjugation of Histone Serotonylation"

### Table of Contents

1. General experimental procedures ... 5

2. Synthesis of AlkTAD probe ... 6

2.1 Synthesis of **S2** ... 6

2.2 Synthesis of **S4** ... 7

2.3 Synthesis of AlkTAD ... 7

3. Synthesis of peptides ... 8

3.1 Synthesis of acetonide-protected dopamine **S7** ... 8

3.2 Synthesis of trityl-protected histamine **S10** ... 9

3.3 List of synthetic peptides used in this study ... 10

3.5 Synthesis of AcNH-AWA-$CONH_2$ and AcNH-AQserA-$CONH_2$ peptides ... 11

3.5.1 AcNH-AWA-$CONH_2$ ... 11

3.5.2 AcNH-AQserA-$CONH_2$ ... 11

4. Reactions between peptides and AlkTAD ... 12

4.1 Reaction between AcNH-AWA-$CONH_2$ and AlkTAD ... 12

4.2 Reaction between AcNH-AQserA-$CONH_2$ and AlkTAD ... 13

4.3 Reaction between AcNH-AWA-$CONH_2$/AcNH-AQserA-$CONH_2$ with AlkTAD at different pH values ... 15

4.4 General procedure for reactions between peptides and AlkTAD ... 15

4.5 AlkTAD-peptide reactions at different pH values ... 15

4.6 The competition reaction between two peptide substrates and AlkTAD ... 24

4.6.1 H3Q5ser and H3W5 ... 24

4.6.2 H3Q5ser and H3Y5 ... 24

4.6.3 H3Q5ser and H3Q5dop ... 25

4.6.4 H3Q5ser and H3Q5tyr ... 25

4.6.5 H3Q5ser and H3Q5daq ... 26

5. *In vitro* labeling assays ... 26

6. Expression of TGM2 in HEK 293T cells ... 27

7. Expression of HA-tagged WT H3 and H3-Q5E in HTC116 cells ... 28

8. Extraction of histones from cultured cells ... 28

9. Cell fractionation ... 28

#### 1. General experimental procedures

UV spectrometry was performed on a NanoDrop 2000c (Thermo Scientific). Biochemicals and media were purchased from Fisher Scientific or Sigma-Aldrich Corporation unless otherwise stated. T4 DNA ligase, DNA polymerase and restriction enzymes were obtained from New England BioLabs. PCR amplifications were performed on an Applied Biosystems Veriti Thermal Cycler using either Taq DNA polymerase (Vazyme Biotech) for routine genotype verification or Phanta Max Super-Fidelity DNA Polymerase (Vazyme Biotech) for high-fidelity amplification. Site-specific mutagenesis was performed according to standard procedures of the QuickChange Site-Directed Mutagenesis Kit purchased from Stratagene (GE Healthcare) or Mut Express II (Vazyme Biotech). Primer synthesis and DNA sequencing were performed by Integrated DNA Technologies and Genewiz, respectively. PCR amplifications were performed on a Bio-Rad T100TM Thermal Cycler. Centrifugal filtration units were purchased from Millipore, and MINI dialysis units purchased from Pierce. Size exclusion chromatography was performed on an AKTA FPLC system from GE Healthcare equipped with a P-920 pump and UPC-900 monitor. Sephacryl S-200 columns were obtained from GE Healthcare. All the western blots were performed using the primary antibodies annotated in **Supplementary Table 1** and fluorophore-labeled secondary antibodies annotated in **Supplementary Table 2** following protocols recommended by the manufacture. Blots were imaged on an Odyssey CLx Imaging System (Li-Cor). Amino acid derivatives and coupling reagents were purchased from AGTC Bioproducts. Dimethylformamide (DMF), dichloromethane (DCM) and triisopropylsilane (TIS) were purchased from Fisher Scientific and used without further purification. Hydroxybenzotriazole (HOBt) and O-(benzotriazol-1-yl)-N,N,N′,N′-tetramethyluronium hexafluorophosphate (HBTU) were purchased from Fisher Scientific. Trifluoroacetic acid (TFA) was purchased from Fisher Scientific. N,Ndiisopropylethylamine (DIPEA) was purchased from Fisher Scientific. Analytical reversed-phase HPLC (RP-HPLC) was performed on an Agilent 1200 series instrument with an Agilent C18 column (5 μm, 4 × 150 mm), employing 0.1% TFA in water (HPLC solvent A), and 0.1% TFA in acetonitrile (HPLC solvent B) as the mobile phases. Analytical gradients were 0-70% HPLC buffer B over 45 minutes at a flow rate of 0.5 mL/minute, unless stated otherwise. Preparative scale purifications were conducted on an Agilent LC system. An Agilent C18 preparative column (15-20 μm, 20 × 250 mm) or a semi-preparative column (12 μm, 10 mm × 250 mm) was employed at a flow rate of 20 mL/min or 4 mL/min, respectively. HPLC Electrospray ionization MS (HPLC-ESI-MS) analysis was performed on an Agilent 6120 Quadrupole LC/MS spectrometer (Agilent Technologies) or LCMS-2050 Single Quadrupole LC/MS spectrometer (Shimadzu Scientific Instruments). All immunoblotting experiments in this research were performed at least 3X. For the synthesis of probe molecules used in this study, all commercial chemicals were purchased from Sigma Aldrich, TCI chemicals, AK Scientific, Fischer Scientific, Broadpharm and used without further purification. The reagents and solvents were handled following the safety processes required as instructed by the manufacturer. Organic and aqueous waste was disposed of following standard safety protocols. Solvents for workup were purchased from Fisher Chemical, and anhydrous solvents were purchased from Sigma Millipore in a sealed bottle and degassed by passing $N_2$ before each use. Reaction progress was monitored by using normal phase TLC silica gel 60 $F_{254}$ plates by Sigma Aldrich (aluminum backed 20 X 20 cm). Developed plates were analyzed by visualizing under a UV-light and/or staining with Phosphomolybdic Acid (PMA) Stain (100 mL absolute ethanol and 10 g PMA). Isolation and purification of the crude reaction materials were performed using silica gel ($SiO_2$) by Acros Organic (0.030-0.200 mm, 60 $A^0$). Organic solvents were removed under vacuum using a Heidolph Rotavapor equipped with a dry ice condenser. Deuterated solvents (such as $D_2O$, DMSO-$d_6$) for NMR characterization were purchased from Sigma Aldrich. $^1$H NMR, 13C NMR data were recorded on a Bruker Avance-600/700 MHz spectrometer at 22 °C. Chemical shifts (δ) are reported in ppm to the internal standard of residual $D_2O$ (δ 4.79: $^1$H NMR). All the $^1$H NMR are reported as follows; chemical shift (δ ppm), multiplicity (s, singlet; d, doublet; t, triplet; q, quartet; m, multiplet; dd, doublet of doublets), coupling constant (Hz), integration and assigned proton. Data for $^{13}$C NMR spectroscopy are reported in chemical shift (δ ppm).

#### 2. Synthesis of AlkTAD probe

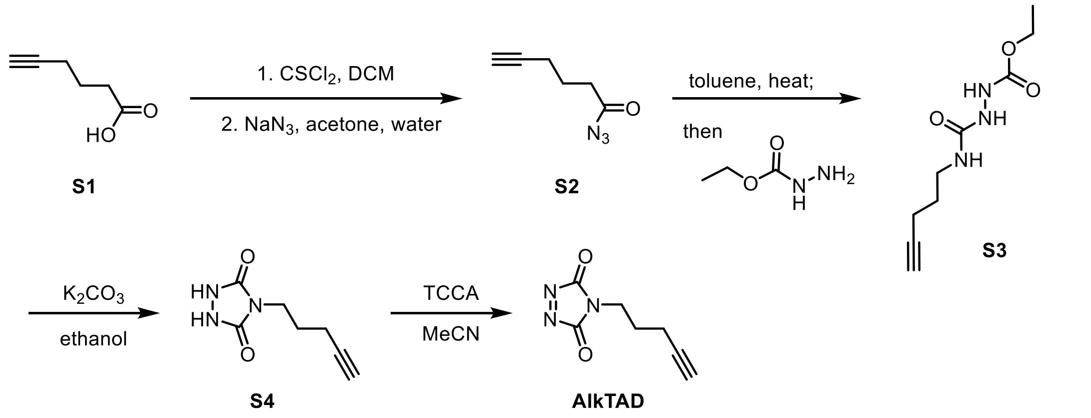

##### 2.1 Synthesis of **S2**

To a solution of **S1** (112 mg, 1.0 mmol, 1.0 equiv.) in DCM (5 mL) was added thiophosgene (230 mg, 2.0 mmol, 2.0 equiv.) and the mixture was stirred at 0 °C for 2 h. The mixture was evaporated under reduced pressure. The residue was dissolved in acetone (5 mL), and added $NaN_3$ (2.0 mmol, 1.0 mL, 2 M in $H_2O$, 2.0 equiv.), and the mixture was stirred at 0 °C. After stirred for 1 h, the reaction mixture was added water (5 mL) and extracted with EtOAc three times. The combined organic layers were washed with brine, dried over sodium sulfate, filtered, concentrated and purified by column chromatography on silica gel (petroleum ether : EtOAc = 10 : 1) to give **S2** as a colorless oil (126 mg, 92%).

#### 2.2 Synthesis of **S4**

To a solution of **S2** (129 mg, 0.92 mmol, 1.0 equiv.) obtained above in toluene (5 mL) and the mixture was stirred at 120 °C. After stirred for 0.5 h, ethyl carbazate (143mg, 1.38 mmol, 1.5 equiv.) was add and the reaction mixture was stirred at 120 °C for 3 hours. The mixture was evaporated under reduced pressure to get crude **S3**. The residue (crude **S3**) was dissolved in ethanol (5 mL), and added $K_2CO_3$ (190 mmol, 1.38 mmol, 1.5 equiv.), and the mixture was stirred at 78 °C overnight. The reaction mixture was filtered through a celite pad and the filtrate was concentrated and purified by column chromatography on silica gel (DCM : MeOH = 30 : 1) to give **S4** as a white solid (87 mg, 52% for 4 steps).

$^1$H NMR (600 MHz, DMSO-*d*$_6$) δ 11.06 (s, 1H), 7.77 (dd, *J* = 8.5, 7.3 Hz, 1H), 7.47 (d, *J* = 8.5 Hz, 1H), 7.40 (d, *J* = 7.2 Hz, 1H), 5.04 (dd, *J* = 13.0, 5.5 Hz, 1H), 4.21 (t, *J* = 6.0 Hz, 2H), 3.72 (t, *J* = 6.5 Hz, 2H), 2.88 – 2.80 (m, 1H), 2.58 – 2.52 (m, 1H), 2.47 – 2.52 (m, 2H), 2.01 – 1.96 (m, 1H), 1.91 – 1.82 (m, 4H).

$^{13}$C NMR (150 MHz, DMSO-*d*$_6$) δ 173.3, 170.4, 167.3, 165.8, 156.3, 137.5, 133.7, 120.2, 116.7, 115.7, 68.7, 49.2, 45.6, 45.6, 31.4, 29.3, 26.3, 22.5.

#### 2.3 Synthesis of AlkTAD

To a solution of **S4** (16.7 mg, 0.1 mmol, 1.0 equiv.) obtained above in MeCN (10 mL) was added trichloroisocyanuric acid (TCCA, 7.7 mg, 0.033 mmol, 0.33 equiv) the mixture was stirred at 25 °C for 2 hours. The mixture was centrifugated and collected the supernatant, and a MeCN solution of AlkTAD with a concentration of 10 mM was provide.

$^1$H NMR (700 MHz, $CD_3CN$) δ 3.67 (t, *J* = 7.0 Hz, 2H), 2.25 (td, *J* = 6.9, 2.7 Hz, 3H), 2.21 (t, *J* = 2.7 Hz, 1H), 1.85 (dt, *J* = 13.1, 6.7 Hz, 3H).

$^{13}$C NMR (175 MHz, $CD_3CN$) δ 196.6, 159.9, 82.8, 69.7, 40.2, 25.7, 15.3.

#### 3. Synthesis of peptides

Standard Fmoc-based Solid Phase Peptide Synthesis (FmocSPPS) was used for the synthesis of peptides in this study. Generally, the peptides were synthesized on ChemMatrix resins with Rink Amide to generate C-terminal amides. Peptides were synthesized using manual addition of the reagents (using a stream of dry $N_2$ to agitate the reaction mixture). For amino acid coupling, 5 equiv. Fmoc protected amino acid were pre-activated with 4.9 equiv. HBTU, 5 equiv. HOBt and 10 equiv. DIPEA in DMF and then reacted with the N-terminally deprotected peptidyl resin. Fmoc deprotection was performed in an excess of 20% (v/v) piperidine in DMF, and the deprotected peptidyl resin was washed thoroughly with DMF to remove trace piperidine. Cleavage from the resin and side-chain deprotection were performed with 95 % TFA, 2.5% TIS and 2.5% $H_2O$ at room temperature for 1.5 hours. The peptides were then precipitated with cold diethyl ether, isolated by centrifugation and dissolved in water with 0.1 % formic acid followed by RP-HPLC and ESI-MS analyses. Preparative RP-HPLC was used to purify the peptides of interest.[1,2]

For the synthesis of site-specific monoaminylated H3 peptides, Fmoc-Glu(Oall)-OH was incorporated at position 5 for the orthogonal deprotection and further monoaminylation. Briefly, the protect group Oall were removed by $Pd(PPh_3)_4$ and $PhSiH_3$ and then conjugated with monoamine donors (i.e., serotonin hydrochloride, and acetonide-protected dopamine) in conditions of PyAOP and DIEA. During synthesis of H3Q5dop and H3Q5his, acetonide-protected dopamine block **S7** and Trt-histamine block **S10** was used.[1,2]

For the synthesis of dopamine quinone (DAQ)-modified H3 peptides (H3Q5daq, **P13**): H3Q5dop (100 µL, 6 mM) was mixed with $FeCl_3$ (100 µL, 60 mM) in aqueous solution. The reaction mixture was gently shaken at room temperature for 10 minutes to yield H3Q5daq (200 µL, final concentration: 3 mM). The product was confirmed by LC-MS analysis.

##### 3.1 Synthesis of acetonide-protected dopamine **S7**

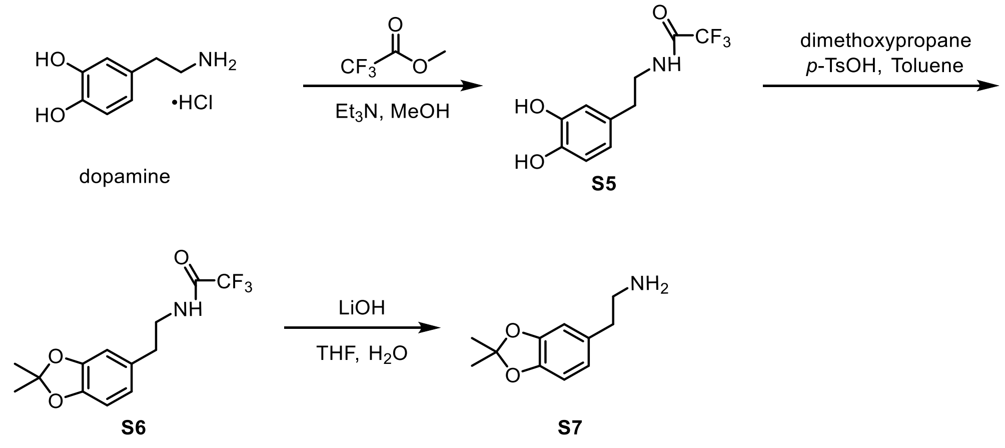

To a solution of dopamine hydrochloride (1.25 g, 6.6 mmol, 1.0 equiv.) obtained above in MeOH (15 mL) was added methyl trifluoroacetate (1.69 g, 13.2 mmol, 2.0 equiv.) and triethylamine (2.67 g, 26.4 mmol, 4.0 equiv.), the mixture was stirred at 60 °C overnight. The mixture was concentrated, and the residue was added toluene (120 mL), dimethoxypropane (5.7 mL) and *p*-TsOH· $H_2O$ (125 mg, 0.66 mmol, 0.1 equiv.). The mixture was stirred at 125 °C overnight. The mixture was concentrated and the residue was purified through column chromatograph on silica gel (hexane : EtOAc 20:1) to give 405 mg **S6** as yellow solid (21% for 2 steps)**.**

$^1$H NMR (600 MHz, $CDCl_3$) δ 6.80 – 6.74 (m, 1H), 6.64 – 6.61 (m, 1H), 6.57–6.52 (m, 2H), 3.51 (q, $J$ = 6.8 Hz, 2H), 2.74 (t, $J$ = 7.2 Hz, 2H), 1.62 (s, 6H).

To a solution of **S6** (289 mg,1.0 mmol, 1.0 equiv.) obtained above in THF (10 mL) and $H_2O$ (1 mL) was added LiOH (36 mg, 1.5 mmol, 1.5 equiv), the mixture was stirred in 0 °C for 4 hours. The reaction was quenched with $NaHCO_3$ saturated solution and extracted with EtOAc (3 × 10 mL). The combined organic layers were washed with brine, dried over $Na_2SO_4$, filtered and concentrated in vacuo. The residue was dissolved in 5 ml DMF to obtain 0.2 M **S7** solution, and used in peptides synthesis without purification.

3.2 Synthesis of trityl-protected histamine **S10**

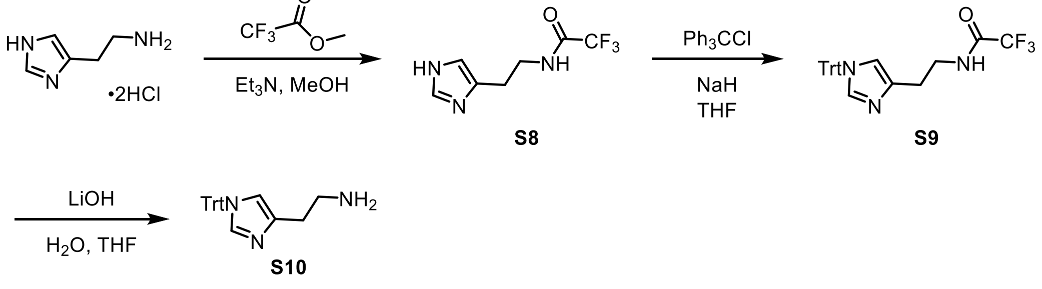

To a solution of histamine dihydrochloride (1.474 g, 8.0 mmol) obtained above in 15 mL MeOH was added methyl trifluoroacetate (1.64 g, 12.8 mmol, 1.6 equiv.) and triethylamine (2.02 g, 20 mmol, 2.5 equiv.) was added to the mixture. The mixture was stirred at 60 °C overnight, and concentrated to obtain crude **S8**.

To a solution of crude **S8** (8.0 mmol, 1.0 equiv.) obtained above in 40 mL THF was added NaH (0.35 g, 8.8 mmol, 60% in oil, 1.1 equiv.) at ice bath and stirred at 0 ºC for 30 min. The mixture was added triphenylmethyl (Trt) chloride (3.3 g, 12 mmol, 1.5 equiv.), and warmed to RT. After stirring 30min, the reaction was quenched with $NaHCO_3$ saturated solution and extracted with EtOAc (3 × 10 mL). The combined organic layers were washed with brine, dried over $Na_2SO_4$, filtered and concentrated in vacuo to obtain crude **S9**.

To a solution of **S9** (8.0 mmol, 1.0 equiv.) obtained above in THF (80 mL) and $H_2O$ (8 mL) was added LiOH (288 mg, 12 mmol, 1.5 equiv.), the mixture was stirred in 0 ºC for 4 hours. The reaction was quenched with $NaHCO_3$ saturated solution and extracted with EtOAc (3 × 10 mL). The combined organic layers were washed with brine, dried over $Na_2SO_4$, filtered and concentrated in vacuo. The residue was dissolved in 16 ml DMF to obtain 0.2 M **S10** solution, and used in peptides synthesis without purification.

3.3 List of synthetic peptides used in this study

| Peptide | Calcd. MW | Obs. MW |
|---|---|---|
| **P1**: ARTKQTARKSTG | 1302.8 | 1302.2 |
| **P2**: ARTKWTARKSTG | 1360.7 | 1361.1 |
| **P3**: ARTKYTARKSTG | 1337.8 | 1338.2 |
| **P4**: ARTKHTARKSTG | 1311.8 | 1312.5 |
| **P5**: ARTKQserTARKSTG | 1461.8 | 1462.4 |
| **P6**: ARTKQ(5-F-Try)TARKSTG | 1464.8 | 1464.3 |
| **P7**: ARTKQ(5-MT)TARKSTG | 1475.8 | 1476.4 |
| **P8**: ARTKQdopTARKSTG | 1438.8 | 1439.4 |
| **P9**: ARTKQtyrTARKSTG | 1422.8 | 1423.4 |
| **P10**: ARTKQ(3-MethoxyTyr)TARKSTG | 1452.8 | 1453.3 |
| **P11**: ARTKQ(MeTyr)TARKSTG | 1436.8 | 1437.3 |
| **P12**: ARTKQhisTARKSTG | 1396.8 | 1397.2 |
| **P13**: ARTKQdaqTARKSTG | 1437.3 | 1436.8 |

3.5 Synthesis of AcNH-AWA-CONH2 and AcNH-AQserA-CONH2 peptides

Following the Fmoc SPPS protocol, before cleavage, replace the Fmoc protecting group with an Ac group to obtain AcNH-AWA-CONH2 and AcNH-AQserA-CONH2.

3.5.1 AcNH-AWA-CONH2

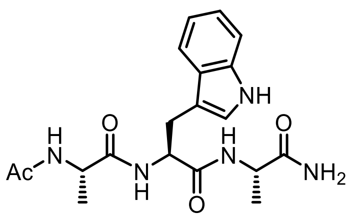

AcNH-AWA-CONH2

$^1$H NMR (600 MHz, $D_2O$) δ 7.65 – 7.62 (m, 1H), 7.52 – 7.49 (m, 1H), 7.27 – 7.23 (m, 2H), 7.18 (ddd, *J* = 7.9, 7.0, 1.0 Hz, 1H), 4.61 (t, *J* = 6.9 Hz, 1H), 4.17 (qd, *J* = 7.2, 1.3 Hz, 2H), 3.34 – 3.24 (m, 2H), 1.89 (s, 3H), 1.26 (d, *J* = 7.3 Hz, 3H), 1.18 (d, *J* = 7.2 Hz, 3H).

$^{13}$C NMR (150 MHz, $D_2O$) δ 177.2, 175.2, 174.3, 173.2, 136.1, 126.8, 124.5, 122.1, 119.5, 118.2, 111.9, 108.4, 54.3, 49.9, 49.3, 26.4, 21.4, 16.4, 16.0.

3.5.2 AcNH-AQserA-CONH2

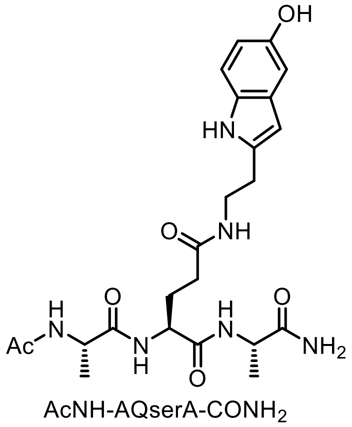

AcNH-AQserA-CONH2

$^1$H NMR (600 MHz, $D_2O$) δ 7.36 (d, *J* = 8.6 Hz, 1H), 7.18 (s, 1H), 7.07 (d, *J* = 2.4 Hz, 1H), 6.82 (dd, *J* = 8.7, 2.4 Hz, 1H), 4.25 (q, *J* = 7.2 Hz, 1H), 4.20 (q, *J* = 7.2 Hz, 1H), 4.16 (dd, *J* = 9.1, 5.4 Hz, 1H), 3.47 (td, *J* = 6.7, 1.9 Hz, 2H), 2.90 (t, *J* = 6.7 Hz, 2H), 2.23 (t, *J* = 7.4 Hz, 2H), 2.10– 1.93(m, 1H), 1.98 (s, 3H), 1.80 (ddt, *J* = 14.0, 9.1, 7.1

Hz, 1H), 1.36 (d, $J$ = 7.2 Hz, 3H), 1.34 (d, $J$ = 7.2 Hz, 3H).

$^{13}C$ NMR (150 MHz, $D_2O$) δ 177.5, 175.4, 174.6, 174.2, 172.8, 148.5, 131.4, 127.7, 124.5, 112.6, 111.5, 111.3, 102.7, 52.8, 49.8, 49.4, 40.0, 31.8, 26.8, 24.1, 21.5, 16.6, 16.4

#### 4. Reactions between peptides and AlkTAD

##### 4.1 Reaction between AcNH-AWA-$CONH_2$ and AlkTAD

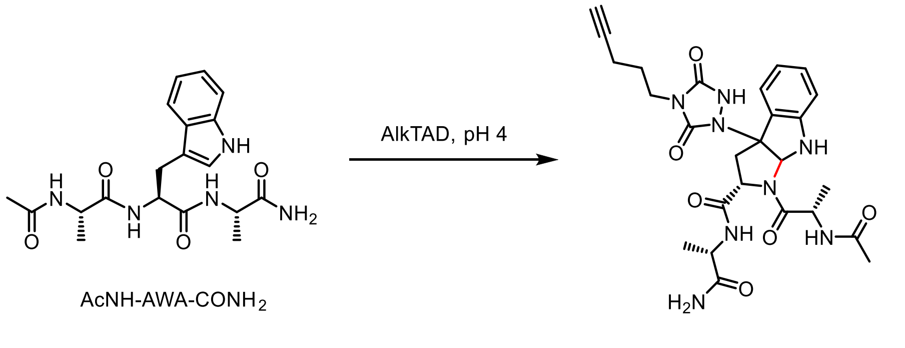

1

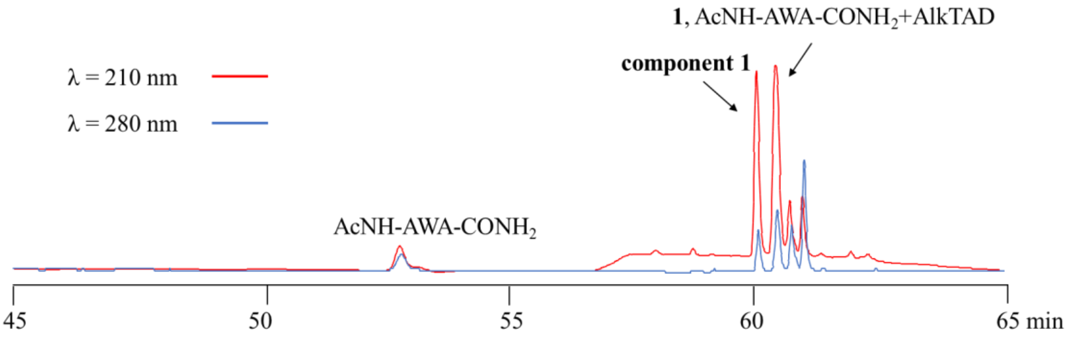

**Figure S4.1** LC chromatogram (red: 210 nm, blue: 280 nm) of the reaction of AcNH-AWA-$CONH_2$ with AlkTAD was conducted at pH 4.

To a solution of AcNH-AWA-$CONH_2$ (7.7 mg, 0.02 mmol, 1.0 equiv) obtained above in buffer (citrate-phosphate buffer, 5.0 mL, pH 4) and $CH_3CN$ (5.0 mL) was added AlkTAD (4.0 mg, 0.024 mmol, 0.006 M in $CH_3CN$, 1.0 mg/mL, 1.2 equiv) and the mixture was stirred at 0 °C for 2 h. The mixture was evaporated under reduced pressure to less than 5 ml, and purified by HPLC (XSelect Peptide CSH C18 OBD Prep Column, eluent A water, eluent B $CH_3CN$ with 0.1% formic acid; gradient T = 0 min: 0% B, T =

5 min: 0% B, T = 40 min: 35% B, T = 55 min: 38% B, 4.0 mL/min; retention time: 40 min, 42 min) to give two Components. **Component 1** was obtained as a 1:1.8 mixture and corresponded to the desired molecular weight. (For specific NMR spectra, referred to the NMR spectrum of **Component 1**. For mass spectra, referred to the MS spectrum of **Component 1**); the second one was compound **1**(5.1 mg, 46%).

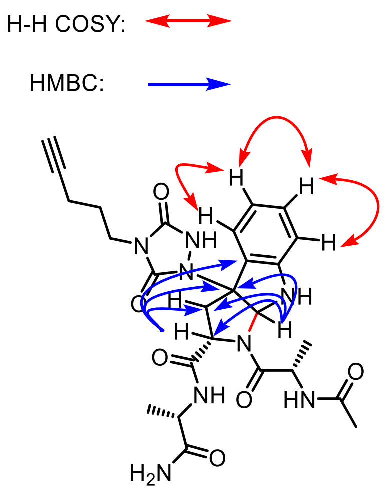

**1**

$^1$H NMR (700 MHz, $D_2O$) δ 7.34 ( d, *J* = 7.0 Hz, 1H), 7.26 (t, *J* = 7.7 Hz, 1H), 6.90 (t, *J* = 7.5 Hz, 1H), 6.81 – 6.79 (m, 1H), 6.20 (s, 1H), 4.66 – 4.59 (m, 2H), 4.19 (q, *J* = 7.2 Hz, 1H), 4.14 (dd, *J* = 9.7, 6.7 Hz, 1H), 3.49 (t, *J* = 6.4 Hz, 2H), 3.02 (dd, *J* = 13.1, 6.7 Hz, 1H), 2.82 (dd, *J* = 13.1, 9.8 Hz, 1H), 2.16 (t, *J* = 2.7 Hz, 1H), 2.01 (tdd, *J* = 6.7, 4.3, 2.8 Hz, 2H), 1.95 (s, 3H), 1.71 – 1.65(m, 2H), 1.36 (d, *J* = 7.0 Hz, 3H), 1.30 (d, *J* = 7.2 Hz, 3H).

$^{13}$C NMR (175 MHz, $D_2O$) δ 177.6, 174.8, 174.2, 171.4, 156.6, 156.2, 149.5, 131.7, 125.3, 124.0, 121.2, 111.8, 111.8, 84.2, 79.6, 76.6, 69.4, 60.0, 49.5, 48.0, 38.7, 37.7, 25.5, 21.2, 16.6, 16.1, 15.1.

#### 4.2 Reaction between AcNH-AQserA-CONH$_2$ and AlkTAD

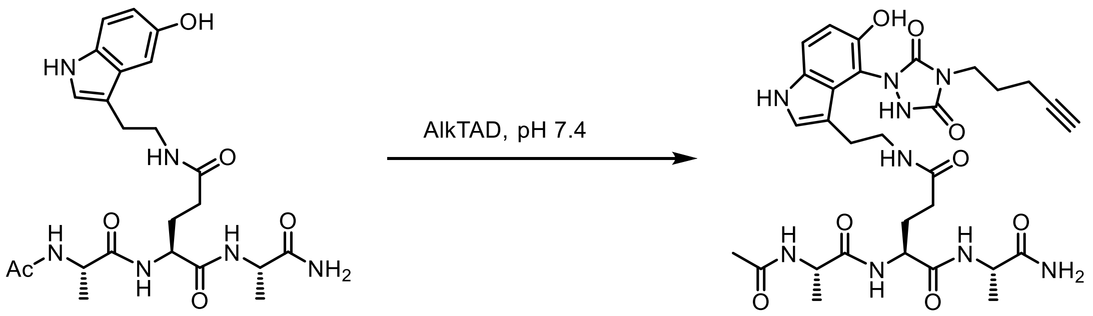

AcNH-AQserA-CONH$_2$

**2**

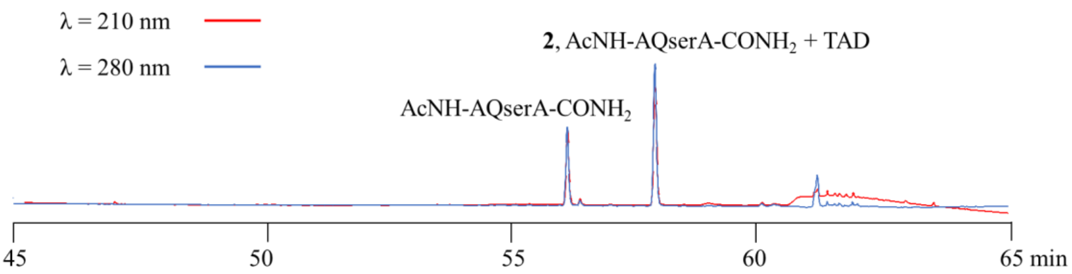

**Figure S4.2** LC chromatogram (red: 210 nm, blue: 280 nm) of the reaction of AcNH-AWA-$CONH_2$ with AlkTAD was conducted at pH 4.

To a solution of AcNH-AQserA-$CONH_2$ (8.9 mg, 0.02 mmol, 1.0 equiv) obtained above in citrate-phosphate buffer (5.0 mL, pH 7.4) and $CH_3CN$ (5.0 mL) was added AlkTAD (4.0 mg, 0.024 mmol, 0.006 M in $CH_3CN$, 1.0 mg/mL, 1.2 equiv) and the mixture was stirred at 0 °C for 2 h. The mixture was evaporated under reduced pressure to less than 5 ml, and purified by HPLC (XSelect Peptide CSH C18 OBD Prep Column, eluent A water, eluent B $CH_3CN$ with 0.1% formic acid; gradient T = 0 min: 0% B, T = 5 min: 0% B, T = 40 min: 35% B, 4.0 mL/min; retention time: 24 min) to give the product **2** (10.2 mg, 78%).

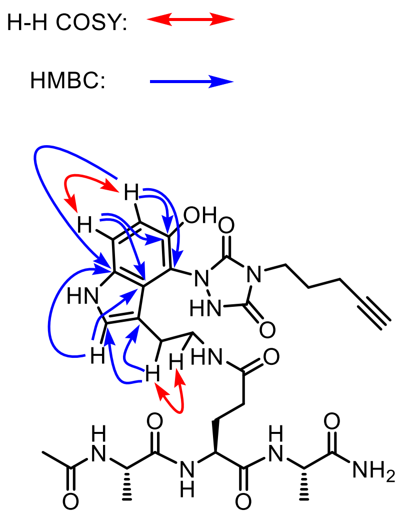

**2**

$^1$H NMR (700 MHz, $D_2O$) δ 7.51 (d, $J$ = 8.8, 1H), 7.25 (s, 1H), 6.93 (d, $J$ = 8.8, 1H), 4.26 (qd, $J$ = 7.3, 1.7 Hz, 1H), 4.24 – 4.18 (m, 2H), 3.80 (td, $J$ = 6.7, 1.7 Hz, 2H), 3.52 – 3.43 (m, 1H), 3.35 – 3.32 (m, 1H), 2.89 – 2.75 (m, 2H), 2.41 (t, $J$ = 2.6 Hz, 1H), 2.35 (td, $J$ = 6.9, 2.7 Hz, 2H), 2.31 – 3.27 (m, 2H), 2.03 – 1.93 (m, 2H), 1.99 (d, $J$ = 7.6, 3H )1.92 – 1.83 (m, 1H), 1.37 (dd, $J$ = 9.2, 7.3 Hz, 3H), 1.34 (dd, $J$ = 15.6, 7.3 Hz, 3H).

$^{13}$C NMR (175 MHz, $D_2O$) δ 177.5, 177.5, 175.4, 174.6, 174.6, 174.2, 174.2, 172.8, 154.1, 154.0, 147.6, 132.1, 126.3, 126.3, 124.9, 115.5, 111.8, 110.3, 109.9, 109.8, 84.2, 69.8, 52.8, 52.7, 49.8, 49.4, 49.4, 39.9, 38.7, 31.8, 31.7, 26.9, 26.9, 26.2, 24.7, 21.5,

21.5, 16.7, 16.7, 16.4, 16.4, 15.2.

4.3 Reaction between AcNH-AWA-$CONH_2$/AcNH-AQserA-$CONH_2$ with AlkTAD at different pH values

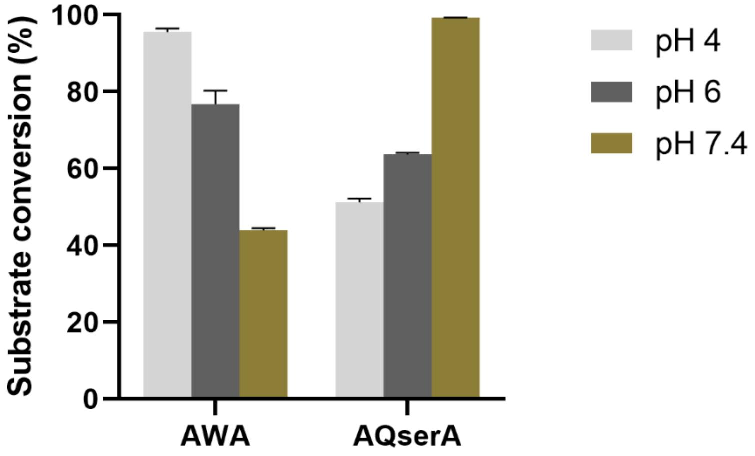

**Figure S4.3** The conversion of the peptides: at pH 4, 6, and 7.4, peptides reacted with 1.5 equivalents of AlkTAD, and the reaction was quenched after 5 minutes using trans,trans-2,4-hexadien-1-ol.[3] The error bars represent the standard deviation from three different experiments.

4.4 General procedure for reactions between peptides and AlkTAD

The purified peptides were dissolved in milliQ $H_2O$ to make stock solutions of 3 mM. 10 mM AlkTAD solution is prepared in MeCN. 7 µL (21 nmol, 1.0 equiv.) of the peptides stock solution is added to 90 µL buffer solution (citrate-phosphate buffer) with a known pH value under ice bath. To this mixture 3 µL of the 10 mM AlkTAD solution (30 nmol, 1.5 equiv.) is added mixed immediately by vortex, after standing in ice bath for 5 minutes, the reaction is quenched by 3 µL of the 100 mM *trans,trans*-2,4-hexadien-1-ol solution in MeCN.[3] The resulting reaction mixture is analyzed via LC-MS.

4.5 AlkTAD-peptide reactions at different pH values

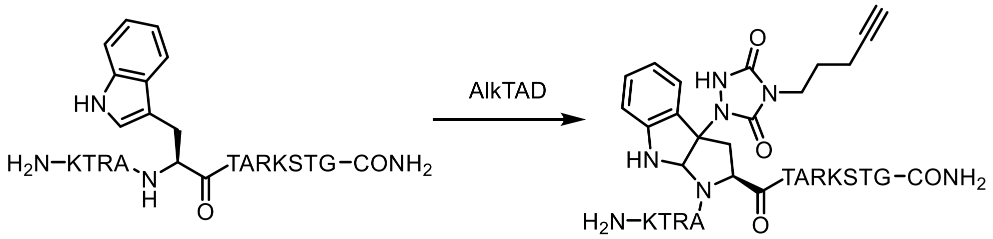

**H3W5** **H3W5 + AlkTAD**

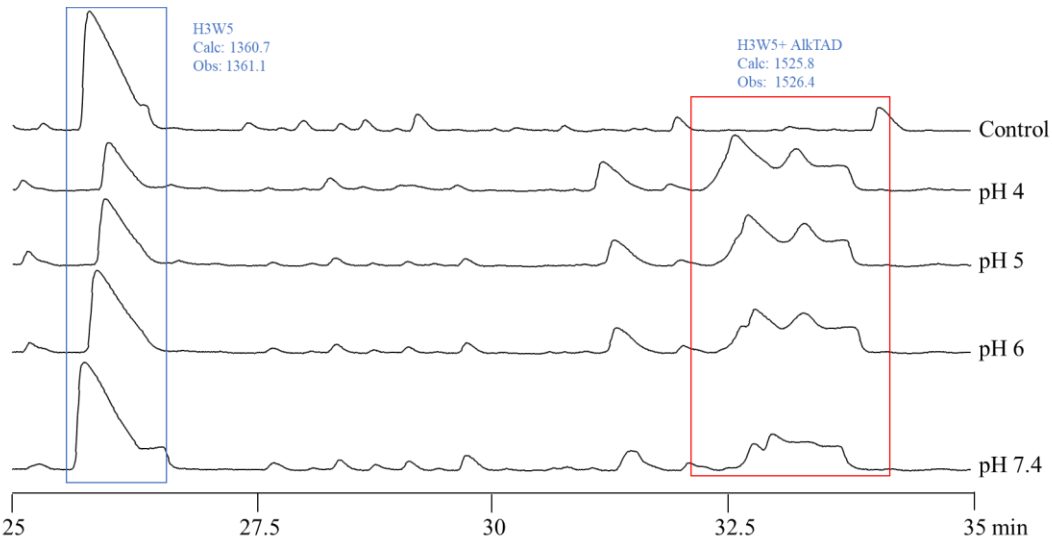

**Figure S4.5.1.** LC chromatogram (210 nm) of the reaction between **P2** (H3W5) and AlkTAD at different pH levels. The conversion rate of **P2** decreased significantly with the increase of pH.

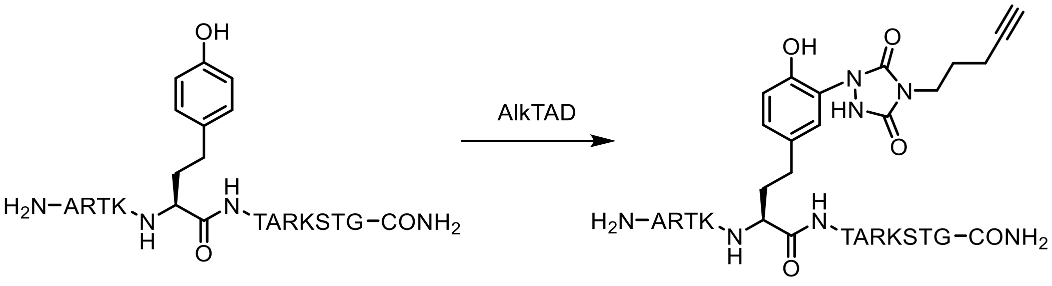

**H3Y5** **H3Y5 + AlkTAD**

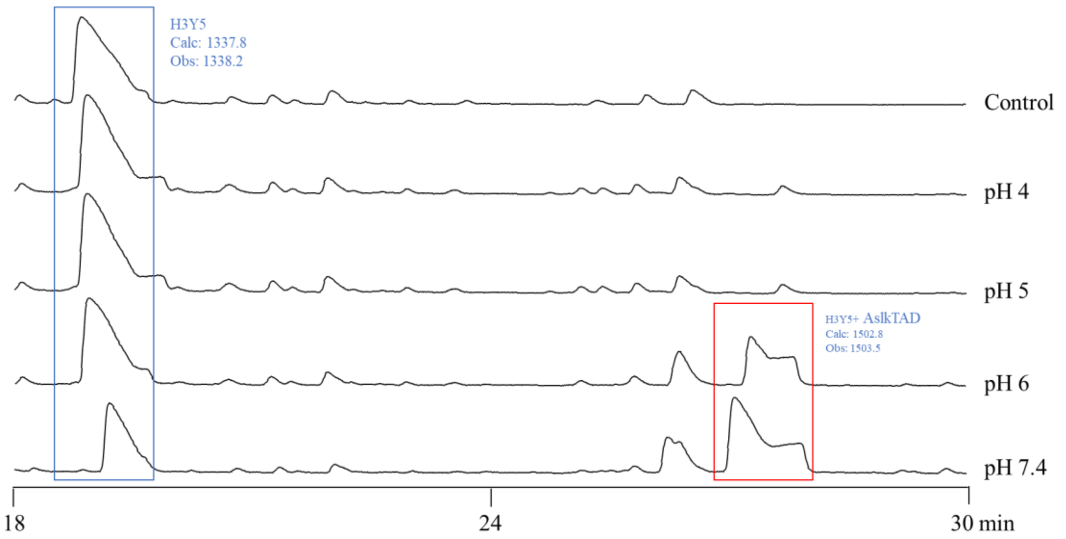

**Figure S4.5.2.** LC chromatogram (210 nm) of the reaction between **P3** (H3Y5) and AlkTAD at different pH levels. The conversion rate of **P3** with TAD varied with pH, showing increased AlkTAD modification at higher pH levels. Notably, significant reaction occurred at pH above

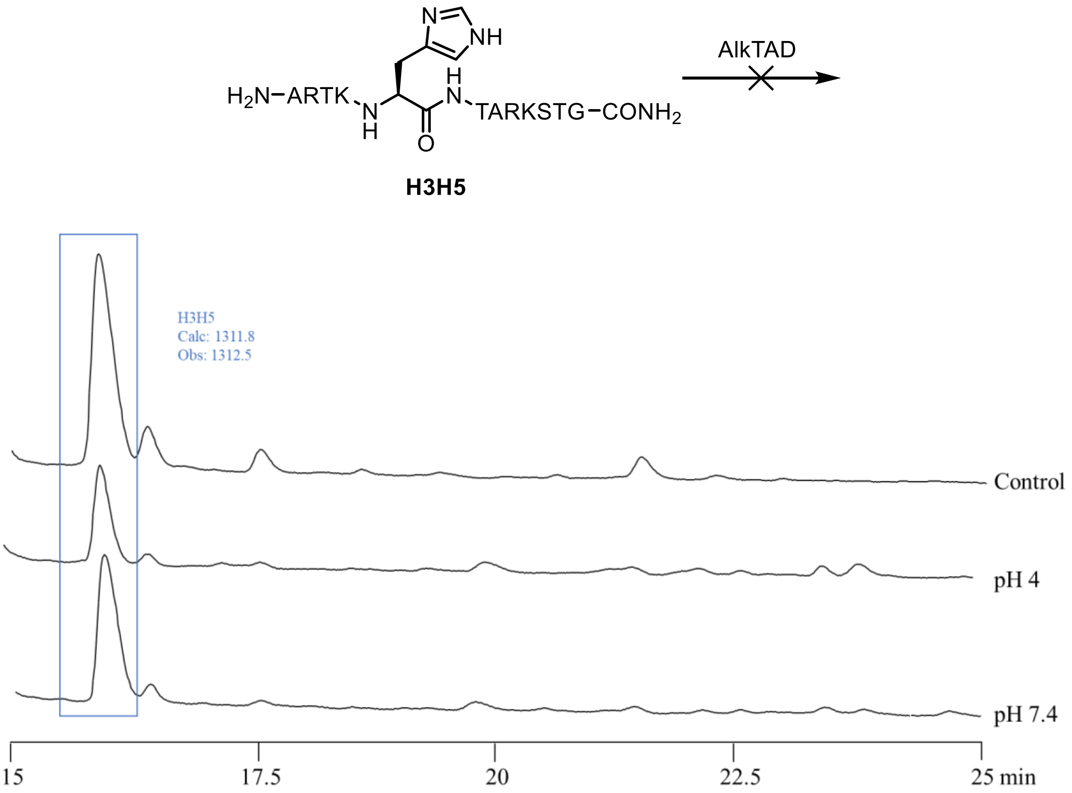

**Figure S4.5.3.** LC chromatogram (210 nm) of the reaction between **P4** (H3H5) and AlkTAD at different pH levels. **P4** did not react with AlkTAD at pH 4 - 7.4.

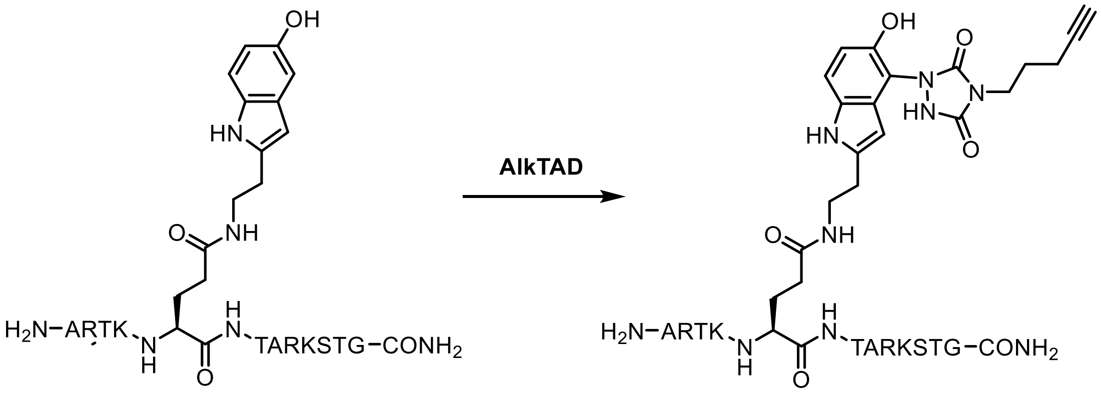

**H3Q5ser** **H3Q5ser + AlkTAD**

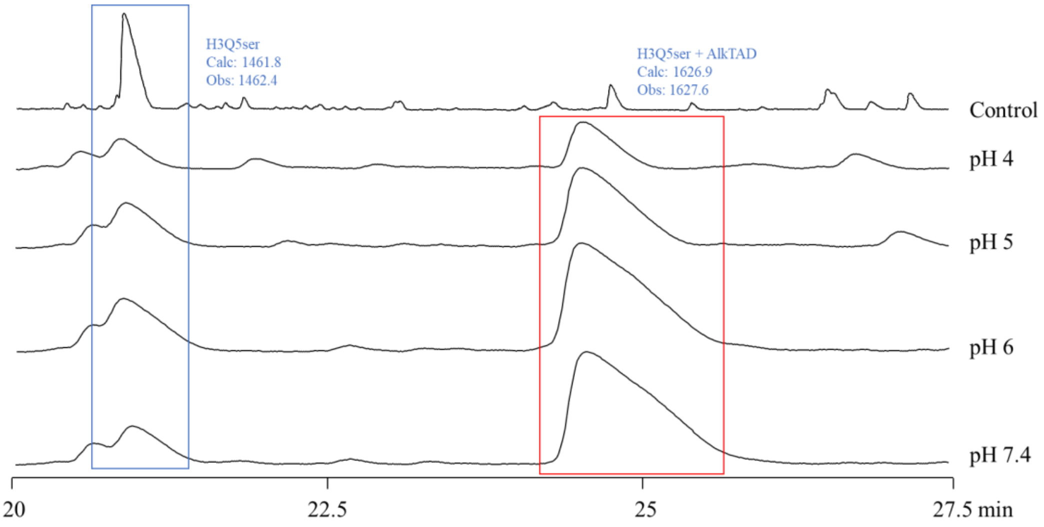

**Figure S4.5.4.** LC chromatogram (210 nm) of the reaction between **P5** (H3Q5ser) and AlkTAD at different pH levels. The conversion rate of peptide **P5** (H3Q5ser) with TAD varied slightly with pH, showing increased TAD modification at higher pH levels. Notably, the conversion rate exceeded 60% even at pH 4.

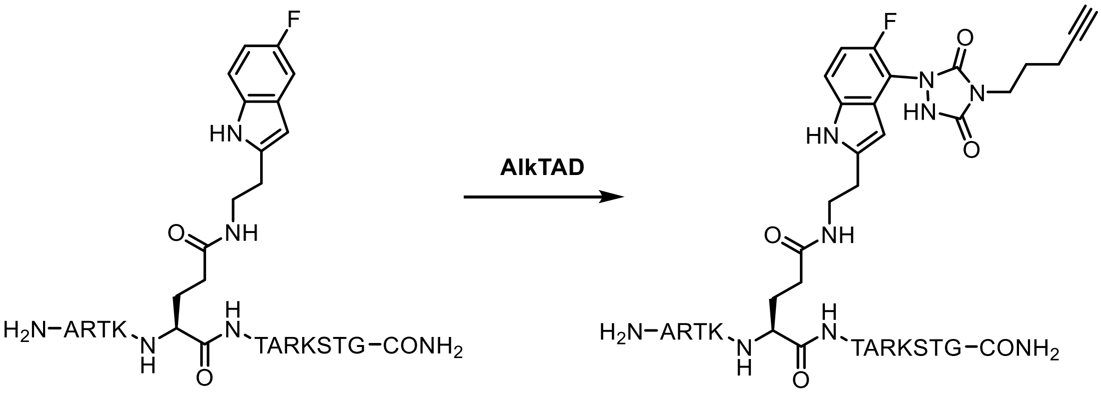

**H3Q5(5F-Try)** **H3Q5(5F-Try) + AlkTAD**

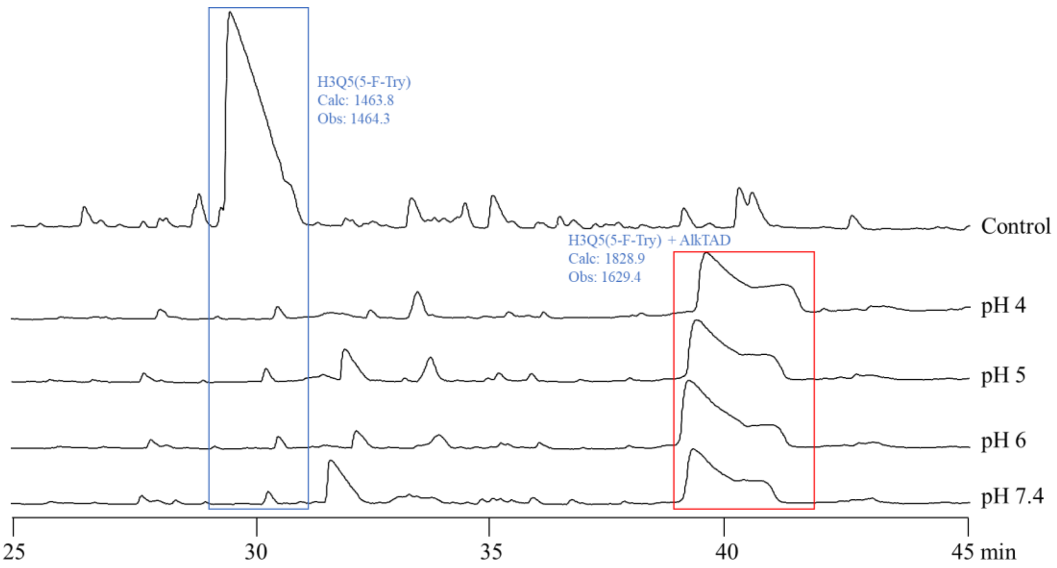

**Figure S4.5.5.** LC chromatogram (210 nm) of the reaction between **P6** (H3Q5(5F-Try)) and AlkTAD at different pH levels. The conversion rate of **P6** with AlkTAD was unaffected by the tested pH levels, with complete reaction observed at pH 4 or above.

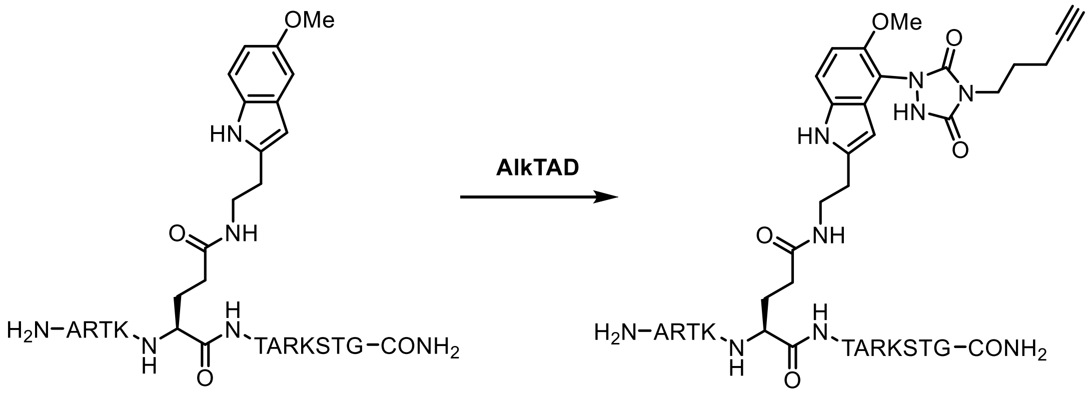

**H3Q5(5-MT)** **H3Q5(5-MT) + AlkTAD**

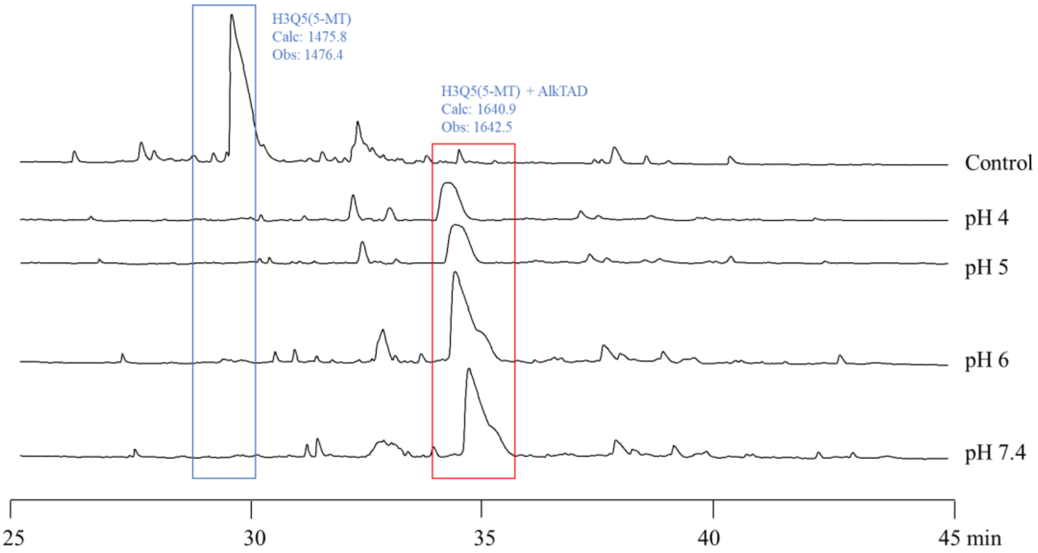

**Figure S4.5.6.** LC chromatogram (210 nm) of the reaction between **P7** (H3Q5(5-MT)) and

AlkTAD at different pH levels. Conversion rate of peptide **P7** with AlkTAD showing minimal changed across different pH levels. Complete reaction was observed at pH 4.

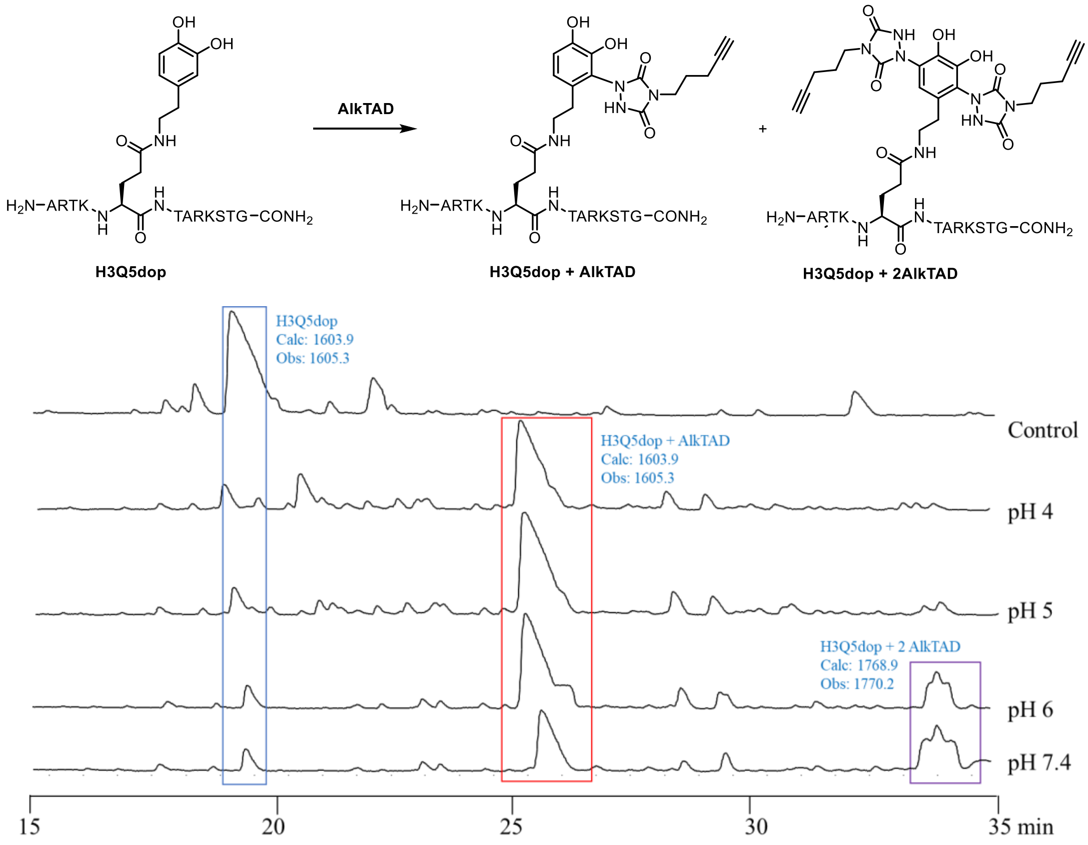

**Figure S4.5.7.** LC chromatogram (210 nm) of the reaction between **P8**(H3Q5dop) and AlkTAD at different pH levels. The conversion rate of peptide **P8** with TAD showed minimal change across pH levels, with complete reaction observed at pH 4. At higher pH, the AalTAD-modified peptide continued to react, leading to a double AlkTAD modification ($t_R$ = 33–34 min).

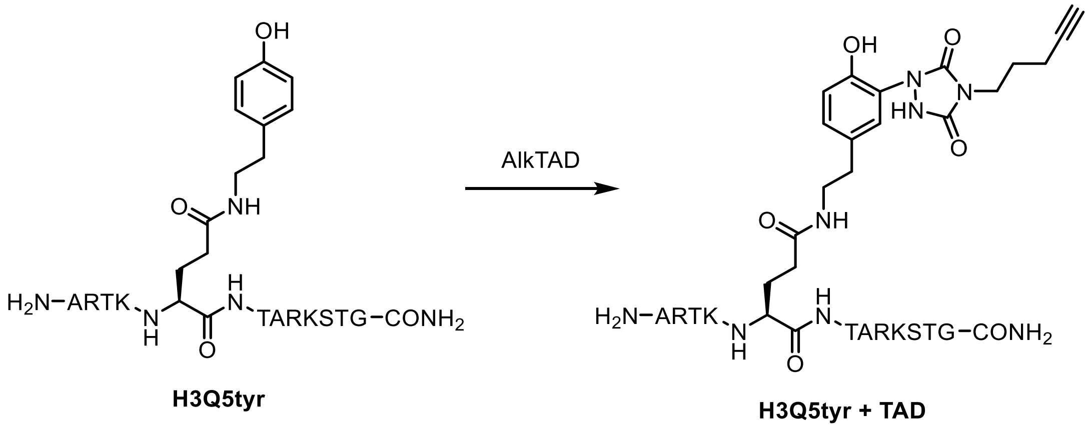

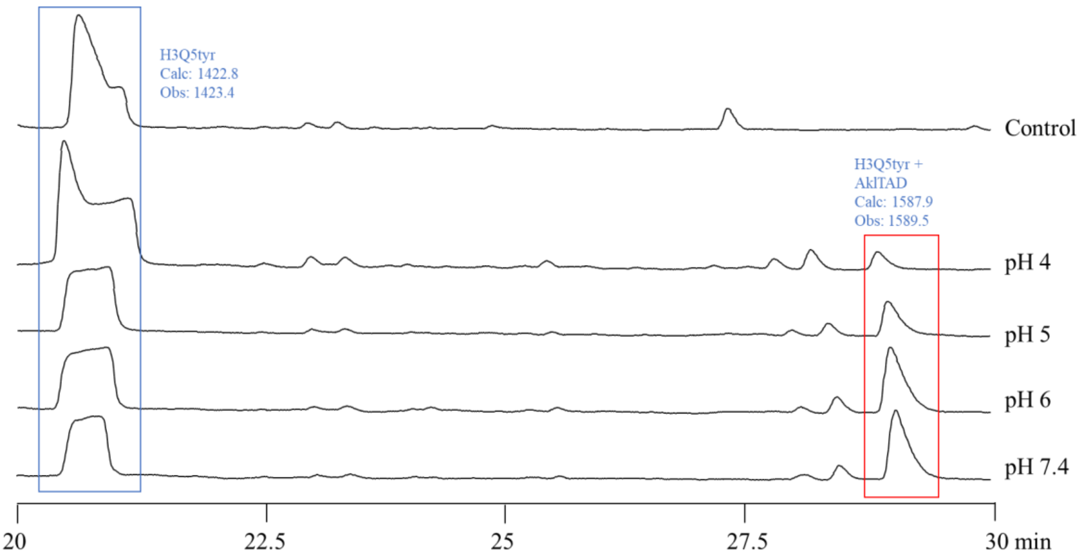

**Figure S4.5.8.** LC chromatogram (210 nm) of the reaction between **P9** (H3Q5tyr) and AlkTAD at different pH levels. The conversion rate of peptide **P9** with AlkTAD was strongly affected by pH, showing increased AlkTAD modification at higher pH levels.

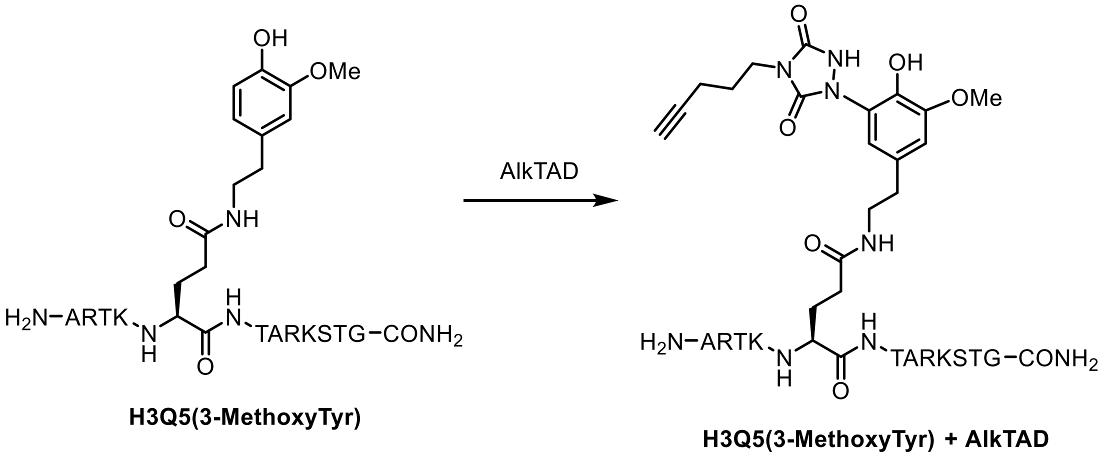

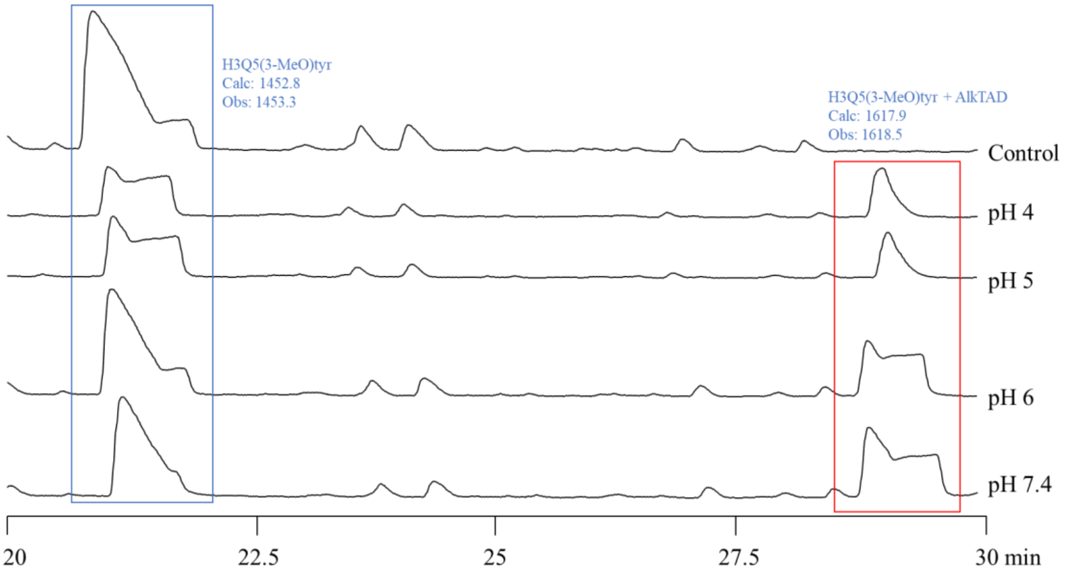

**Figure S4.5.9.** LC chromatogram (210 nm) of the reaction between **P10** (H3Q5(3-MethoxyTyr)) and AlkTAD at different pH levels. At higher pH levels, increased amounts of TAD-modified peptide were observed

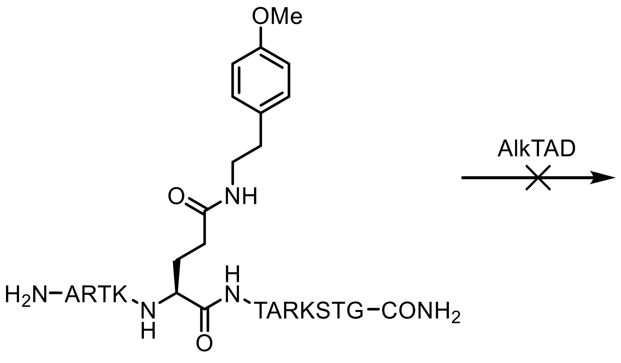

**H3Q5metyr**

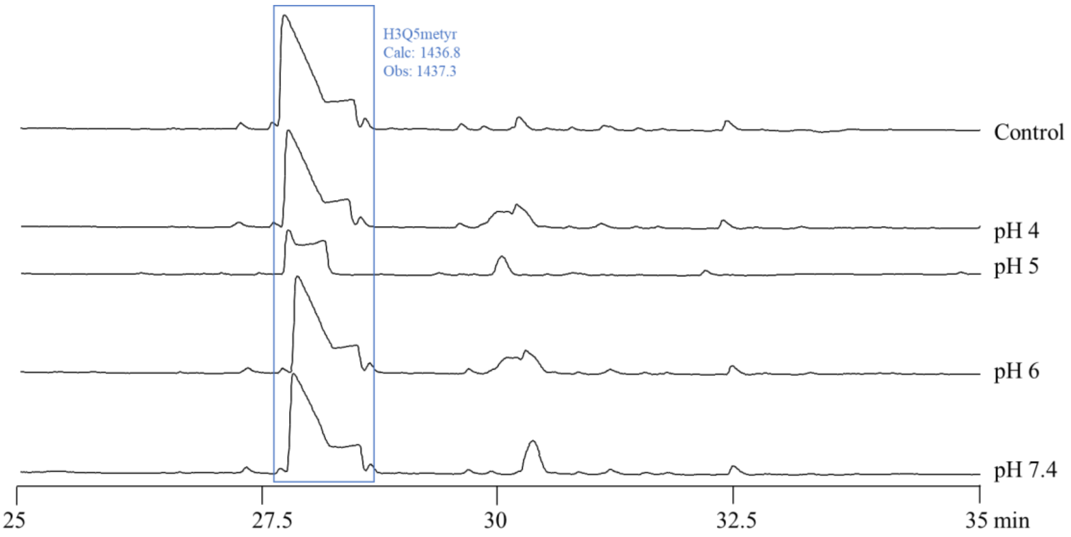

**Figure S4.5.10.** LC chromatogram (210 nm) of the reaction between **P11** (H3Q5metyr) and AlkTAD at different pH levels. **P11** did not react with TAD at pH 4 - 7.4.

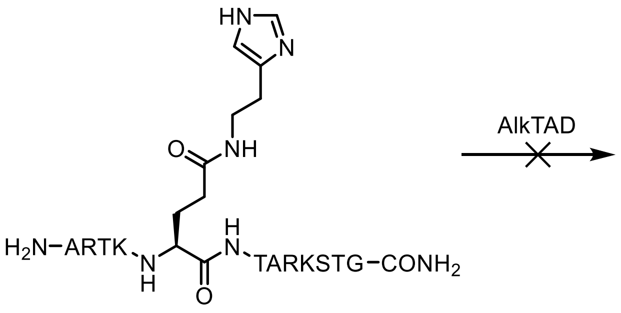

**H3Q5his**

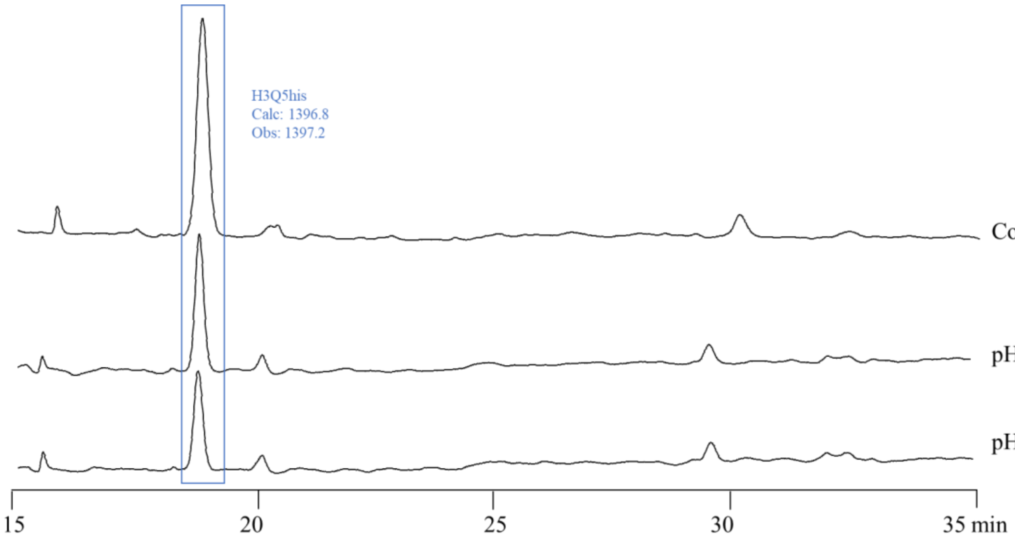

**Figure S4.5.11.** LC chromatogram (210 nm) of the reaction between **P12** (H3Q5his) and TAD at different pH levels. The **P12** did not react with AlkTAD at pH 4-7.4.

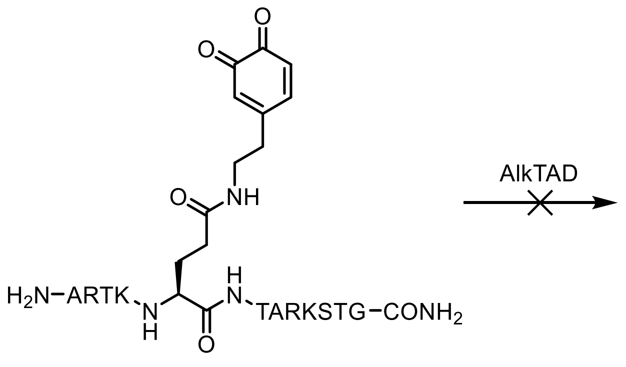

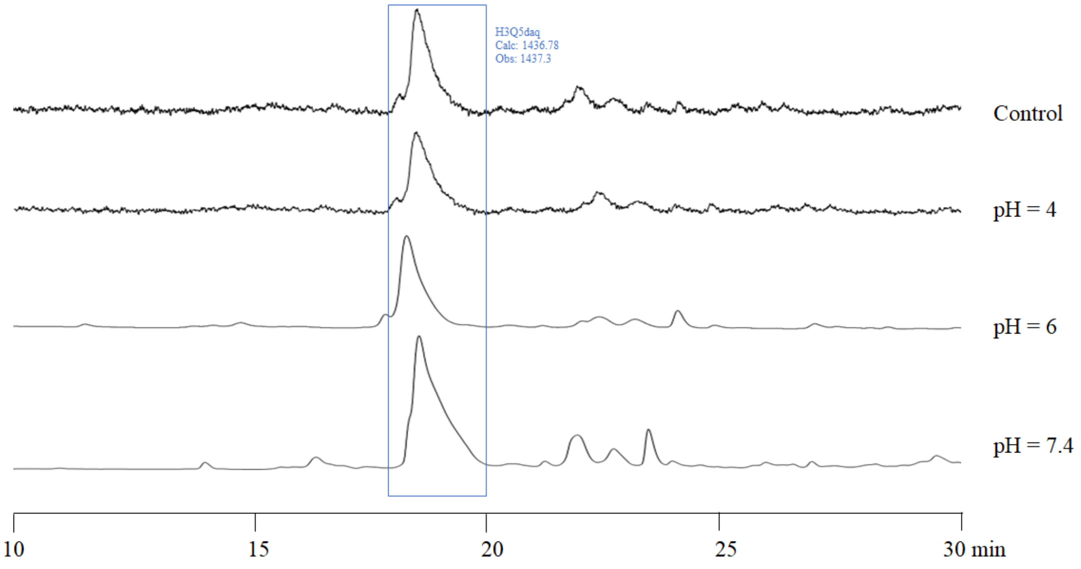

**Figure S4.5.12.** LC chromatogram (210 nm) of the reaction between **P13** (H3Q5daq) and TAD at different pH levels. Notably, **P13** could not react with AlkTAD at pH 4-7.4.

4.6 The competition reaction between two peptide substrates and AlkTAD

7 µL (21 nmmol, 1.0 equiv.) of the peptides stock solution is mixture with 7 µL (21 nmmol, 1.0 equiv.) of peptide **P5** (H3Q5ser), added to 84 µL pH 4 buffer solution (citrate-phosphate buffer) under ice bath. To this mixture 2 µL of the 10 mM AlkTAD solution (20 nmmol, 1.0 equiv.) is added mixed immediately by vortex, after standing in ice bath for 5 minutes, the reaction is quenched by 2 µL of the 100 mM *trans,trans*-2,4-Hexadien-1-ol solution in MeCN. The resulting reaction mixture is analyzed via LC-MS.

4.6.1 H3Q5ser and H3W5

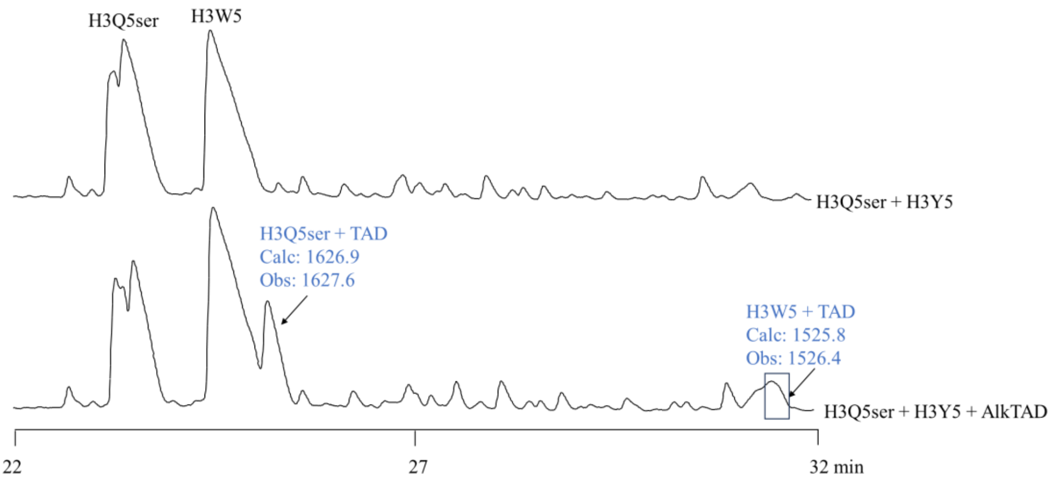

**Figure S4.6.1** LC chromatogram (210 nm) of the competitive reaction of **P5** (H3Q5ser) with **P2** (H3W5) was conducted at pH 4.

4.6.2 H3Q5ser and H3Y5

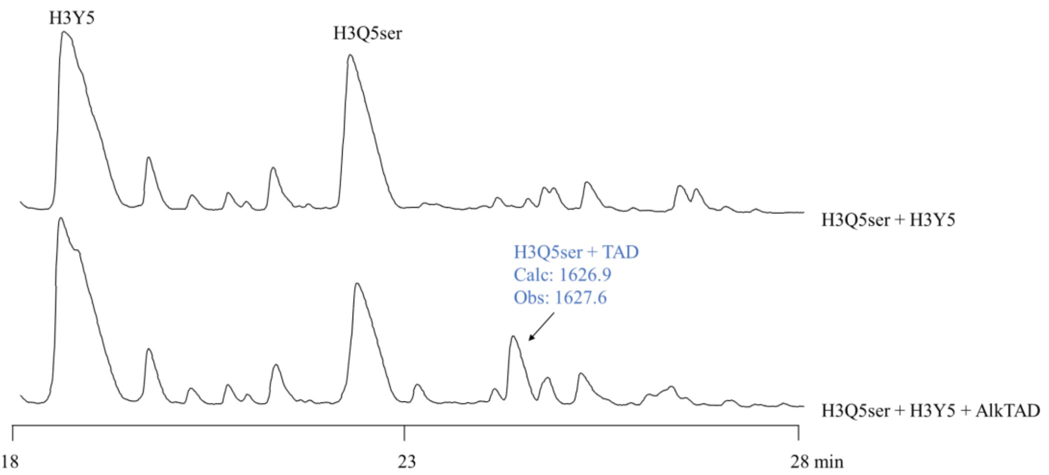

**Figure S4.6.2** LC chromatogram (210 nm) of the competitive reaction of **P5** (H3Q5ser) with **P3** (H3Y5) was conducted at pH 4.

#### 4.6.3 H3Q5ser and H3Q5dop

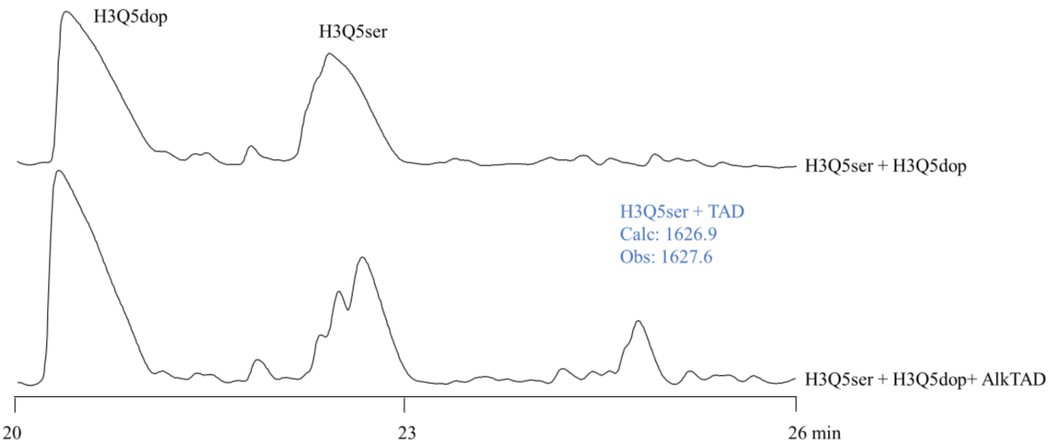

**Figure S4.6.3** LC chromatogram (210 nm) of the competitive reaction of **P5** (H3Q5ser) with **P8** (H3Q5dop) was conducted at pH 4.

#### 4.6.4 H3Q5ser and H3Q5tyr

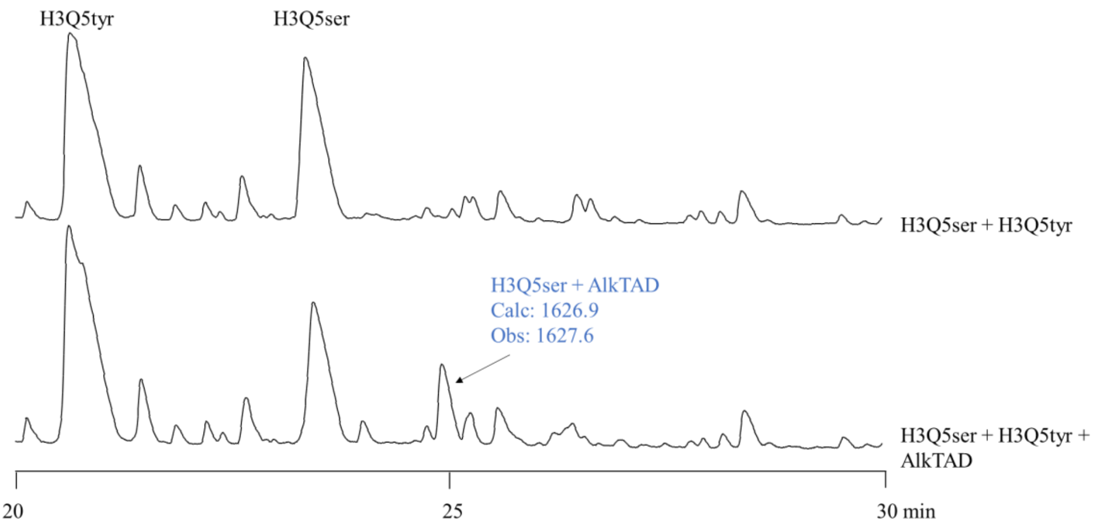

**Figure S4.6.4** LC chromatogram (210 nm) of the competitive reaction of **P5** (H3Q5ser) with **P9** (H3Q5tyr) was conducted at pH 4.

4.6.5 H3Q5ser and H3Q5daq

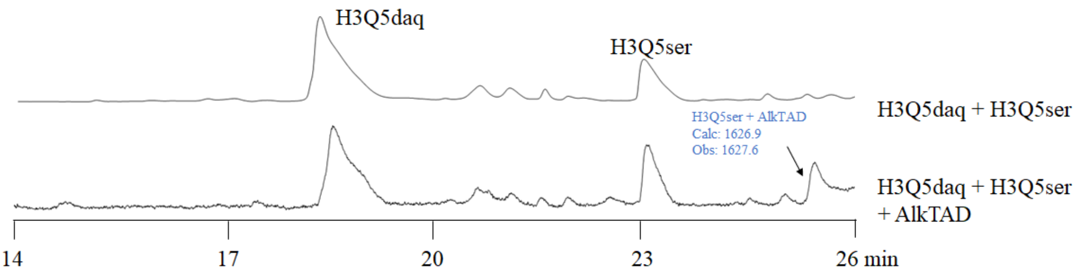

**Figure S4.6.4** LC chromatogram (210 nm) of the competitive reaction of **P5** (H3Q5ser) with **P13** (H3Q5daq) was conducted at pH 4.

#### 5. *In vitro* labeling assays

For nucleosome core particle (NCP) labeling assays, 1 µM NCPs were treated with 0.1 µM TGM2 in the buffer (pH 7.5) containing 50 mM Tris-HCl, 5 mM $CaCl_2$, and 2 mM DTT (freshly added) at 37 °C in the presence of corresponding monoamaines (0.5 µM) for 2 hours.[1,2] The buffer exchange for the further labeling assays was performed using 0.5 mL Centrifugal Filter (3K, Millipore) with a 120-fold v/v for the removal of excess monoamine from the old reaction buffer systems. Specifically, the modified NCPs were dissolved in McIlvaine buffer (pH 4) via buffer exchange and then incubated with freshly prepared AlkTAD (10 µM) on ice for 5 min, followed by the addition of trans,trans-2,4-hexadien-1-ol (10 µM) to quench the unreacted TAD. For the Cy5 labeling of AlkTAD-modified NCPs, 0.4 % SDS was added to the NCP solution to enhance the yield of CuAAC. 50 μL of NCP solution was added to a premixed solution containing 3 μL of 10 mM Cy5-azide (Sigma-Aldrich, 777323), 10 μL of a 3:7 mixture of 50 mM $CuSO_4$ and 100 mM THPTA, and then vortexed. Thereafter, 5 μL of 100 mM freshly made TCEP was added to initiate the click reaction followed by incubation (1-2 hours) at 30 °C. Then, 10 μL of 0.5 M EDTA was added to quench the reactions. Excess reagents were removed by MeOH/$CHCl_3$ protein precipitation or concentration-dilution using a 0.5 mL Centrifugal Filter (3K, Millipore). Pellets were washed by 500 μL MeOH/H2O (9:1) prior to a second centrifugation. The labeled NCPs were analyzed by sodium dodecyl sulfate polyacrylamide gel electrophoresis (SDS-PAGE) followed by western blot analysis using IRDye 680RD Streptavidin (LI-COR).The air-dried samples were then analyzed by SDS-PAGE followed by in-gel imaging using Odyssey CLx Imaging System (wavelength 680 nm). H3 was used as the loading control in SDS-PAGE and western blot analyses.

For labeling the histones extracted from cell or tissue samples, acid-extracted histones were desalinated, lyophilized, and then resuspended in McIlvaine buffer (pH 4). The freshly dissolved histones were first treated with 0.1 mM $K_3[Fe(CN)_6]$ and then incubated with freshly prepared AlkTAD (10 μM) on ice for 5 min, followed by the addition of trans,trans-2,4-hexadien-1-ol (10 μM) to quench the unreacted TAD. 50 μL of the modified histones containing 0.4% SDS was added to a premixed solution containing 3 μL of 10 mM Cy5-azide (Sigma-Aldrich, 777323), 10 μL of a 3:7 mixture of 50 mM $CuSO_4$ and 100 mM THPTA, and then vortexed. Thereafter, 5 μL of 100 mM freshly made TCEP was added to initiate the click reaction followed by incubation (1-2 hours) at 30 °C. Then, 10 μL of 0.5 M EDTA was added to quench the reactions. Excess reagents were removed by MeOH/$CHCl_3$ protein precipitation or concentration-dilution using a 0.5 mL Centrifugal Filter (3K, Millipore). Pellets were washed by 500 μL MeOH/H2O (9:1) prior to a second centrifugation. The air-dried samples were then analyzed by SDS-PAGE followed by in-gel imaging using Odyssey CLx Imaging System (wavelength 680 nm). H3 was used as the loading control in SDS-PAGE and western blot analyses.

#### 6. Expression of TGM2 in HEK 293T cells

The plasmid of TGM2 and its mutant were constructed expressed in HEK 293T cells based on the protocol of our previous research.[1,2] Briefly, wild-type TGM2 and the TGM2-C277A mutant were overexpressed in HEK 293T cells using Lipofectamine 2000 Transfection Reagent (Thermo Fisher Scientific) according to the manufacturer's protocol. HEK 293T cells (ATCC) were cultured at 37 °C with 5% CO2 in DMEM medium supplemented with 10% fetal bovine serum (FBS) (Sigma-Aldrich), 2 mM L-glutamine and 500 units $mL^{-1}$ penicillin and streptomycin. The cells were stimulated with 2 μM calcium ionophore (Sigma-Aldrich, A23187) for 6 h at 37 °C before lysis in DPBS buffer (Gibco), and then the expression of TGM2 was detected by western blot analyses with anti-TGM2 antibody (CST, #3557).

#### 7. Expression of HA-tagged WT H3 and H3-Q5E in HTC116 cells

The pCMV-HA-H3 plasmid was constructed and used in our previous research.[4] The pCMV-HA-H3-Q5E mutant plasmid was constructed by site-directed mutagenesis using 5′-GCTCGTACTAAGGAAACCGCTCGCAAG-3′ and 5′-CTTGCGAGCGGTTTCCTTAGTACGAGC-3′ as primers. The HA-tagged WT H3 and H3-Q5E were expressed in HTC116 cells using Lipofectamine 2000 Transfection Reagent (Thermo Fisher Scientific) according to the manufacturer's protocol. HA-H3 overexpression was detected by western blot analysis with anti-H3 and anti-HA antibodies.

#### 8. Extraction of histones from cultured cells

The extraction of histones from cells was performed according to the previously described high salt extraction method.[1,2] Briefly, the cell lysis solution was prepared using extraction buffer (10 mM HEPES pH 7.9, 10 mM KCl, 1.5 mM MgCl2, 0.34 M sucrose, 10% glycerol, 0.2% NP40, protease and phosphatase inhibitors to 1 × fromstock). After spinning down, the pellet was extracted using a no-salt buffer (3 mM EDTA, 0.2 mM EGTA). After discarding the supernatant, the final pellet was extracted by using high-salt buffer (50 mM Tris pH 8.0, 2.5 M NaCl, 0.05% NP40) in 4 °C cold room for 1 hour. After spinning down, the supernatant containing extracted histones was collected, desalted, and lyophilize for further labeling assays and analyses.

#### 9. Cell fractionation

Cytosolic and nuclear fractions were prepared using NEPER Nuclear and Cytoplasmic Extraction Reagents (Thermo Scientific) according to the manufacturer's protocol. Histones were extracted from the pellet using high salt extraction protocol, as described above (1). Purity of fractionation was evaluated using the following antibodies: anti-Actin (cytosol), anti-MEK ½ (nucleoplasm) and anti-H3 (chromatin).[1,2]

#### 10.Protein extraction from tumor tissues

The mouse colorectal cancer samples were obtained in our previous study.[2] The cytoplasmic and nuclear proteins from the mouse colon tissues were extracted by using Cytoplasmic and Nuclear Protein Extraction Kit (BOSTER BIO) according to the manufacturer's protocol. The mixture of cytoplasmic and nuclear protein fraction was used for immunoblotting analysis of TGM2 expression, and the pellet left was used for histone extraction. The extraction of histone was performed according to the previously described acid extraction method.[1,2] Briefly, the chromatin pellet was resuspended in 0.4 N $H_2SO_4$ and rotated at 4 ºC overnight to extract histones. After centrifugation at 16,000 X g for 10 minutes, the supernatant was transferred to a new tube followed by adding 132 μL of 100% TCA to precipitate histones overnight. At last, histones were dried at room temperature after precipitation by centrifugation and being washed with

1 mL cold acetone. The dried histones were then dissolved in 100 μL Milli-Q water. The concentration of each sample was determined using 280 nm wavelength on a Quickdrop (MOLECULAR DEVICES). The histone solution was then diluted to ~0.5 μg/μL for the further labeling assays and analyses.

SUPPLEMENTARY TABLES

| Host | Epitope | Dilution | Vendor |
|---|---|---|---|
| Rabbit | Anti-TGM2 | 1: 500 (tumor)<br>1: 1000 (*in vitro* and *in cellulo*) | CST (3557S) |
| Chicken | Anti-H3 | 1: 1000 | Abcam (ab134198) |
| Mouse | Anti-H3 | 1: 1000 | Abcam (ab10799) |
| Mouse | Anti-Actin | 1: 1000 | CST (3700S) |
| Rabbit | Anti-H3Q5ser | 1: 1000 | Millipore (ABE1791) |
| Rabbit | Anti-H3 | 1: 500 (tumor) | Abcam (ab1791) |
| Rabbit | Anti-H4 | 1: 1000 | CST (13919S) |
| Mouse | Anti-HA | 1: 1000 | CST (2367S) |
| Rabbit | Anti-GAPDH | 1: 1000 | CST (2118S) |

**Supplementary Table 1.** Primary antibodies used in this study.

| Host | Epitope | Label | Dilution | Vendor |
|---|---|---|---|---|
| Donkey | Anti-Chicken | IRDye 800CW | 1: 15000 | Li-Cor |
| Goat | Anti-Mouse | IRDye 680RD | 1: 15000 | Li-Cor |
| Goat | Anti-Mouse | IRDye 800CW | 1: 15000 | Li-Cor |
| Goat | Anti-Rabbit | IRDye 800CW | 1: 15000 | Li-Cor |
| Goat | Anti-Rabbit | IRDye 680RD | 1: 15000 | Li-Cor |

**Supplementary Table 2.** Secondary antibodies used in this study.

#### 11. NMR Spectrums

$^1$H NMR spectrum of **S4**

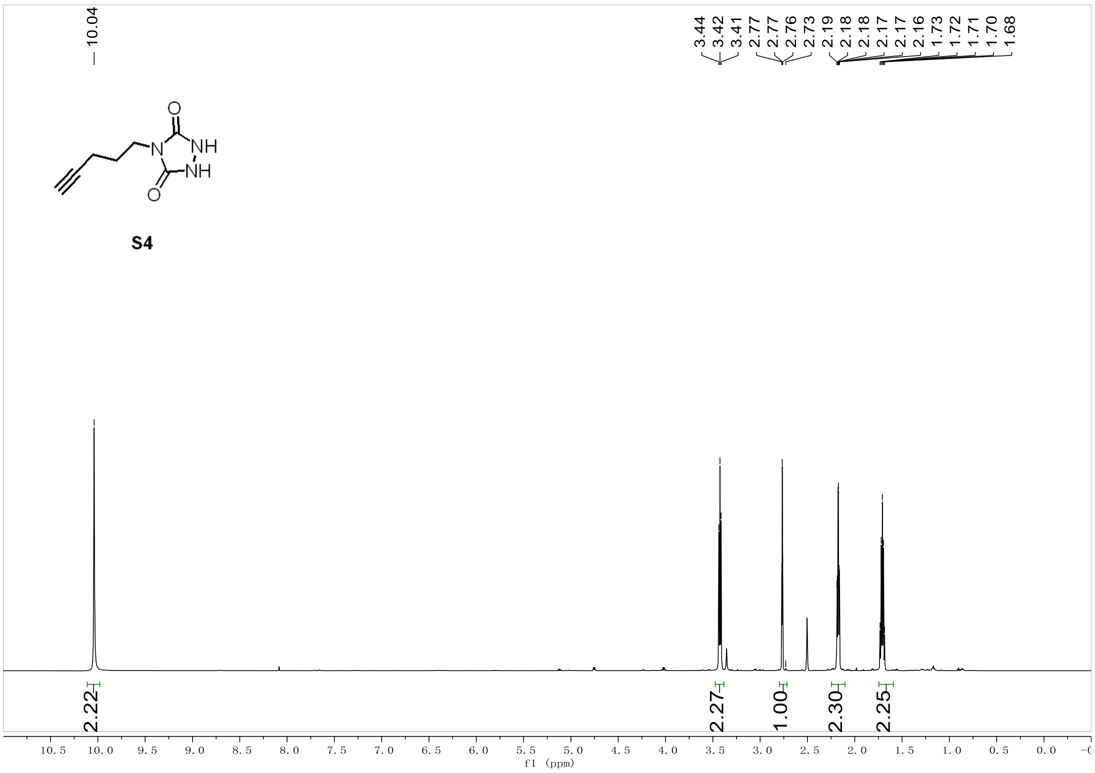

<sup></sup>

$^{13}$C NMR spectrum of **S4**

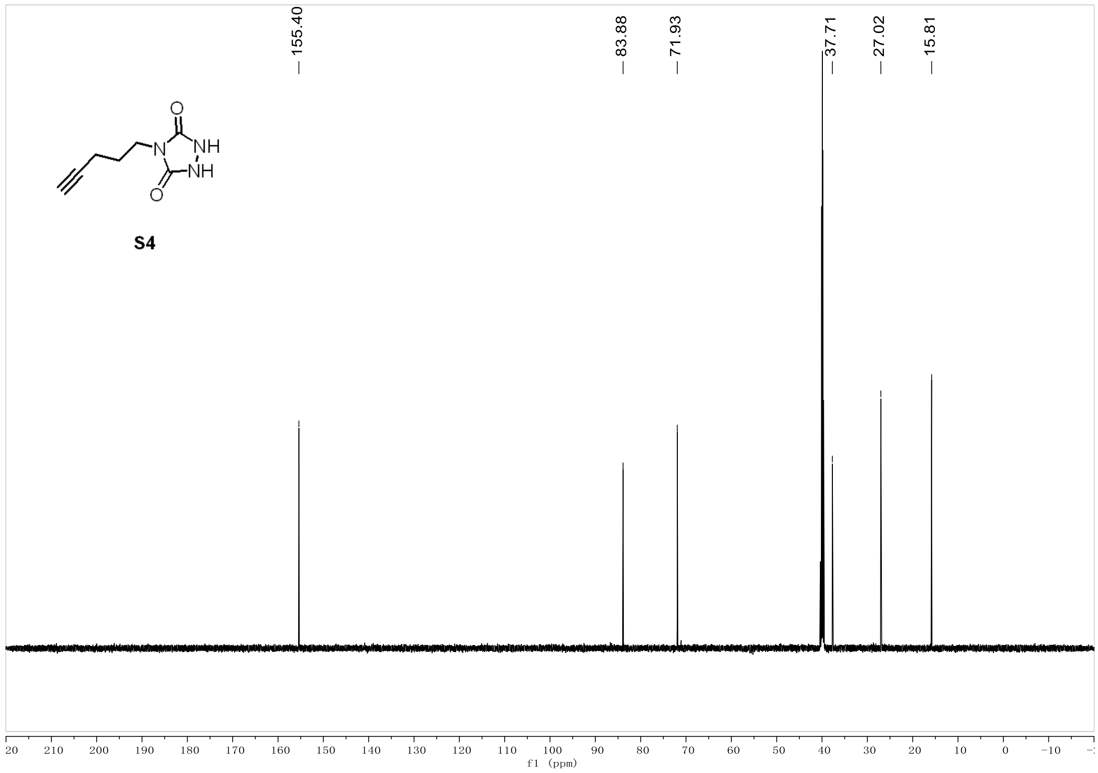

<sup></sup>

$^1$H NMR spectrum of AlkTAD

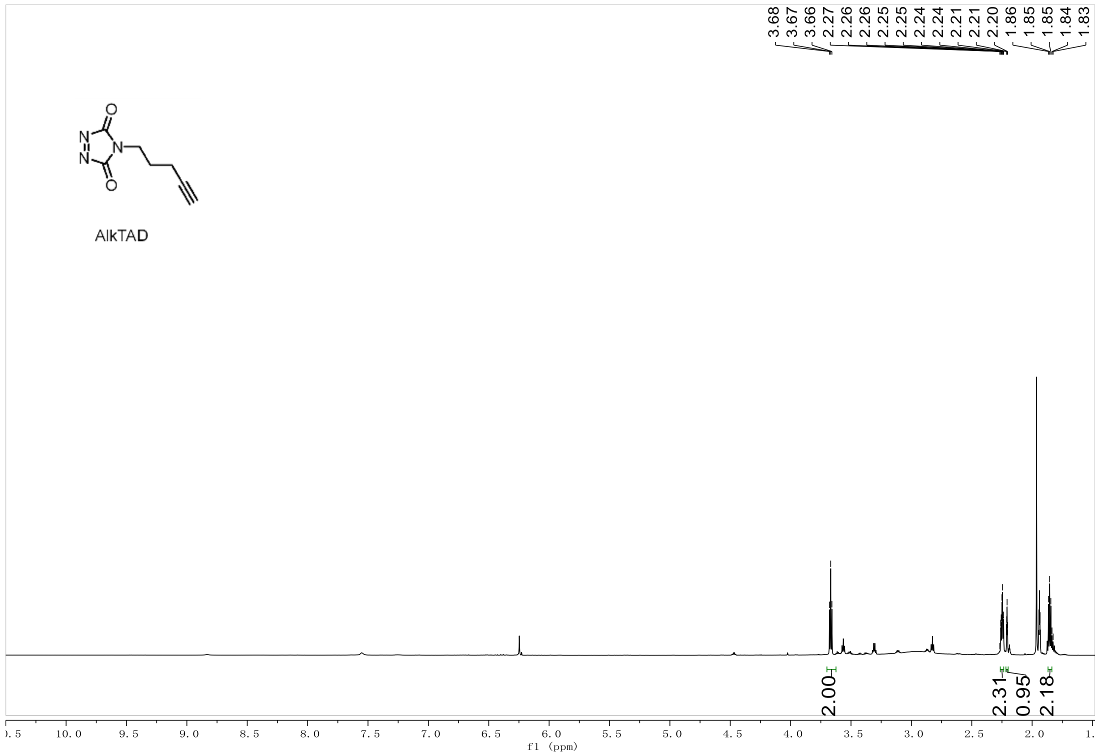

$^{13}$C NMR spectrum of AlkTAD

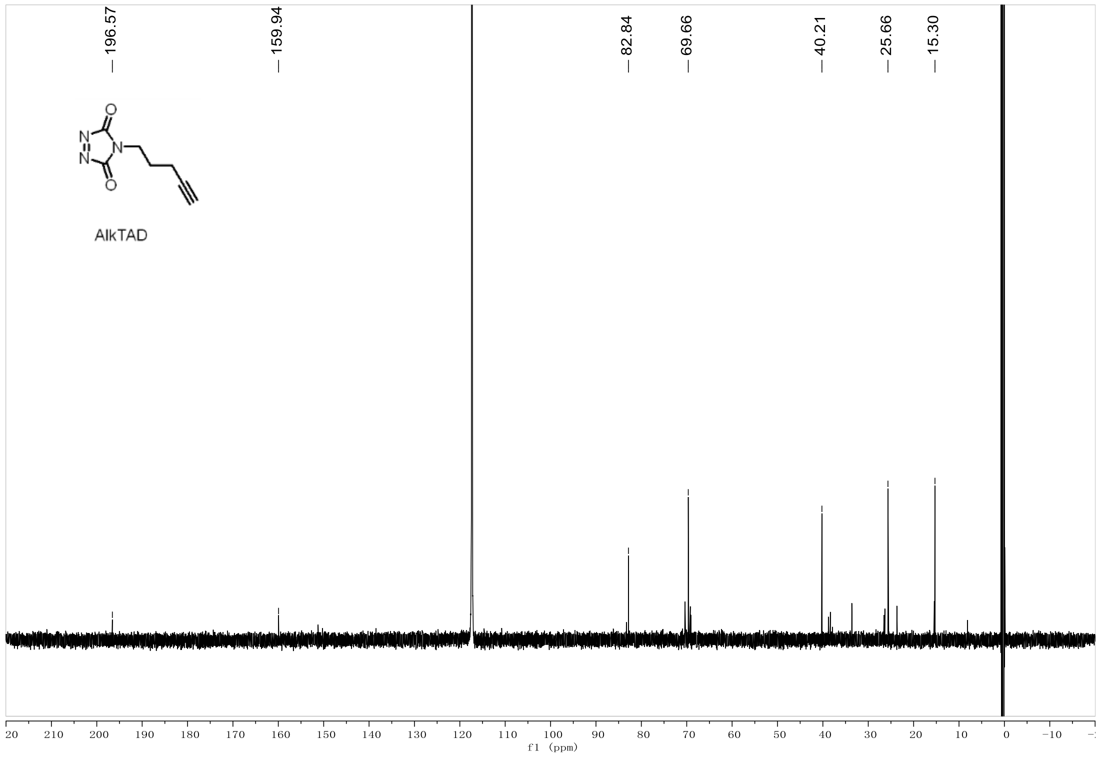

$^1$H NMR spectrum of **S6**

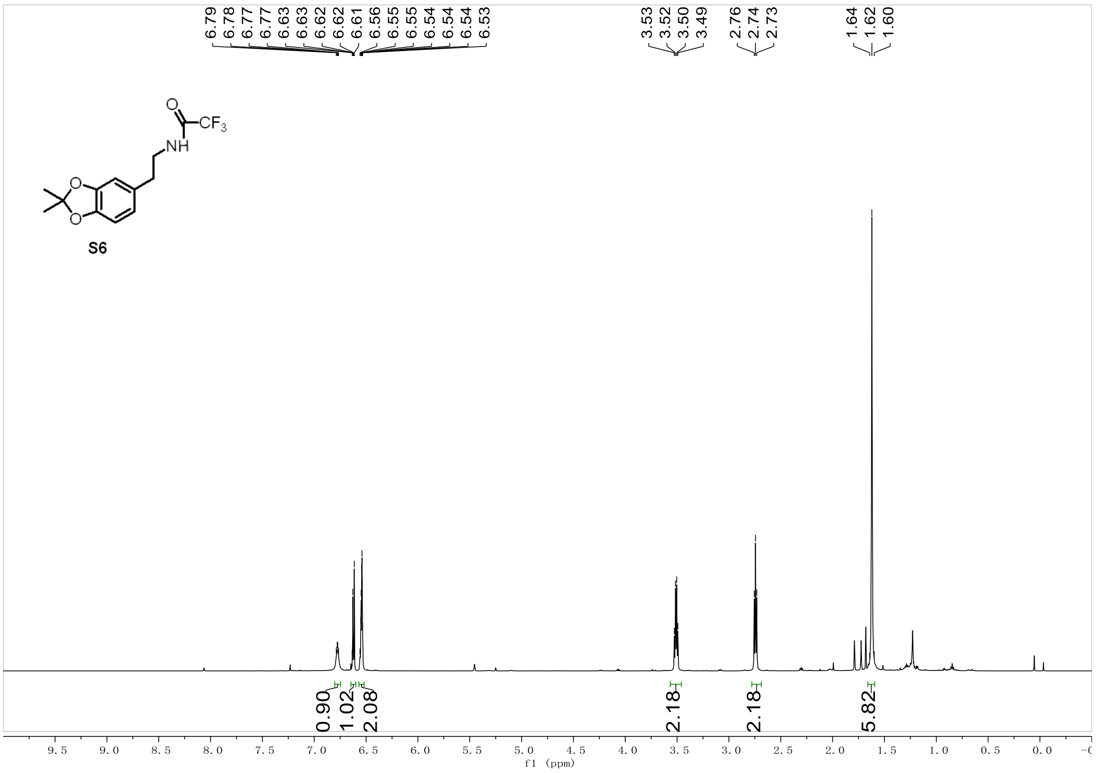

$^1$H NMR spectrum of AcNH-AWA-CONH2

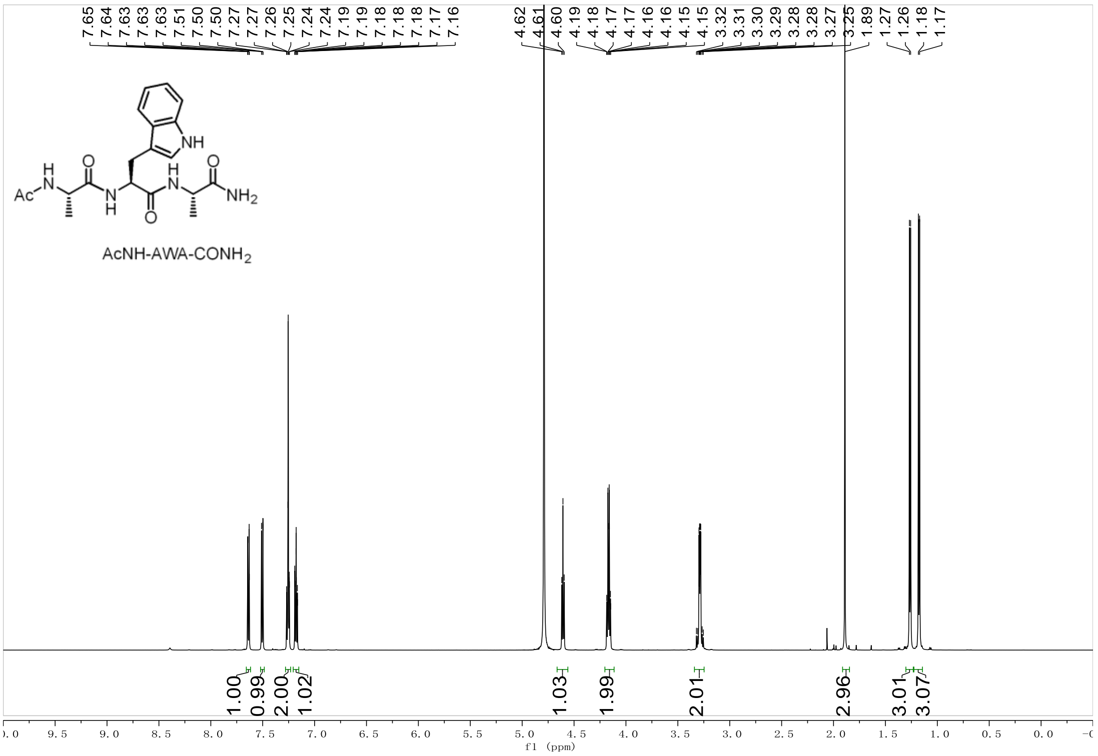

$^{13}C$ NMR spectrum of AcNH-AWA-$CONH_2$

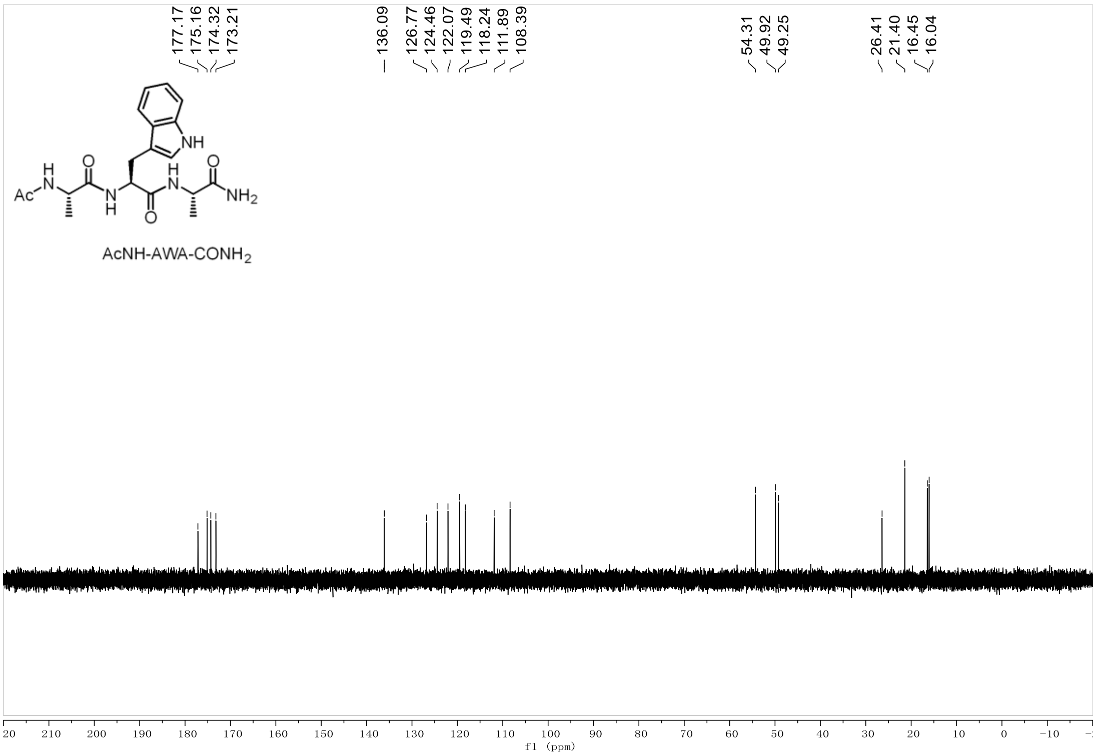

$^{1}H$ NMR spectrum of AcNH-AQserA-$CONH_2$

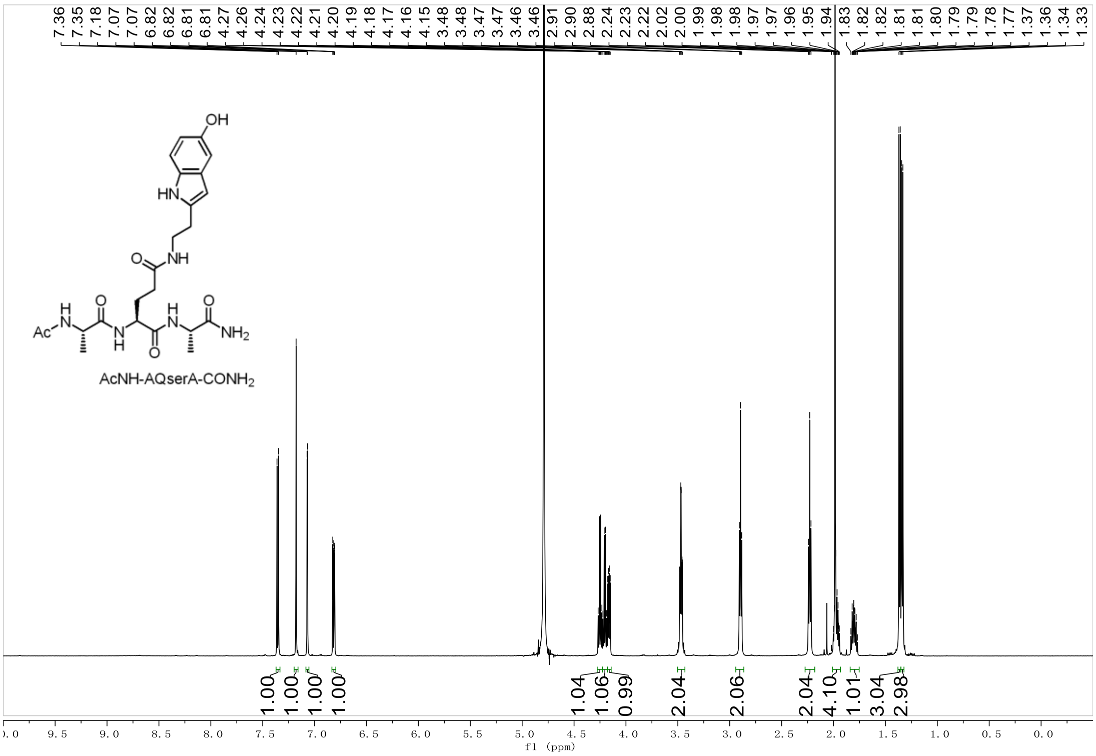

#### $^{13}C$ NMR spectrum of AcNH-AQserA-$CONH_2$

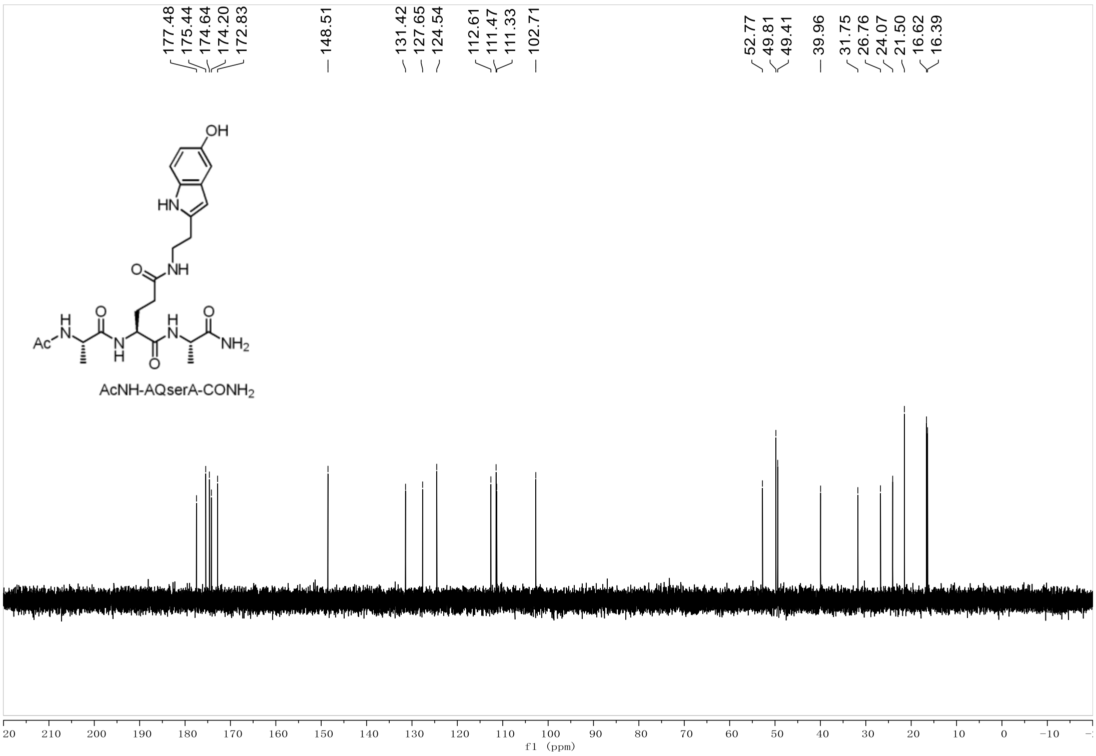

#### $^{1}H$ NMR spectrum of **1**

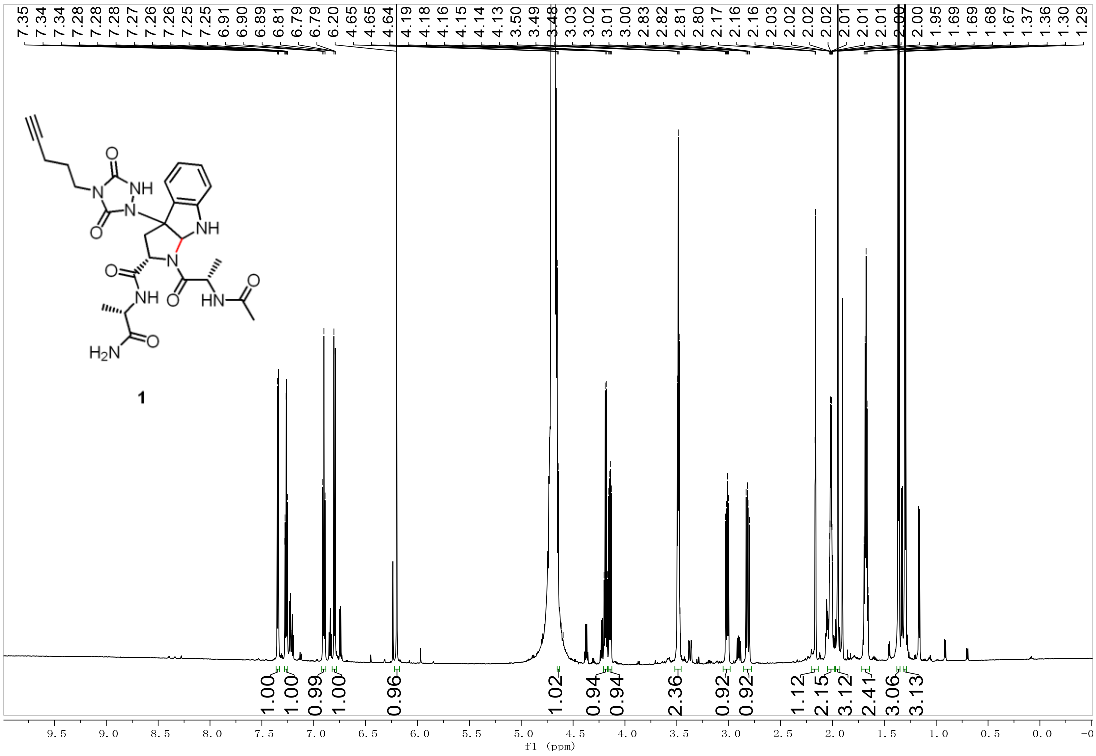

$^{13}C$ NMR spectrum of **1**

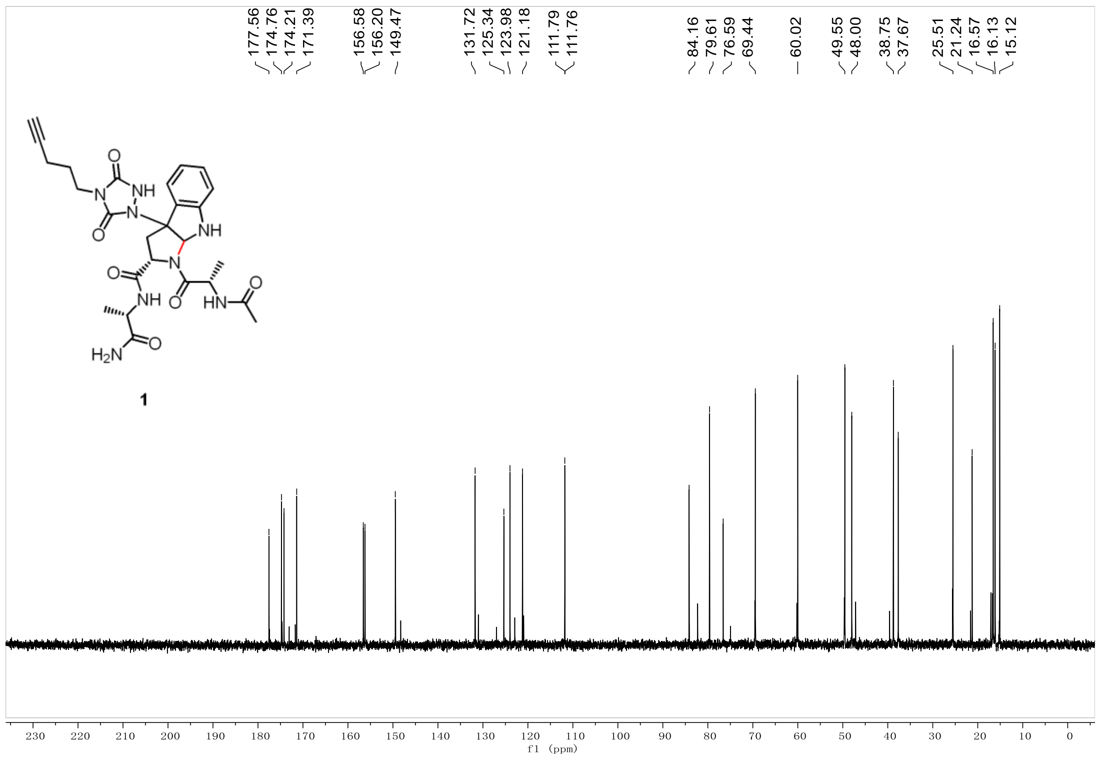

2D NMR (HMBC) spectrum of **1**

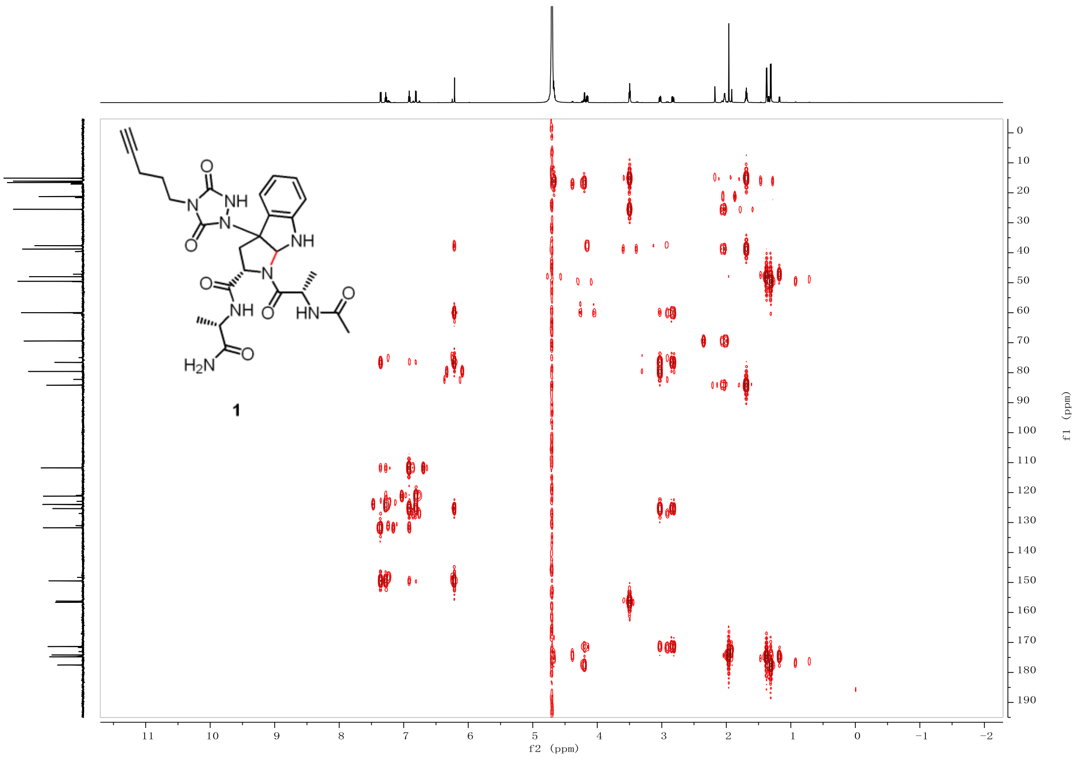

2D NMR (HSQC) spectrum of **1**

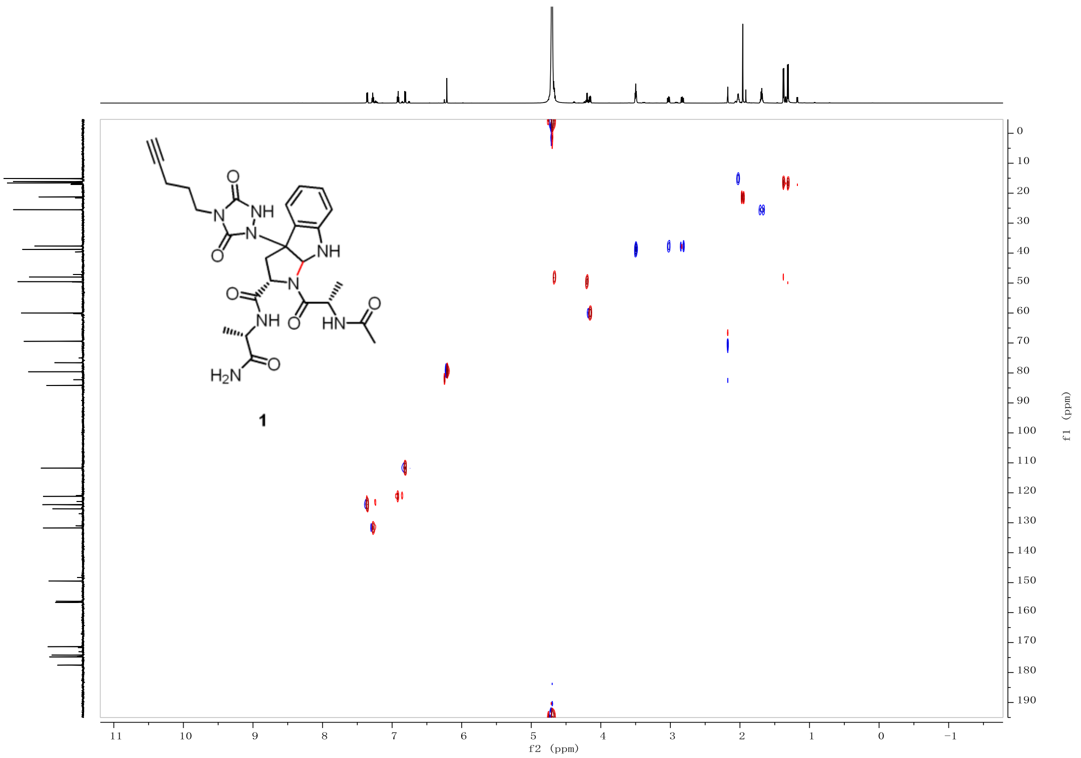

2D NMR (H-H COSY) spectrum of **1**

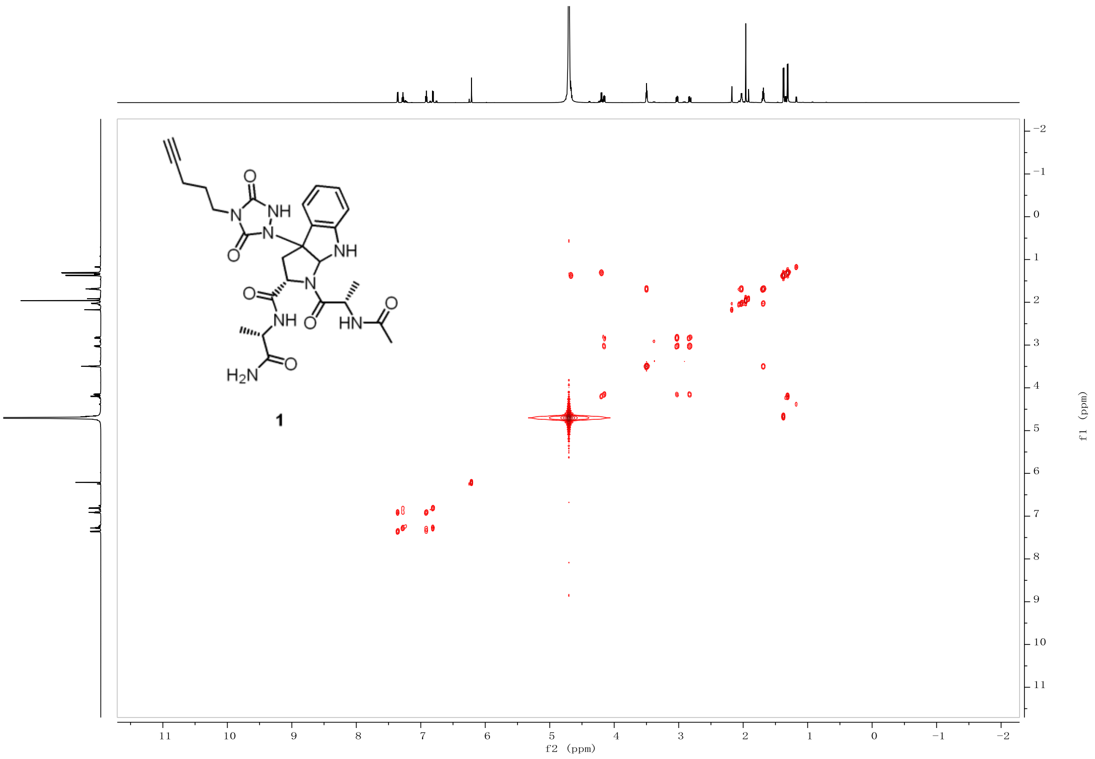

$^{1}$H NMR spectrum of AcNH-AWA-CONH$_2$ + AlkTAD **Component 1**.

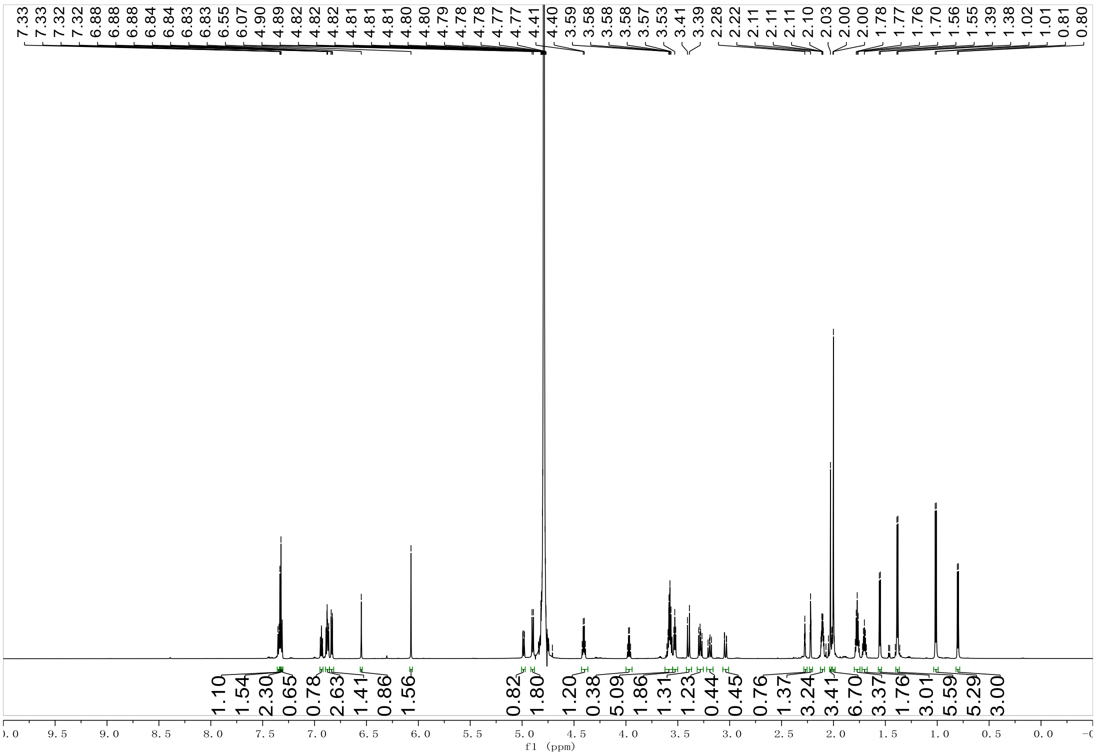

[13]C NMR spectrum of AcNH-AWA-CONH$_2$ + AlkTAD **Component 1**.

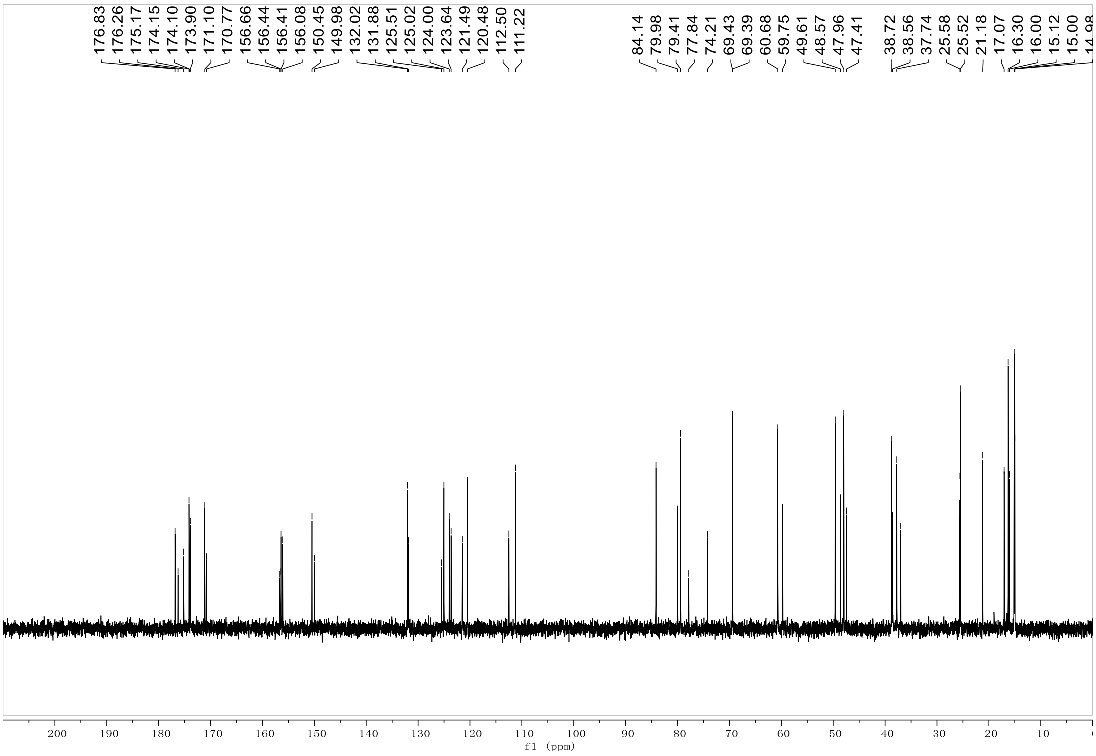

[1]H NMR spectrum of **2**

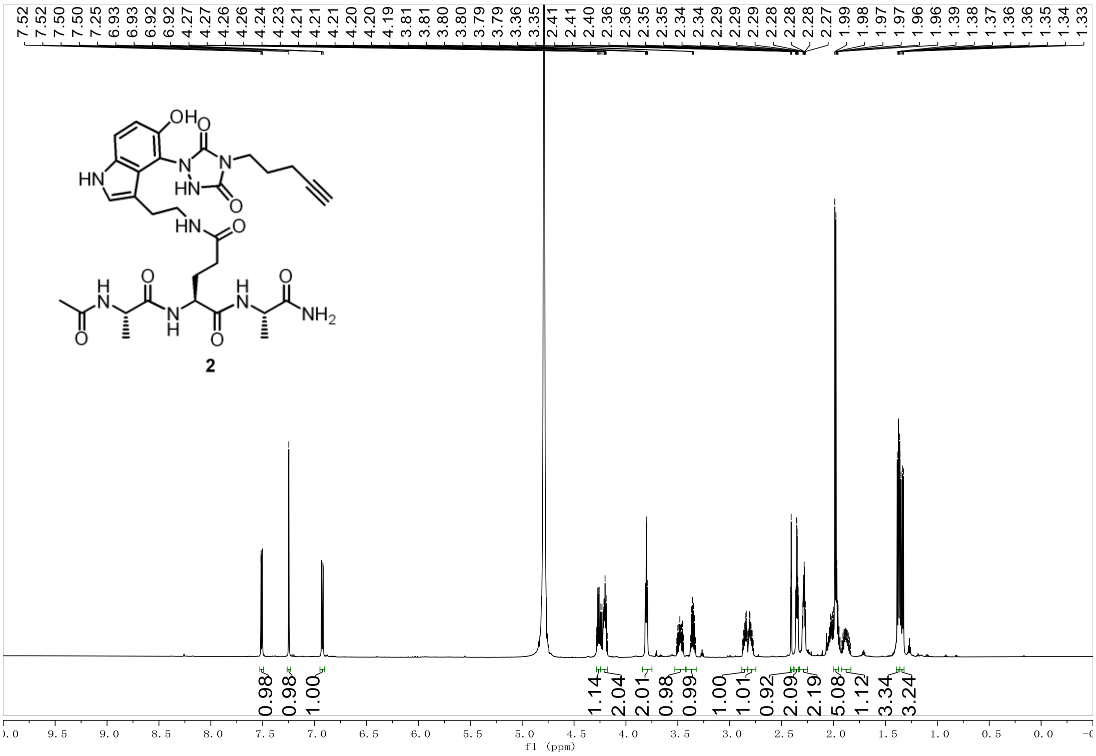

$^{13}C$ NMR spectrum of **2**

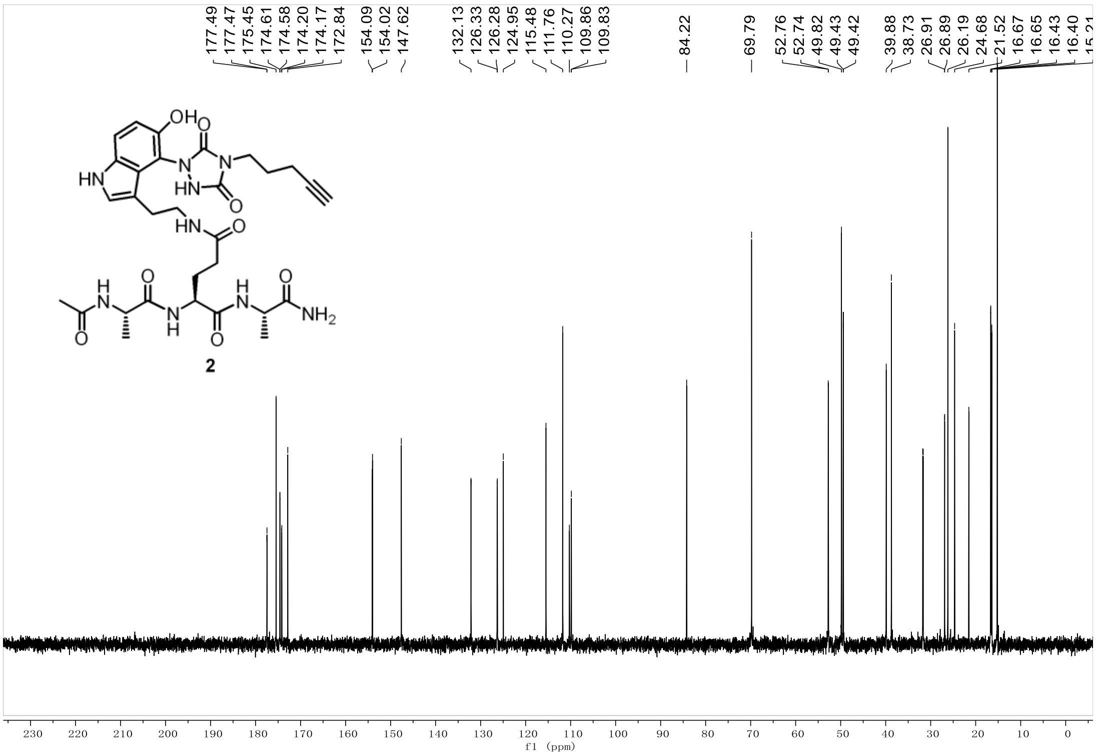

2D NMR (HMBC) spectrum of **2**

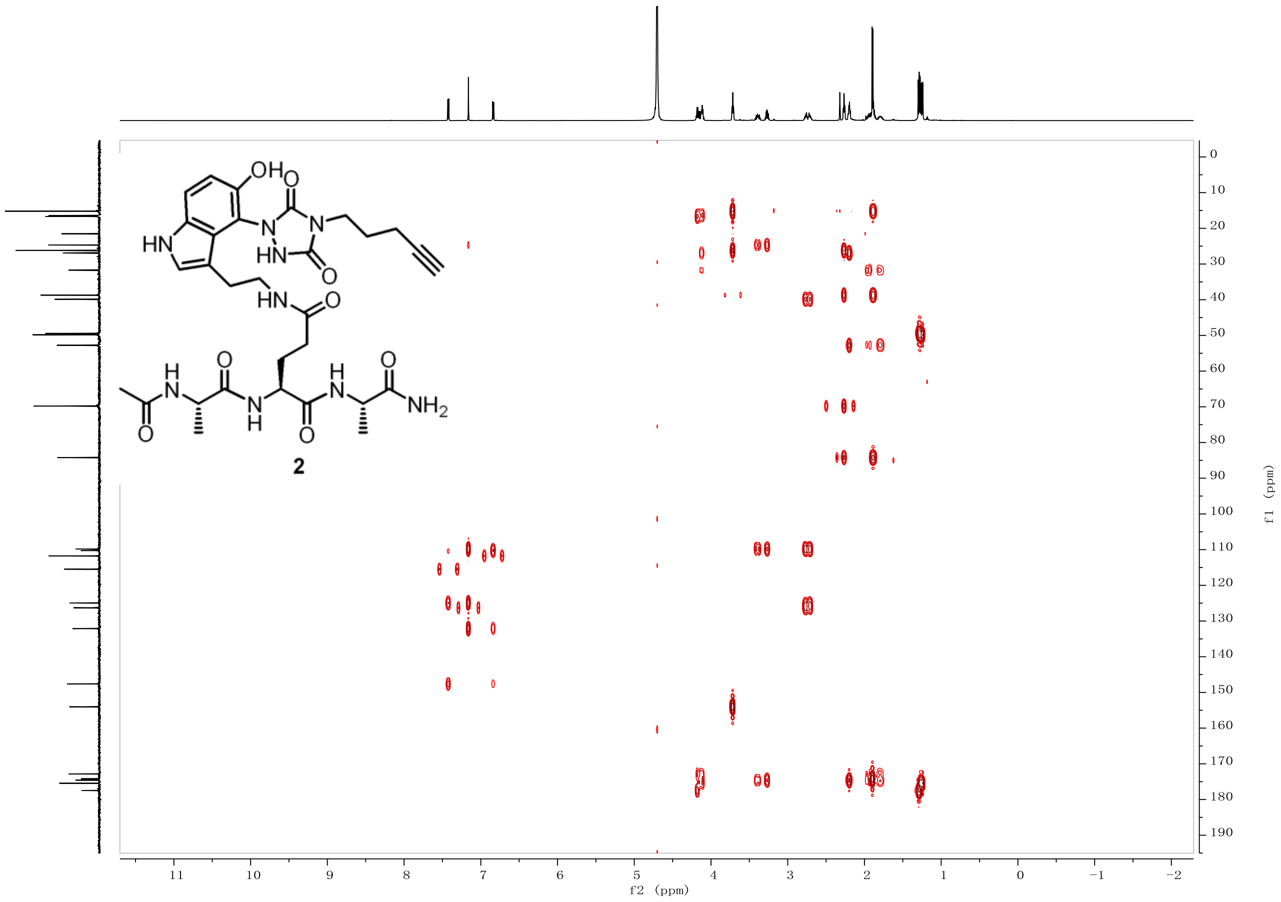

2D NMR (HSQC) spectrum of **2**

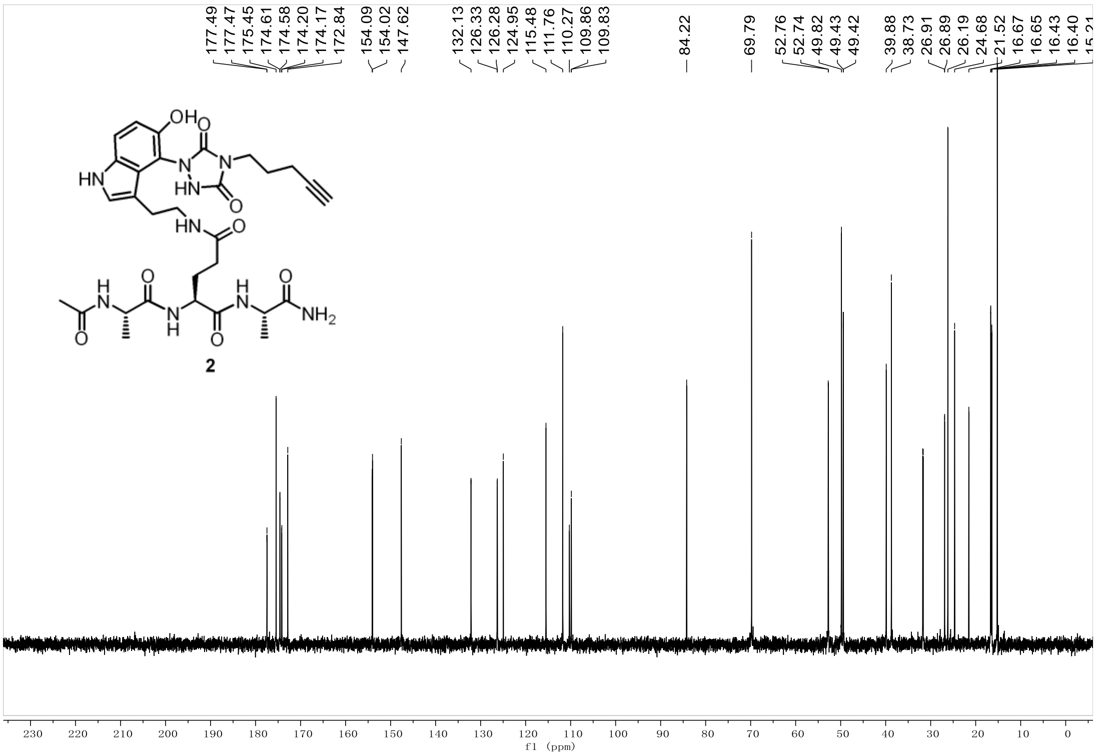

2D NMR (H-H COSY) spectrum of **2**

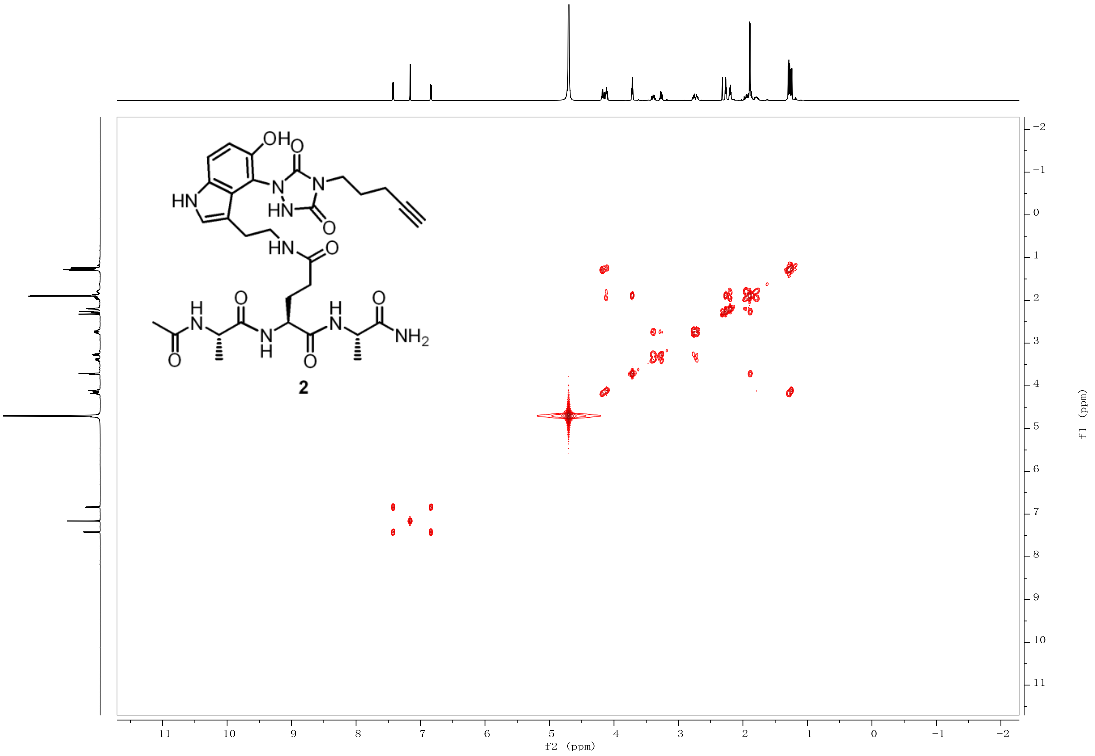

#### 12.MS spectrums

MS spectrums of ARTKQTARKSTG

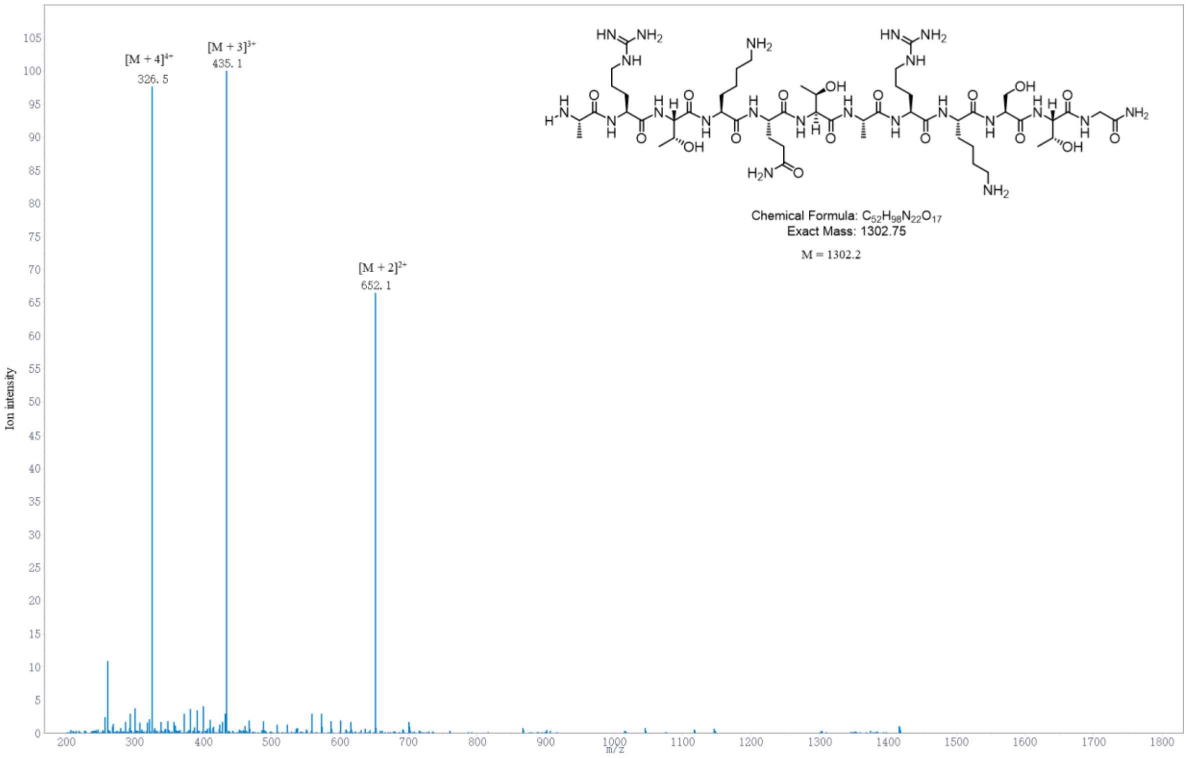

MS spectrums of ARTKQserTARKSTG

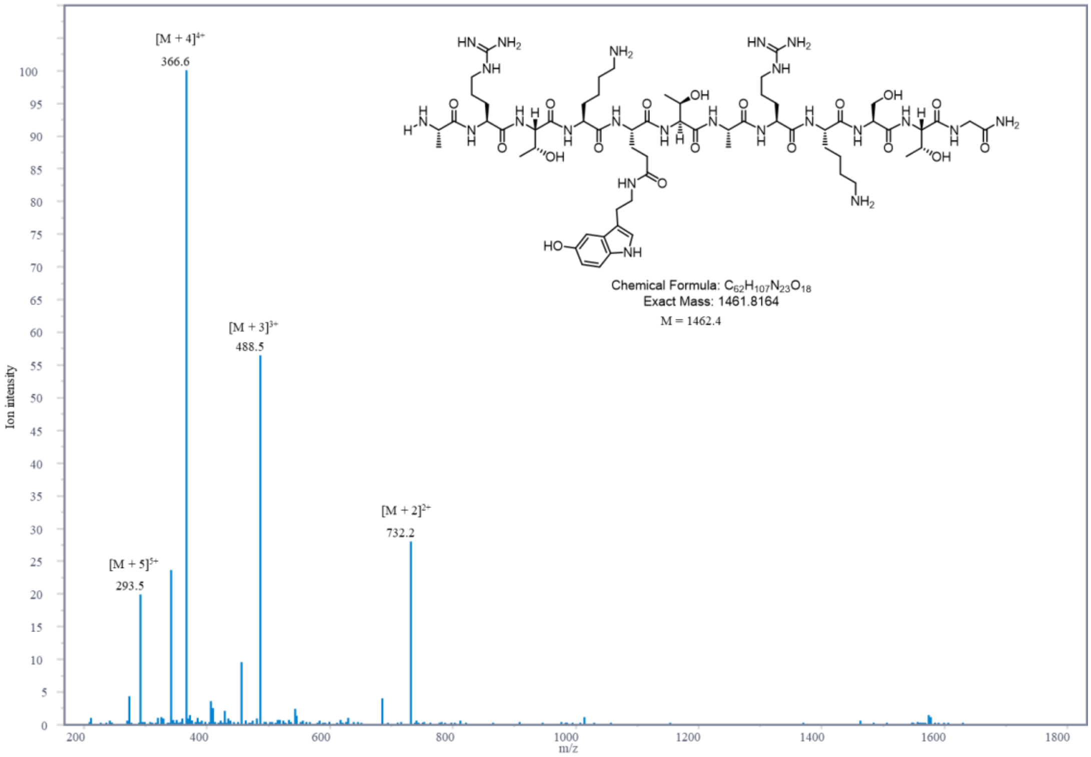

MS spectrums of ARTKQserTARKSTG + AlkTAD

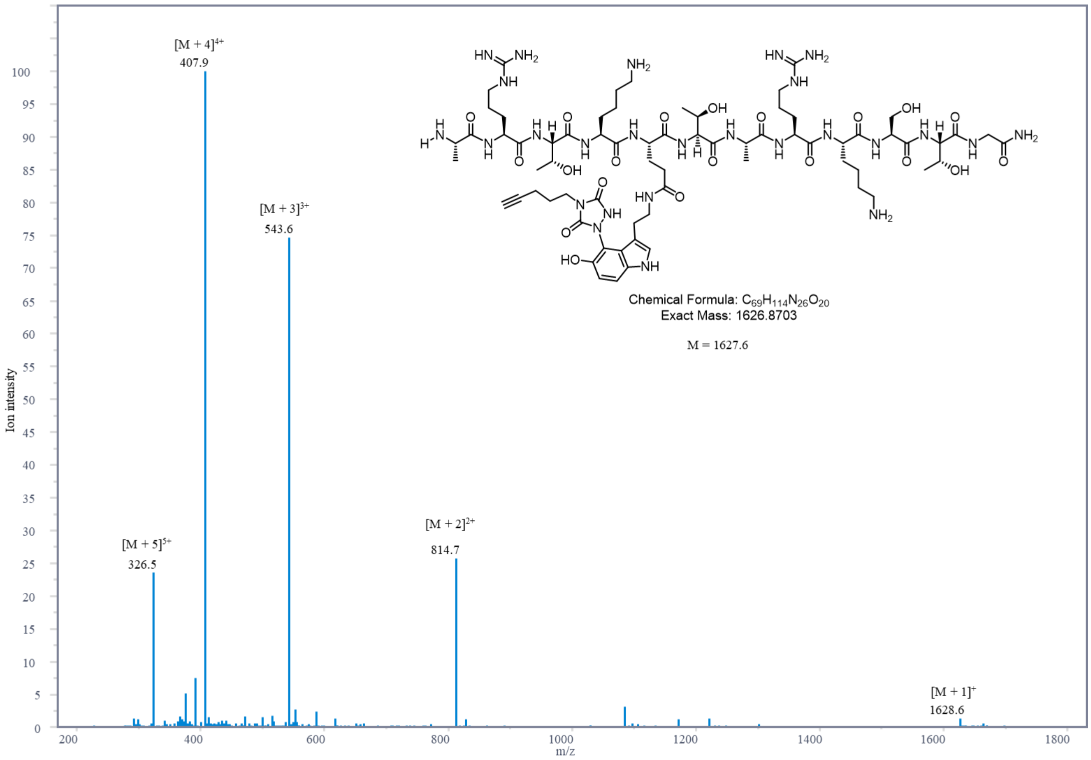

MS spectrums of ARTKWTARKSTG

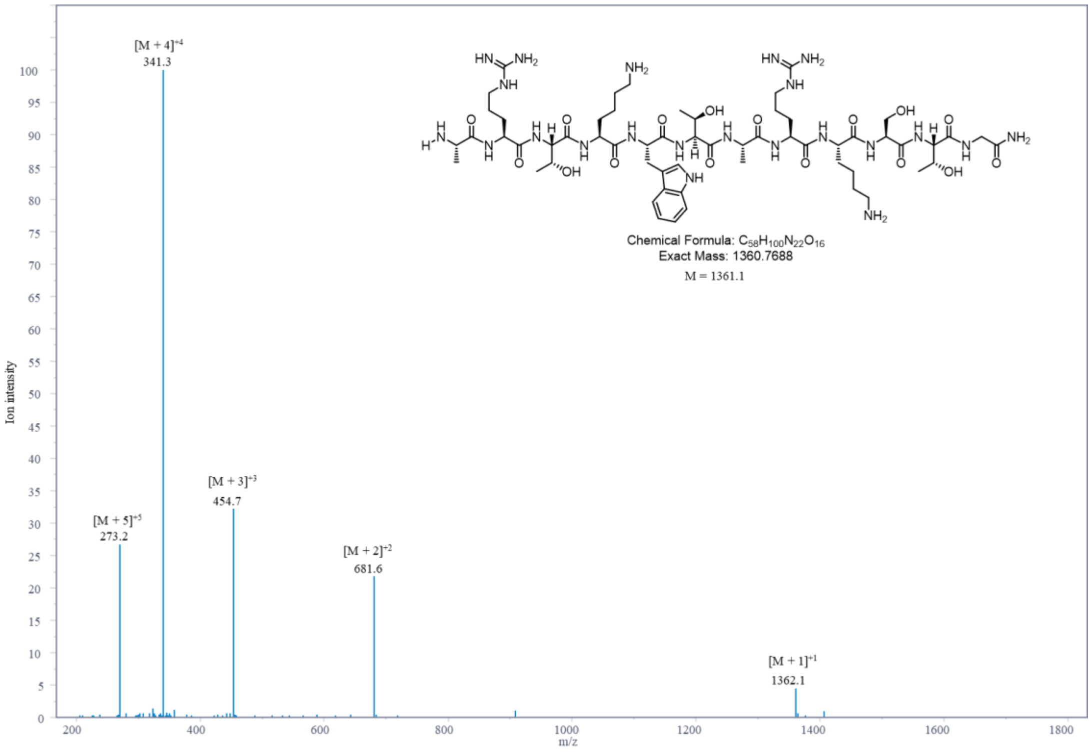

MS spectrums of ARTKWTARKSTG + AlkTAD

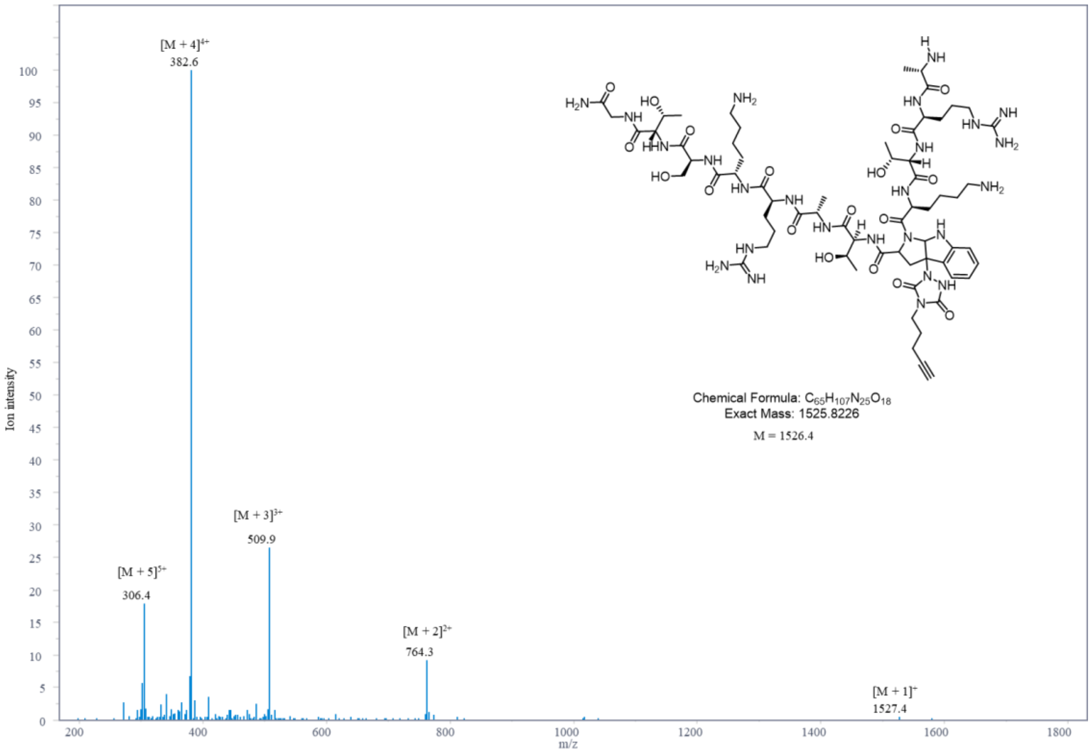

MS spectrums of ARTKYTARKSTG

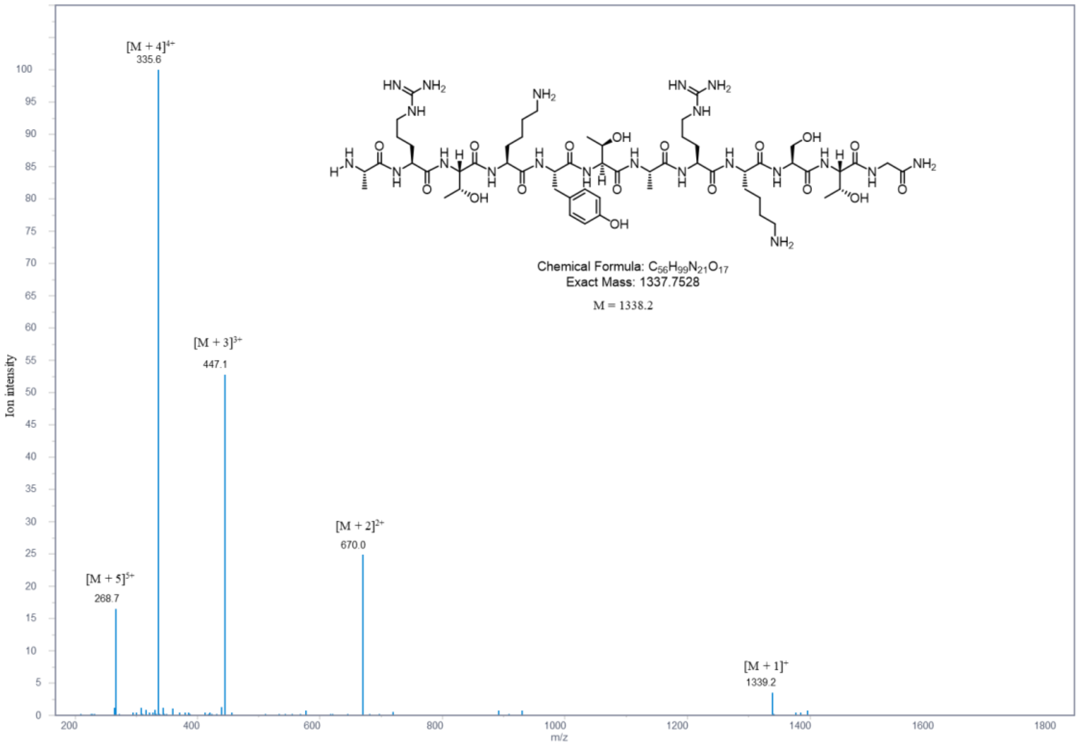

MS spectrums of ARTKYTARKSTG + AlkTAD

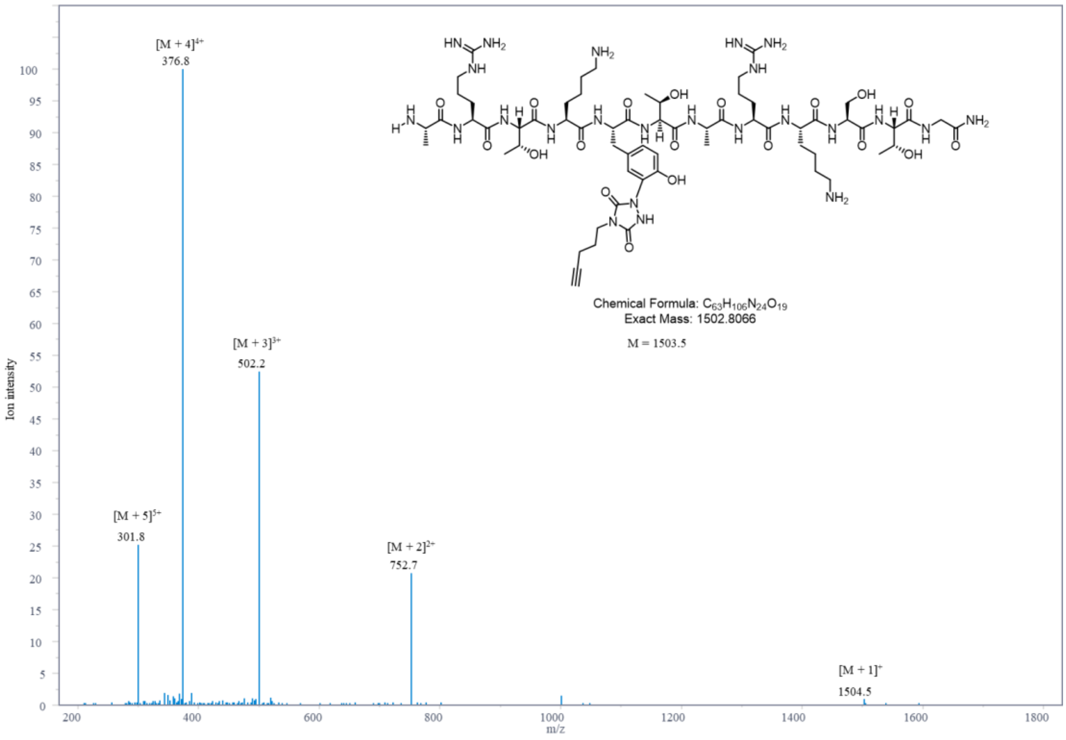

MS spectrums of ARTKHTARKSTG

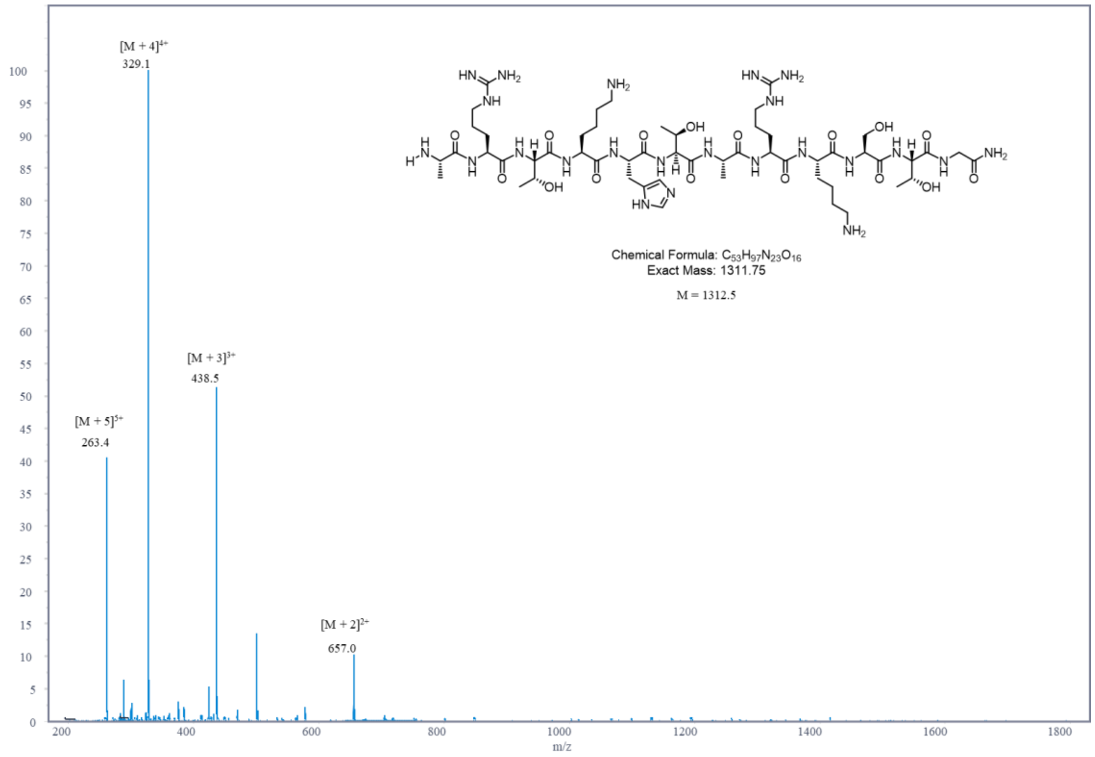

MS spectrums of ARTKQtyrTARKSTG

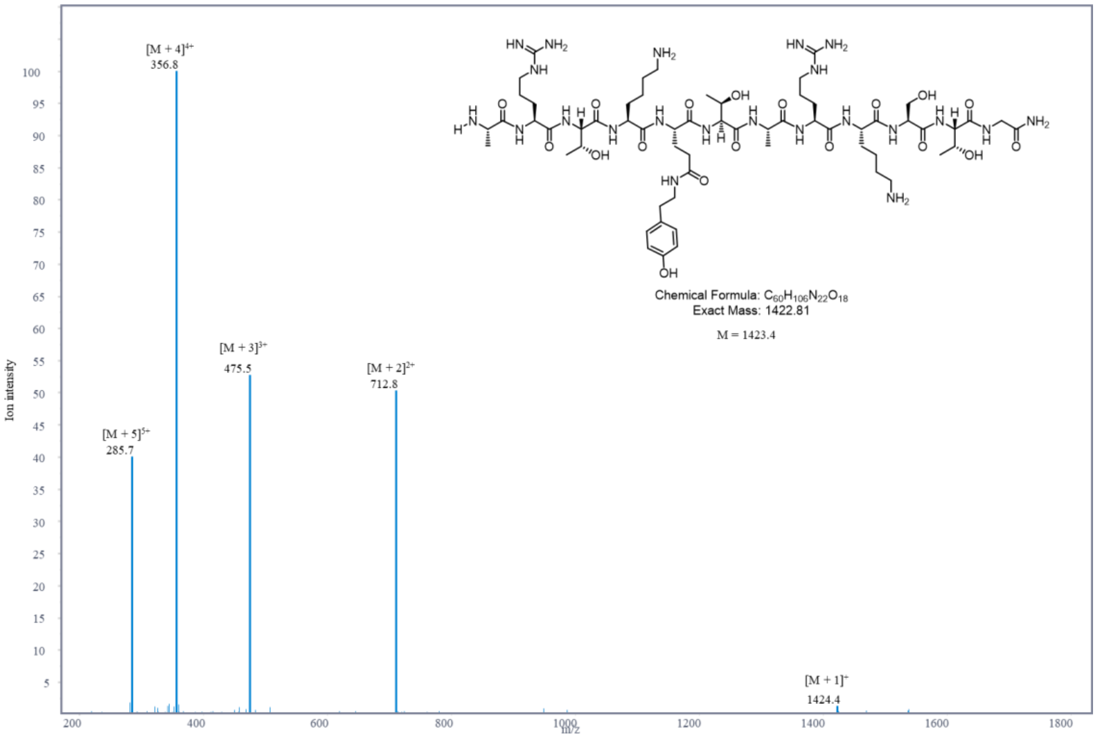

MS spectrums of ARTKQtyrTARKSTG + AlkTAD

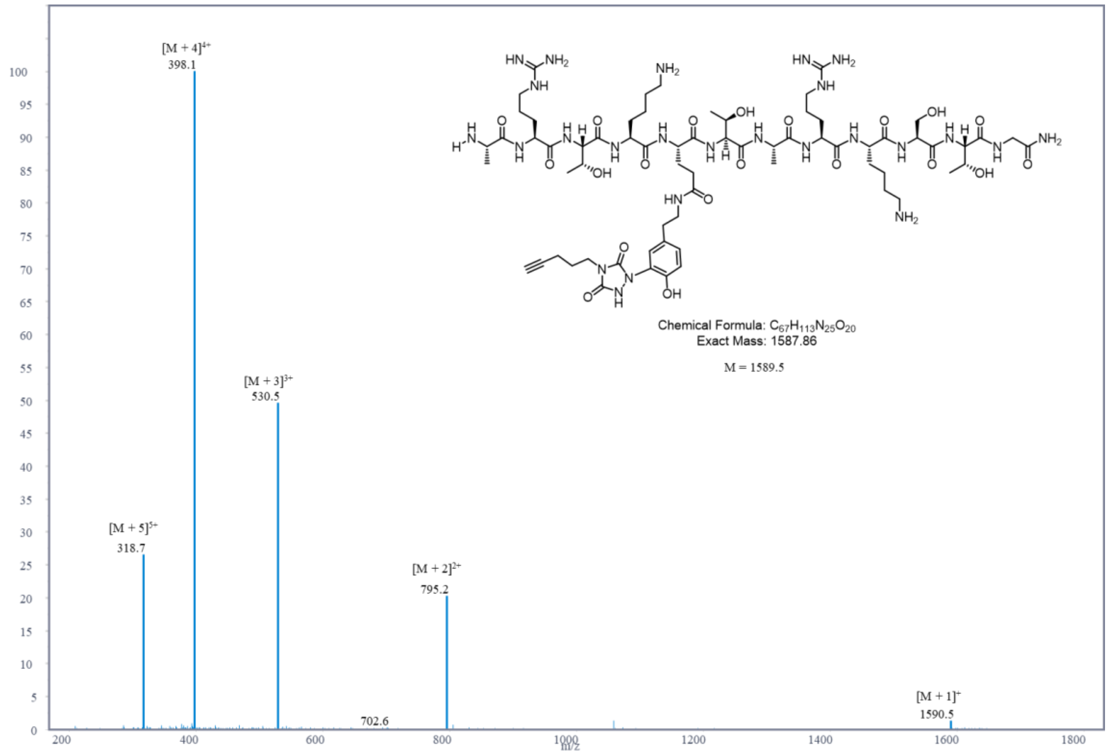

MS spectrums of ARTKQhisTARKSTG

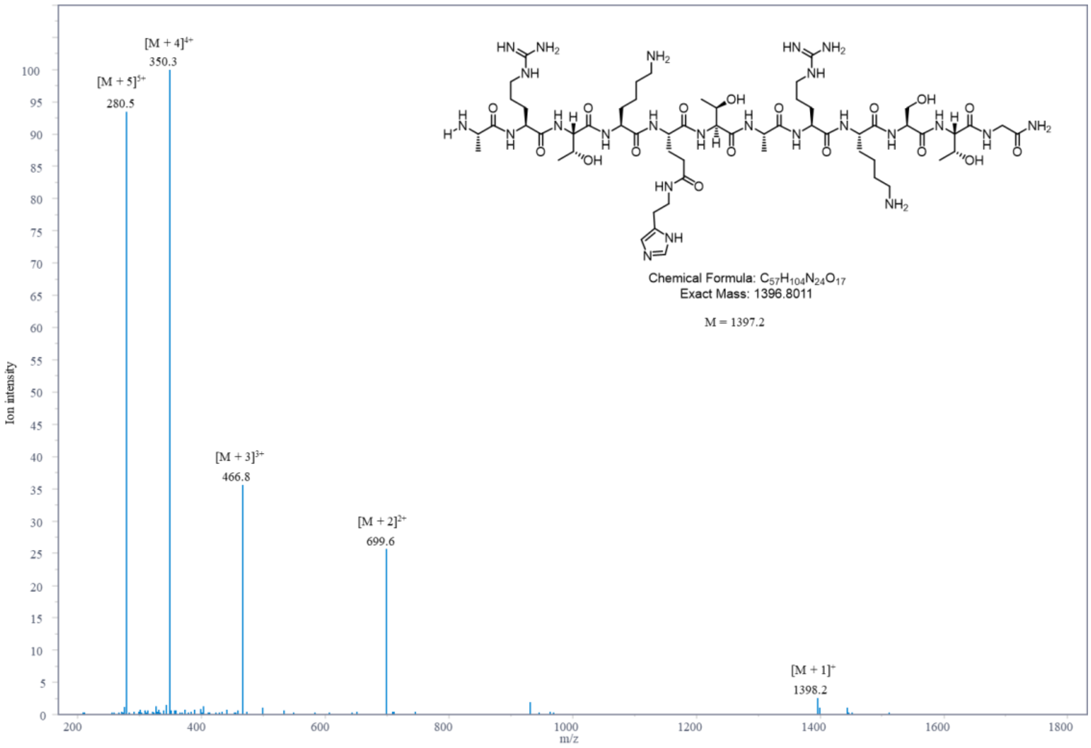

MS spectrums of ARTKQdopTARKSTG

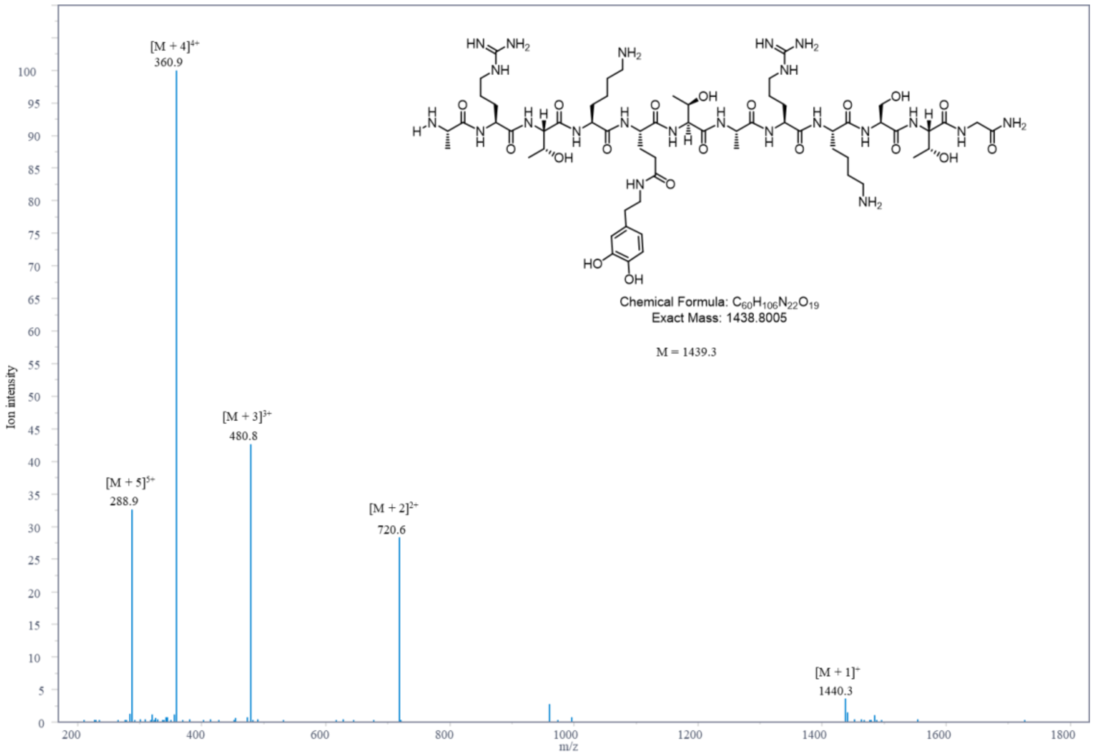

MS spectrums of ARTKQdopTARKSTG + AlkTAD

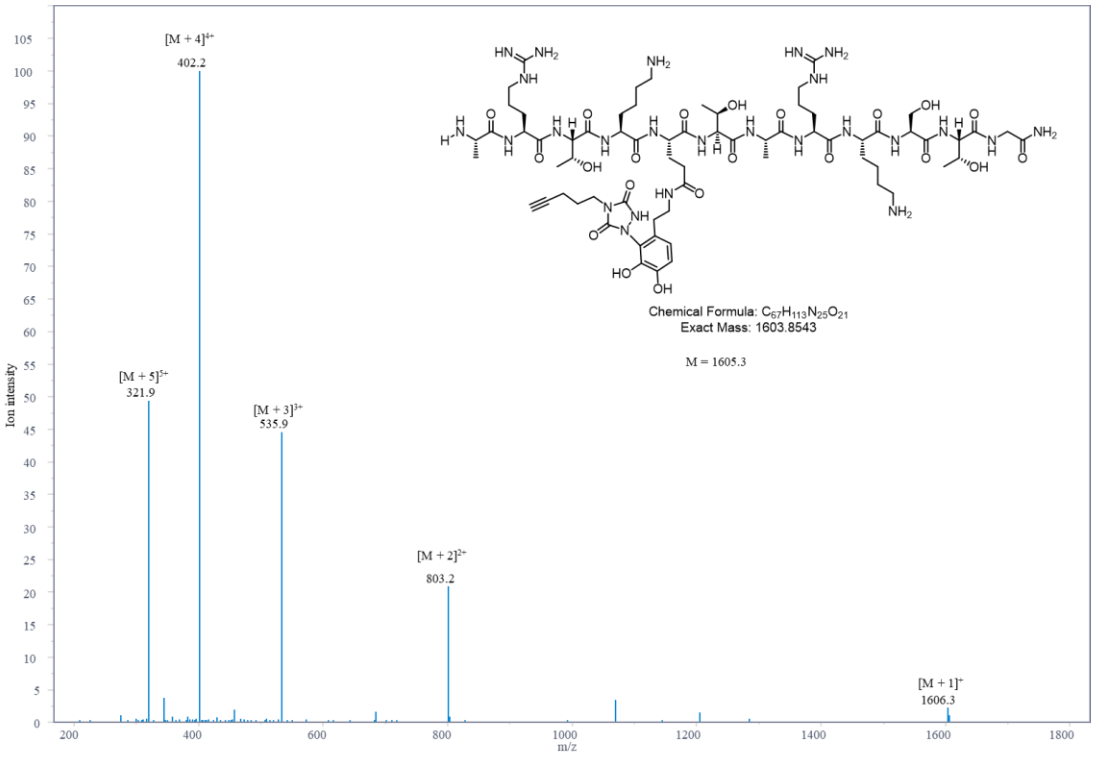

MS spectrums of ARTKQdopTARKSTG + 2AlkTAD

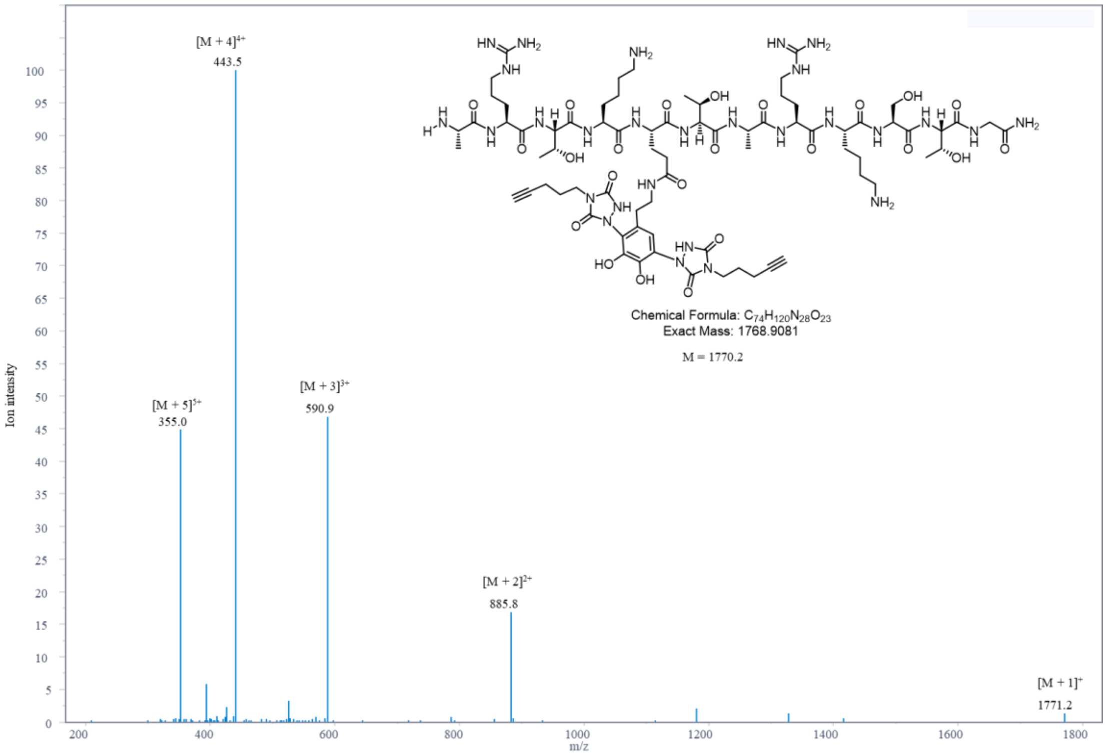

MS spectrums of ARTKQ(3-MethoxyTyr)TARKSTG

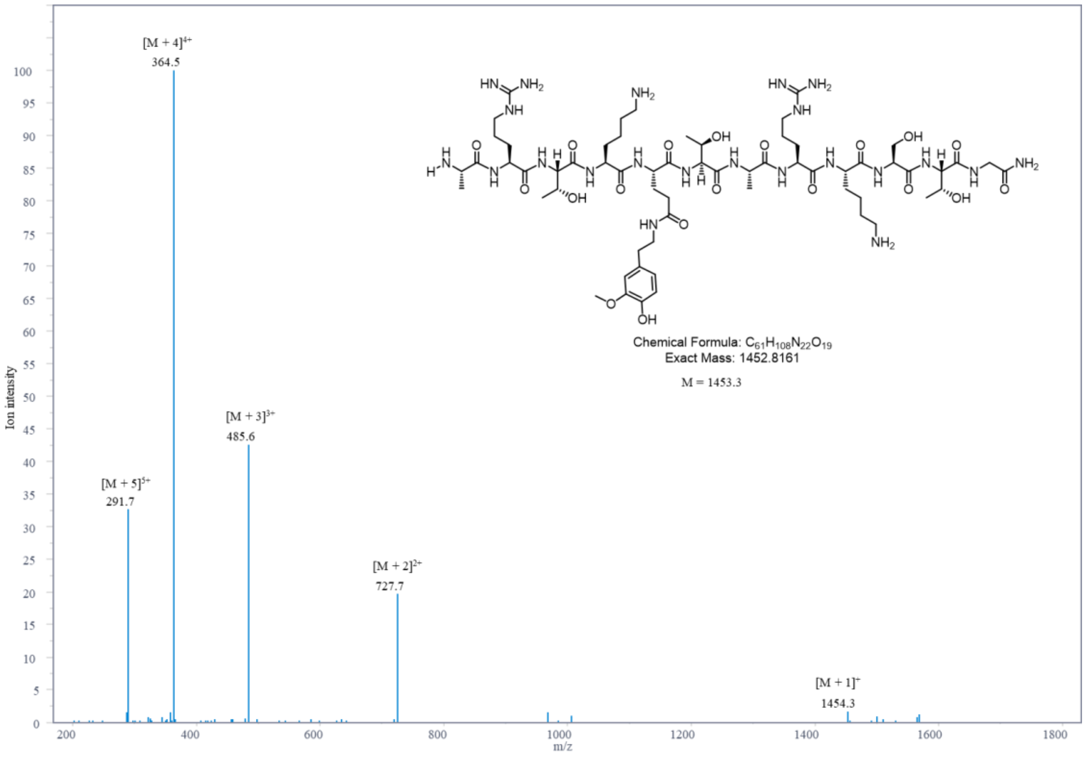

MS spectrums of ARTKQ(3-MethoxyTyr)TARKSTG + AlkTAD

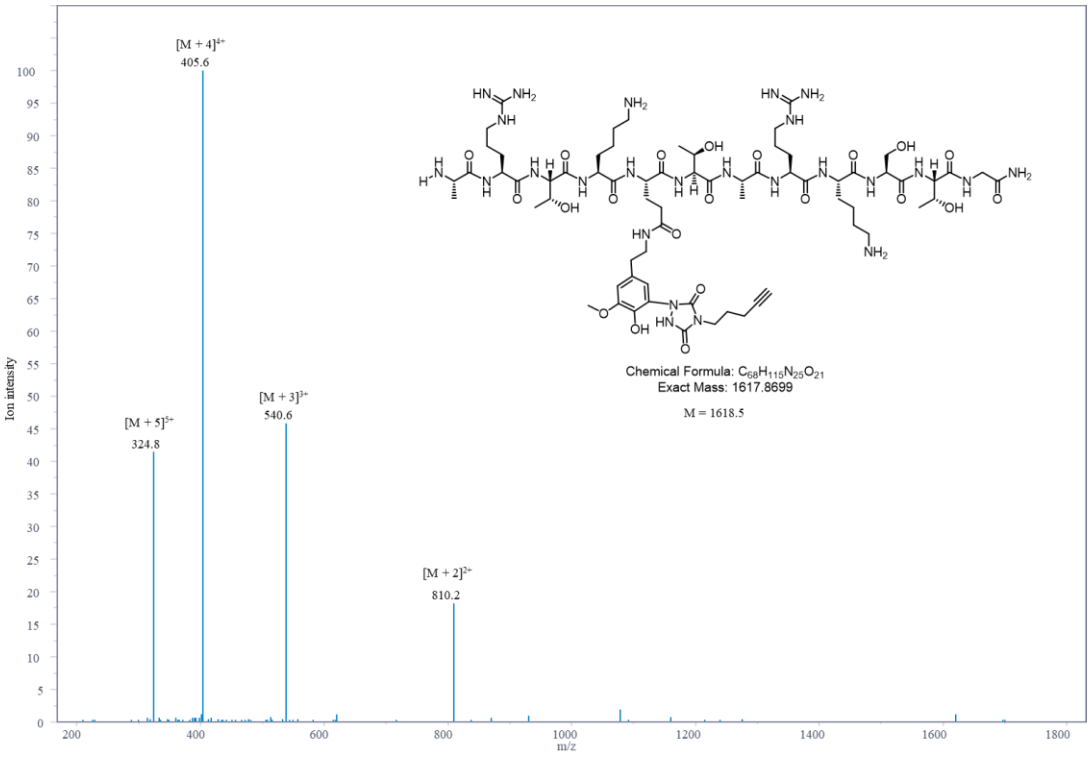

MS spectrums of ARTKQmetyrTARKSTG

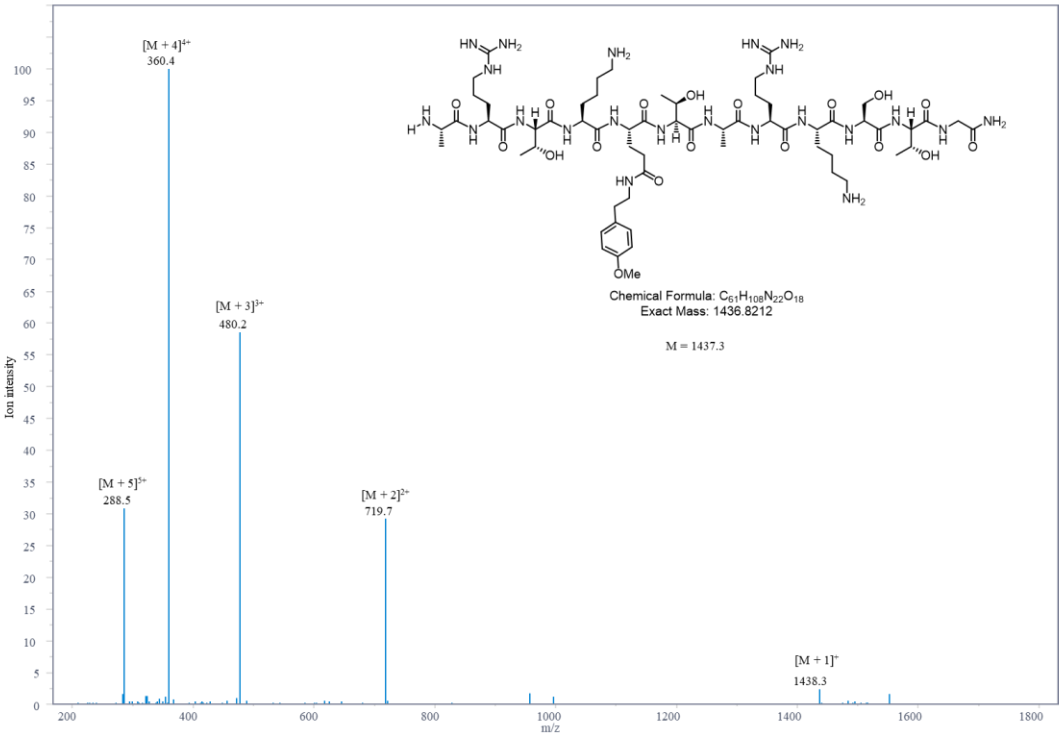

MS spectrums of ARTKQ(5-MT)TARKSTG

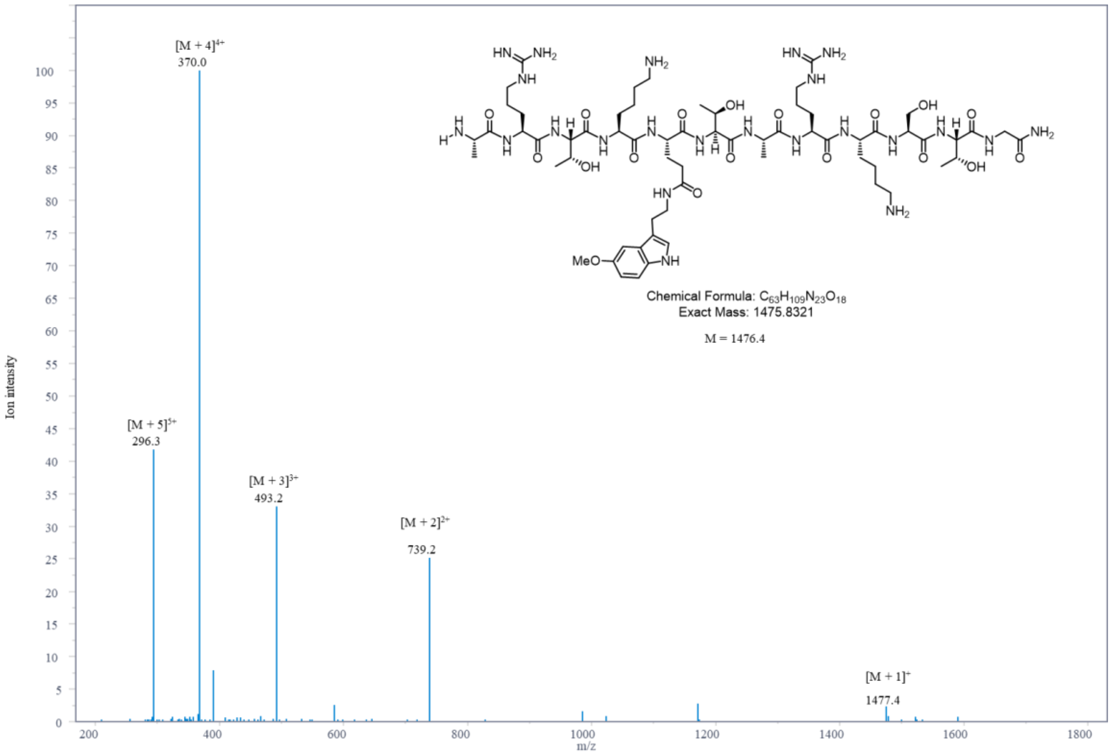

MS spectrums of ARTKQ(5-MT)TARKSTG + AlkTAD

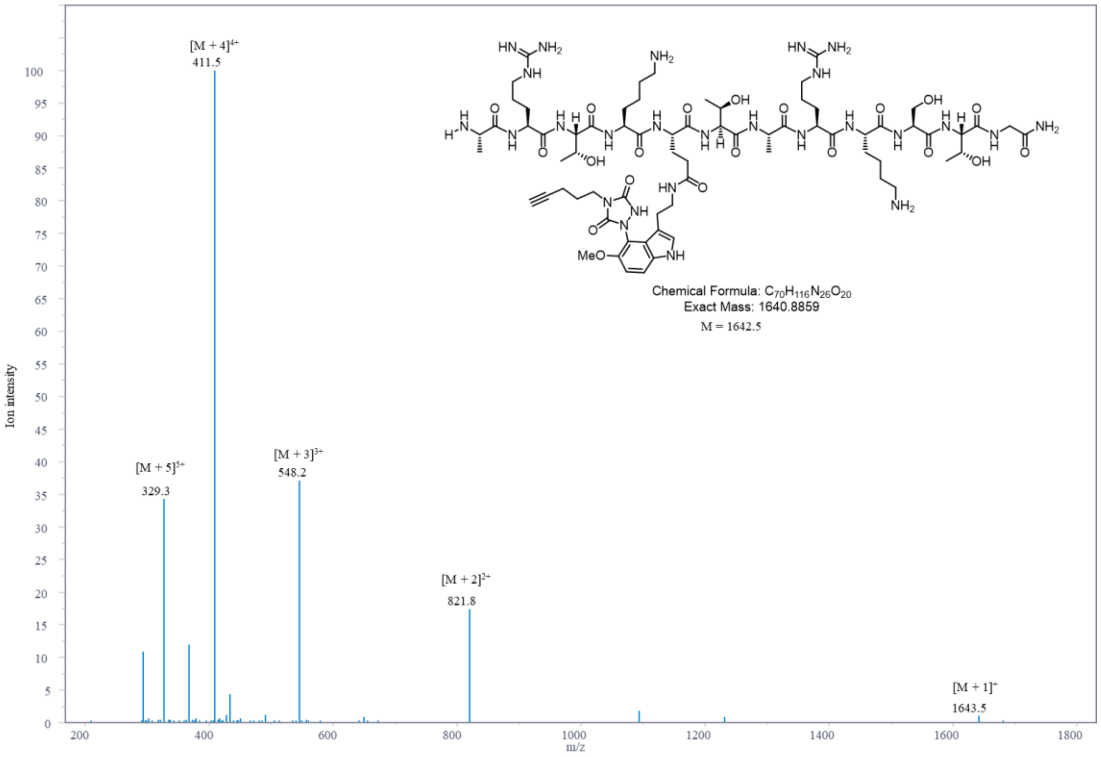

MS spectrums of ARTKQ(5-F-Try)TARKSTG

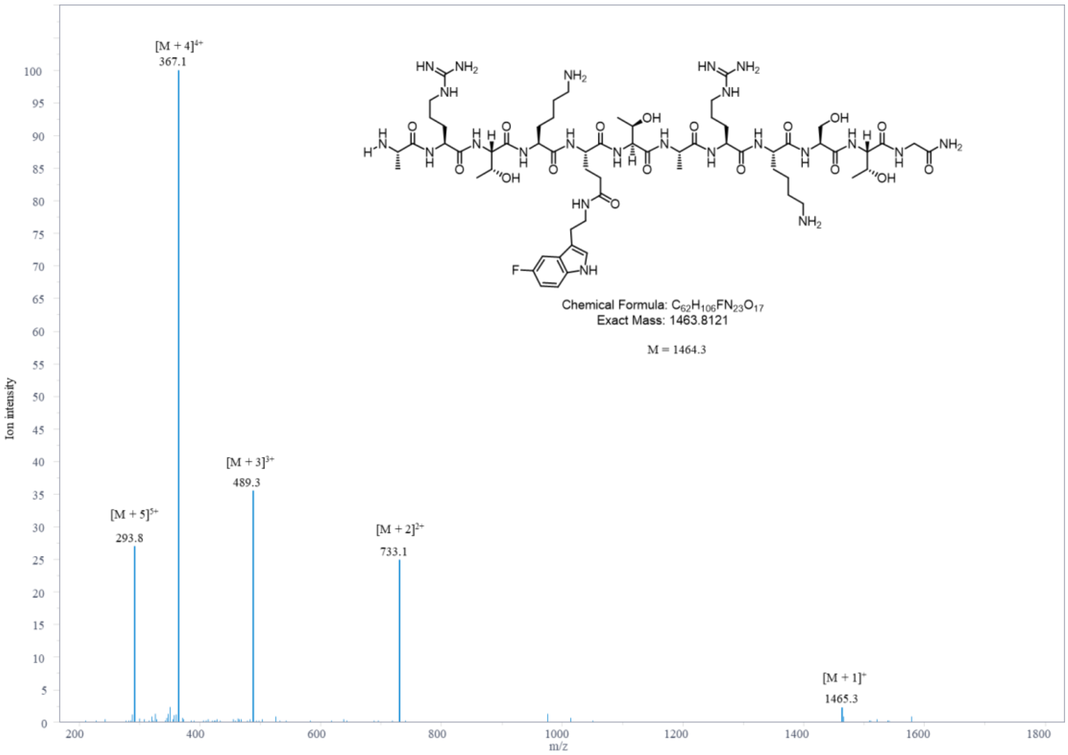

MS spectrums of ARTKQ(5-F-Try)TARKSTG + AlkTAD

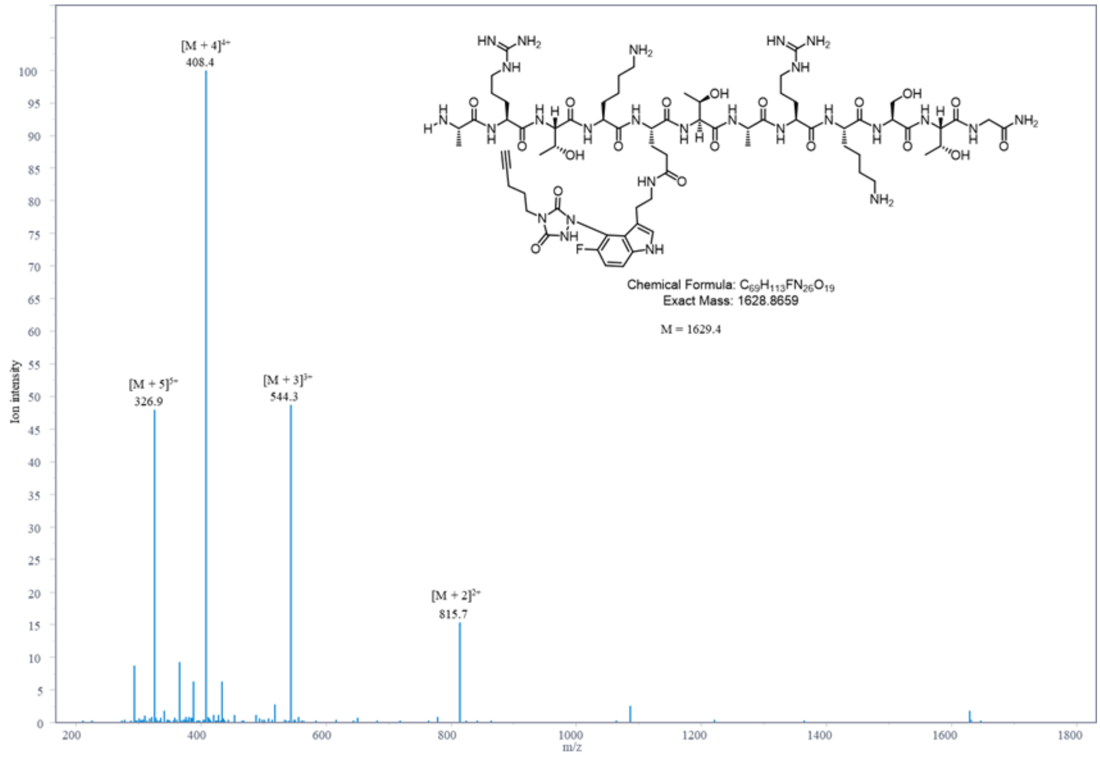

MS spectrums of ARTKQdaqTARKSTG

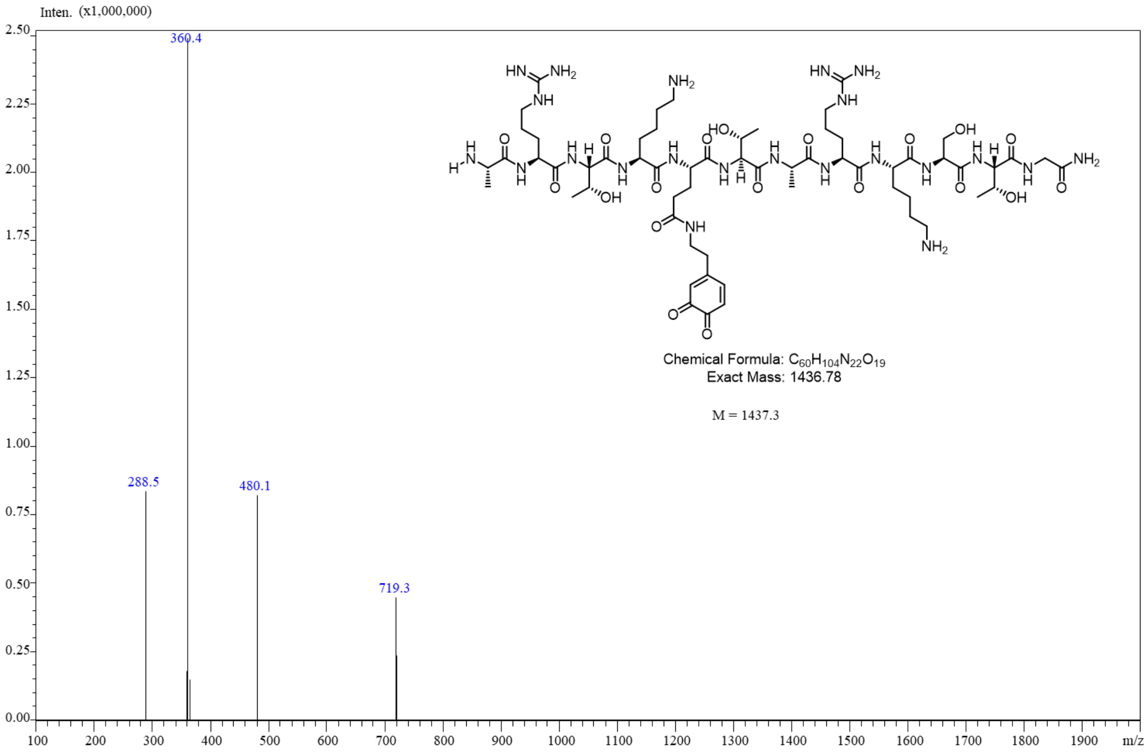

MS spectrum of AcNH-AWA-$CONH_2$

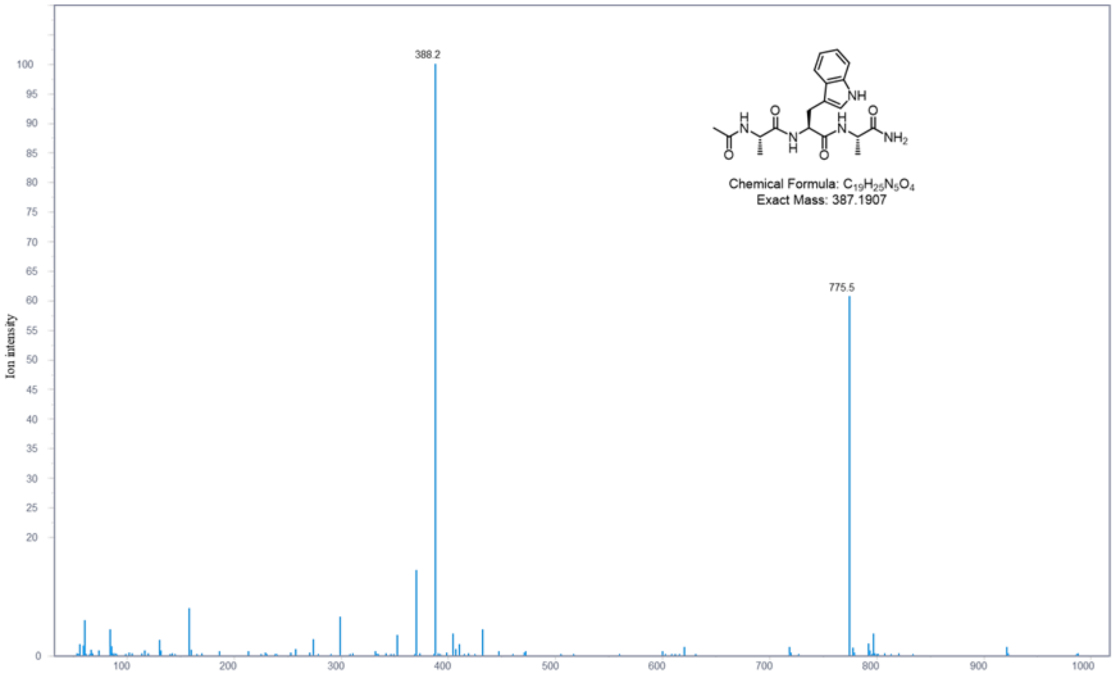

MS spectrum of AcNH-AWA-$CONH_2$ + AlkTAD

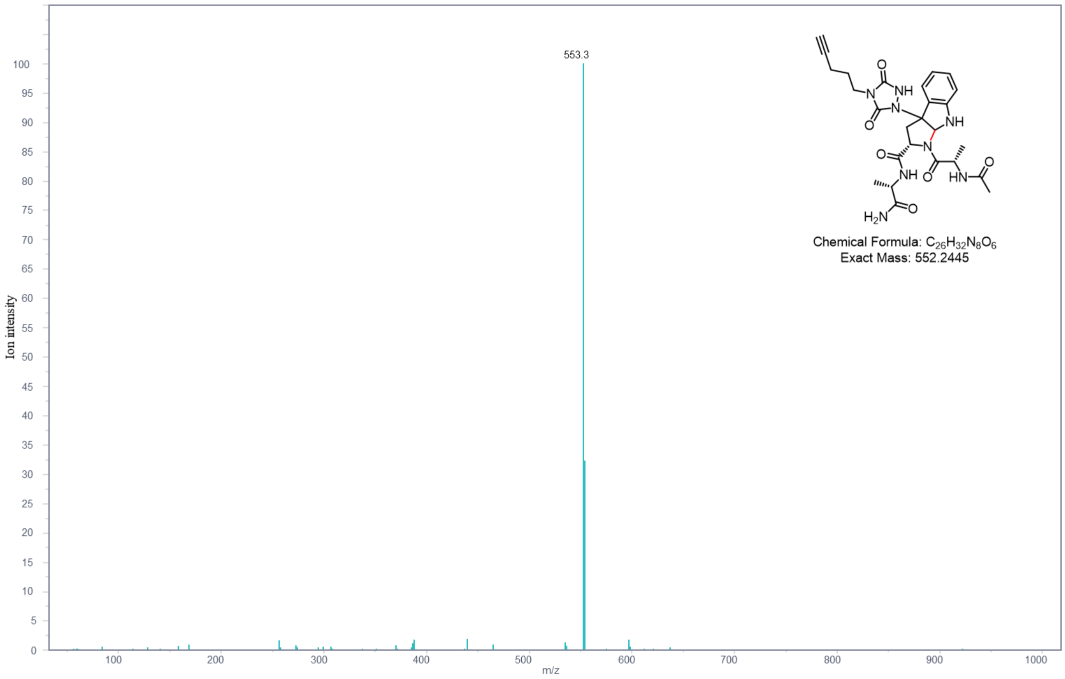

MS spectrum of AcNH-AWA-$CONH_2$ + AlkTAD **Component 1**

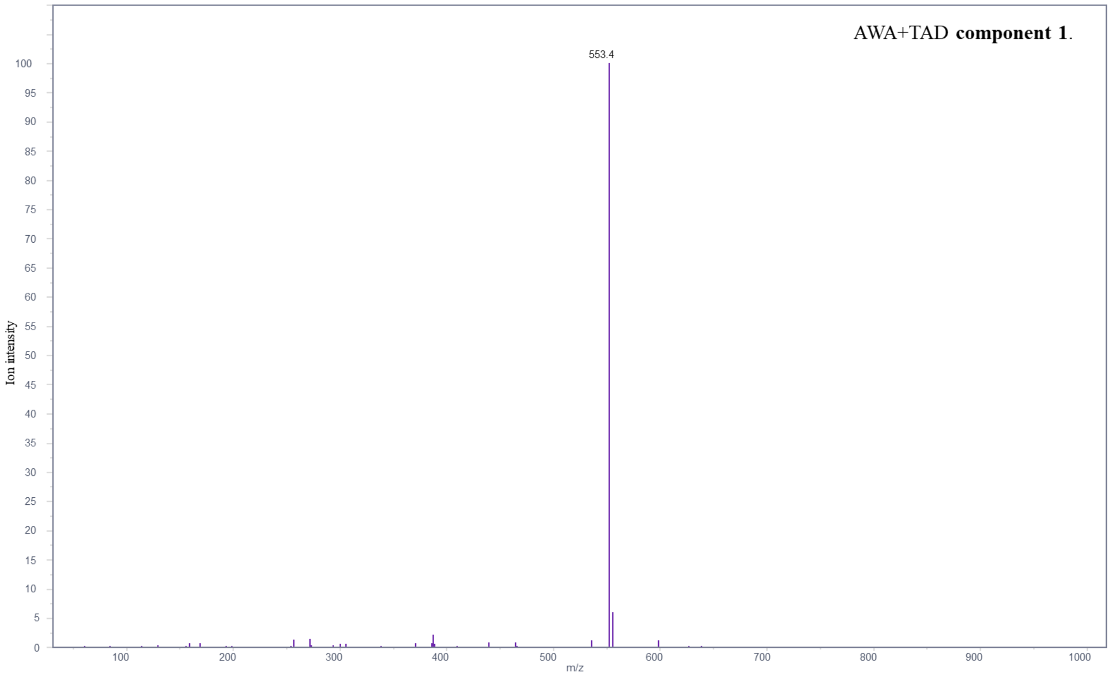

MS spectrum of AcNH-AQserA-CONH$_2$

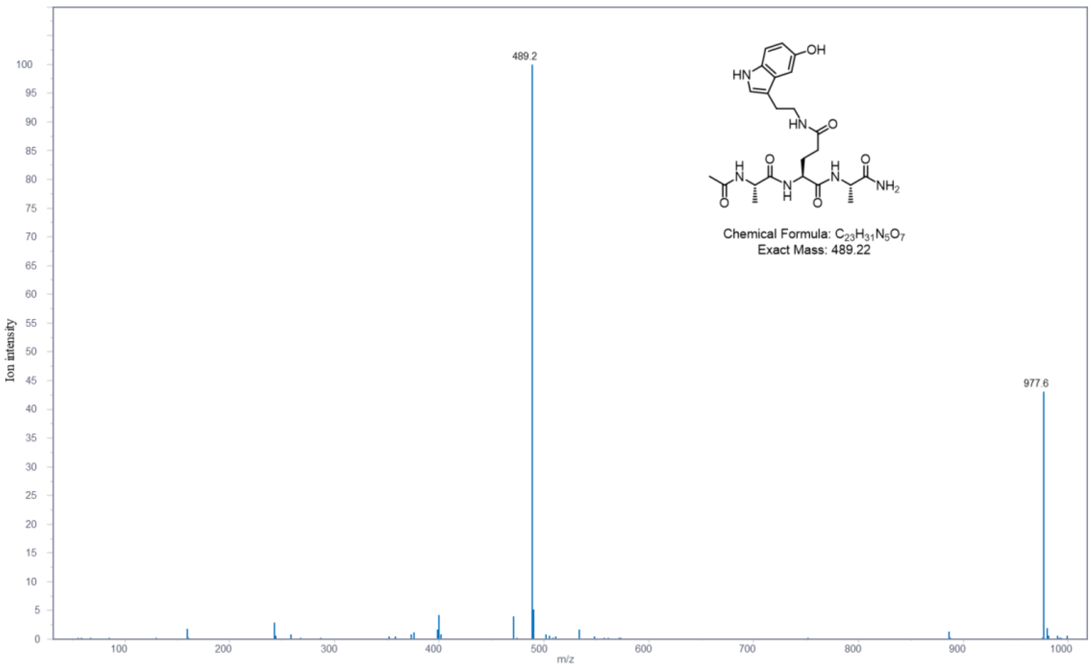

MS spectrum of AcNH-AQserA-CONH$_2$ + AlkTAD

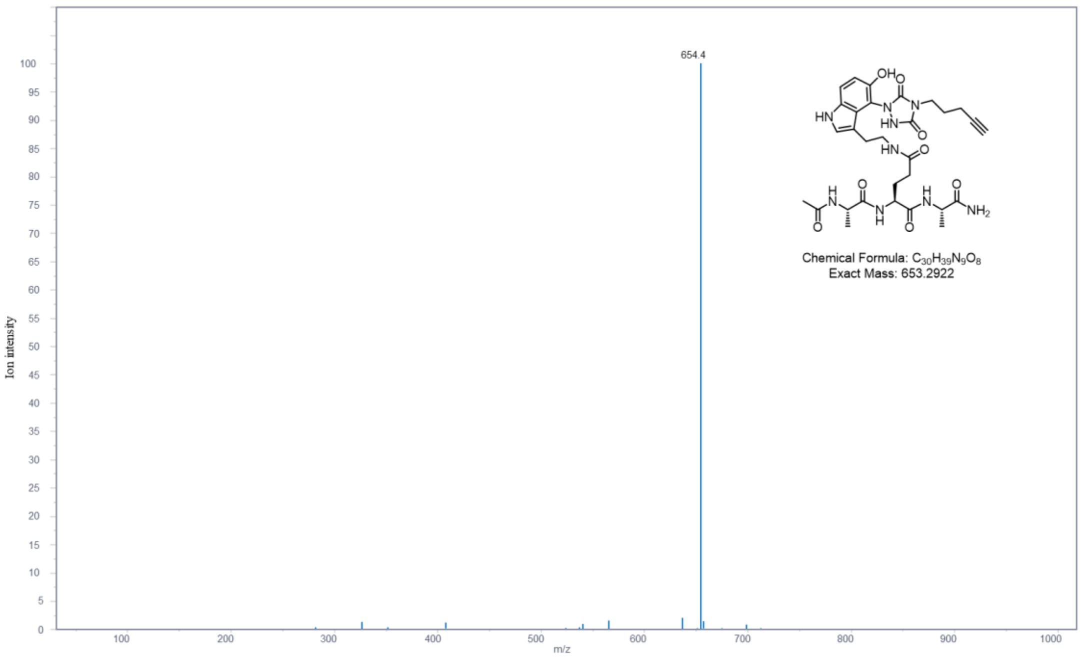

### 13. REFERENCES AND NOTES

1. Zheng, Q.; Weekley, B. H.; Vinson, D. A.; Zhao, S.; Bastle, R. M.; Thompson, R. E.; Stransky, S.; Ramakrishnan, A.; Cunningham, A. M.; Dutta, S.; Chan, J. C.; Di Salvo, G.; Chen, M.; Zhang, N.; Wu, J.; Fulton, S. L.; Kong, L.; Wang, H.; Zhang, B.; Vostal, L.; Upad, A.; Dierdorff, L.; Shen, L.; Molina, H.; Sidoli, S.; Muir, T. W.; Li, H.; David, Y.; Maze, I. Bidirectional histone monoaminylation dynamics regulate neural rhythmicity. *Nature* **2025**, *637*, 974-982.

2. Zhang, N.; Wu, J.; Hossain, F.; Peng, H.; Li, H.; Gibson, C.; Chen, M.; Zhang, H.; Gao, S.; Zheng, X.; Wang, Y.; Zhu, J.; Wang, J.J.; Maze, I.; Zheng, Q. Bioorthogonal labeling and enrichment ofhistone monoaminylation reveal its accumulation and regulatoryfunction in cancer cell chromatin. *J. Am. Chem. Soc.* **2024**, *146*, 16714-16720.

3. Houck, H. A.; Bruycker, K. D.; Barner-Kowollik, C.; Winne, J. M.; Prez, F. E. D. Tunable blocking agents for temperature-controlled triazolinedione-based cross-linking reactions. *Macromolecules* **2018**, *51*, 3156-3164.

4. Ray, D. M.; Jennings, E. Q.; Maksimovic, I.; Chai, X.; Galligan, J. J.; David, Y.; Zheng, Q. Chemical labeling and enrichment of histone glyoxal adducts. *ACS Chem. Biol.* **2022**, *17*, 4, 756-761.
